# Sculpting New Visual Categories into the Human Brain

**DOI:** 10.1101/2020.10.14.339853

**Authors:** Coraline Rinn Iordan, Victoria J. H. Ritvo, Kenneth A. Norman, Nicholas B. Turk-Browne, Jonathan D. Cohen

## Abstract

Learning requires changing the brain. This typically occurs through experience, study, or instruction. We report a proof-of-concept for a new way for humans to acquire visual knowledge by directly sculpting activity patterns in the human brain that mirror those expected to arise through learning. We used a non-invasive technique (closed-loop real-time functional magnetic resonance imaging neurofeedback) to create new categories of visual objects in the brain, without the participants’ explicit awareness. After neural sculpting, participants exhibited behavioral and neural biases for the sculpted, but not for the control categories. The ability to sculpt new perceptual distinctions in the human brain offers a new paradigm for human fMRI research that allows for non-invasive, causal testing of the link between neural representations and behavior. As such, beyond its current application to perception, our work potentially has broad relevance to other domains of cognition such as decision-making, memory, and motor control.

**Significance Statement:** Objects that belong to the same category tend to elicit similar patterns of brain activity. Here we reverse this mapping and ask whether neural similarity is sufficient to induce increased perceptual discrimination and categorical perception. We do this by using real-time fMRI to modify neural representations of objects in high-level visual cortex. Participants viewed an object and received closed-loop neurofeedback that pushed them to represent the object more similarly to a brain activity pattern we chose for that category. After successfully self-modulating their brain activity, participants began to perceive objects assigned to the same brain pattern as more categorically distinct from those assigned to a different brain pattern. These findings open a new avenue for understanding and accelerating human learning.

## Introduction

*“For if someone were to mold a horse [from clay], it would be reasonable for us on seeing this to say that this previously did not exist but now does exist.”*

Mnesarchus of Athens, ca. 100 BCE (*1*)

Humans continuously learn through experience, both implicitly (e.g., through statistical learning; *2,3*) and explicitly (e.g., through instruction; *4,5*). Brain imaging has provided insight into the neural correlates of acquiring new knowledge (*6*) and learning new skills (*7*). As humans learn to group distinct items into a novel category, neural patterns of activity for those items become more similar to one another and, simultaneously, more distinct from patterns of other categories (*8–10*). We hypothesized that we could leverage this process using neurofeedback to devise a fundamentally new way for humans to acquire perceptual knowledge. Specifically, sculpting patterns of activity in the human brain (’molding the neural clay’) that mirror those expected to arise through learning of new visual categories may lead to enhanced perception of the sculpted categories (’they now exist’), relative to similar, control categories that were not sculpted. To test this hypothesis, we implemented a closed-loop system for neurofeedback manipulation (*11–18*) using functional magnetic resonance imaging (fMRI) measurements recorded from the human brain in real time (every 2s) and used this method to create new neural categories for complex visual objects. Crucially, in contrast to prior neurofeedback studies that focused exclusively on reinforcing or suppressing existing neural representations (*11,12*), in the present work we sought to use neurofeedback to create novel categories of objects that previously did not exist in the brain; we test whether this process can be used to generate significant changes in the neural representations of complex stimuli in human cortex, and, as a result, alter perception.

## Results

### Constructing a Normed Stimulus Set of Complex Objects for Neural Sculpting

Using radial frequency components (RFCs) of an image (*19–21*), we generated a two-dimensional manifold of closed-contour shape stimuli that varied smoothly in appearance as a function of distance from a center shape (Fig. 1A). To obtain each shape, sine waves determined by the seven RFCs were added together and the resulting wave was wrapped around a circle to obtain a closed contour which was then filled in to create a shape. We chose this technique for building a complex visual stimulus space because prior work has shown that radial shapes generated using subsets of the same RFCs are perceived monotonically and that their neural representation is also monotonically related to parametric changes in the amplitude of the RFCs across multiple brain regions (*19,21*). We created a two-dimensional shape space by independently varying the amplitude of two of the seven RFCs (from 12.6 to 36.6 for the 4.94 Hz component and from −6.0 to +42.0 for the 1.11 Hz component), while holding the amplitudes of all other components constant. In this manifold, each of six equally spaced diameters (AG, BH, CI, DJ, EK, FL) defined a novel and arbitrary category boundary between the two groups of shapes on either side. We verified that the shapes were perceived similarly across categories in a psychophysical experiment with 10 participants using a self-paced two-alternative-forced-choice (2AFC) task. For each diameter through the space, we presented shapes along the diameter and participants judged which of the two “endpoint” shapes (e.g., A vs. G) more closely resembled the presented shape. We then used these categorization data to compute a psychometric function (Fig. 1B-G). Participants did not have *a priori* biases in how they categorized items across multiple partitions (psychometric function slope, repeated measures ANOVA, factor=‘direction’; F(5,54)=0.29, p=0.918).

**Fig. 1.**
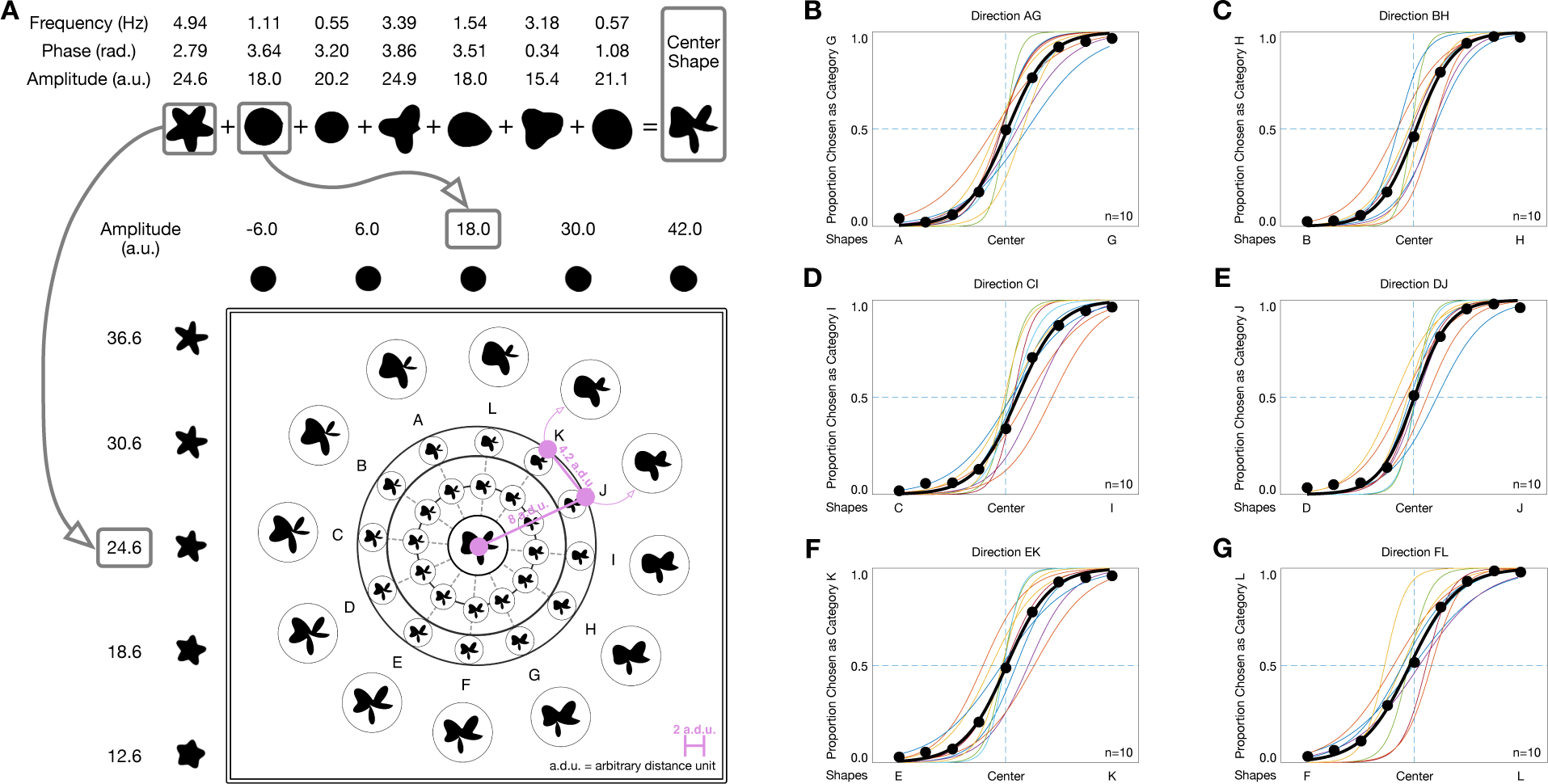
Shape space stimulus set construction and perceptual norming. (A) Stimuli were constructed using combinations of seven radial frequency components (RFCs) of fixed frequency F and phase P (*19,20*). We parametrically varied the amplitude A of two of the seven RFCs independently to generate a two-dimensional manifold of complex shapes: a displacement of 1 arbitrary distance unit (a.d.u.) on the manifold corresponded to an amplitude change of 6 arbitrary units for the 1.11 Hz component and an amplitude change of 3 arbitrary units for the 4.94 Hz component (changes in component appearance shown above and to the right of the shape space). Each point on this manifold corresponded to a unique shape that could be computed using the mathematical convention described above. Example shapes at 8 a.d.u. from the center are enlarged and shown on the outer edge (e.g., A, B, etc.); they lie 4.2 a.d.u. apart from each other on the manifold. (B-G) Psychometric function estimates for the 2AFC behavioral experiment conducted for each of six diameters: AG, BH, CI, DJ, EK, and FL. Thin lines in distinct colors represent psychometric functions for each participant (n=10) and thick black lines represent participant averages for each diameter.

Using this set of complex visual objects, we trained a new group of 10 participants over 10 days each in a real-time fMRI experiment. The participant sample size was planned during the experiment design and is comparable with (or exceeds) that of most other neurofeedback studies (*11–18*) that target primarily visual regions in human cortex. Notably, our 10 participants corresponded to 98 total sessions (20 behavioral, 78 in-scanner, 2h each), providing considerably denser sampling and a larger dataset than conventional fMRI or other similar fMRI neurofeedback studies. We used a closed-loop neurofeedback procedure to sculpt a categorical distinction between the objects on either side of one diameter through shape space. By sculpting divided neural representations for these objects, we sought to change how they are perceived (Fig. 2A). On Day 1, participants performed a 2AFC behavioral task (identical to the one used to norm the stimulus space) to obtain a baseline measure (psychometric function slope) of how they categorized stimuli along each diameter in the space. We verified that the training cohort of participants did not have any biases across categorization directions (replicating the norming cohort results) and we also verified that there were no significant differences between the training and norming cohorts (psychometric function slope, repeated measures ANOVA, factors=‘direction’, ‘cohort’; interaction: F(5,108)=0.19, p=0.964; main effect of direction: F(5,108)=0.57, p=0.722; main effect of cohort: F(1,108)=1.71, p=0.194) (Fig. S1). On Days 2-3, we ran two fMRI sessions in which we mapped how each individual participant’s brain represented the stimulus space, defined candidate brain regions for neurofeedback, and built a model of how the shape space was represented in participants’ brains to use for real-time tracking during the training phase of the experiment. After Day 3, we randomly selected one of the diameters as the category boundary to be sculpted during training for each participant (not disclosed to them). On Days 4-9, participants underwent real-time fMRI neurofeedback training, during which they were shown wobbling versions of the shapes (Movie S1) and asked to generate a mental state that would stabilize the shape in their visual display. Unbeknownst to the participants, they were given positive visual feedback (reduced wobbling) and monetary rewards when the neural representation of the shape they were viewing resembled the neural representations of other shapes in the category selected to be sculpted during training. Finally, on Day 10, participants repeated a version of the 2AFC behavioral task, in which we evaluated changes in categorical perception across both sculpted and control category boundaries.

**Fig. 2.**
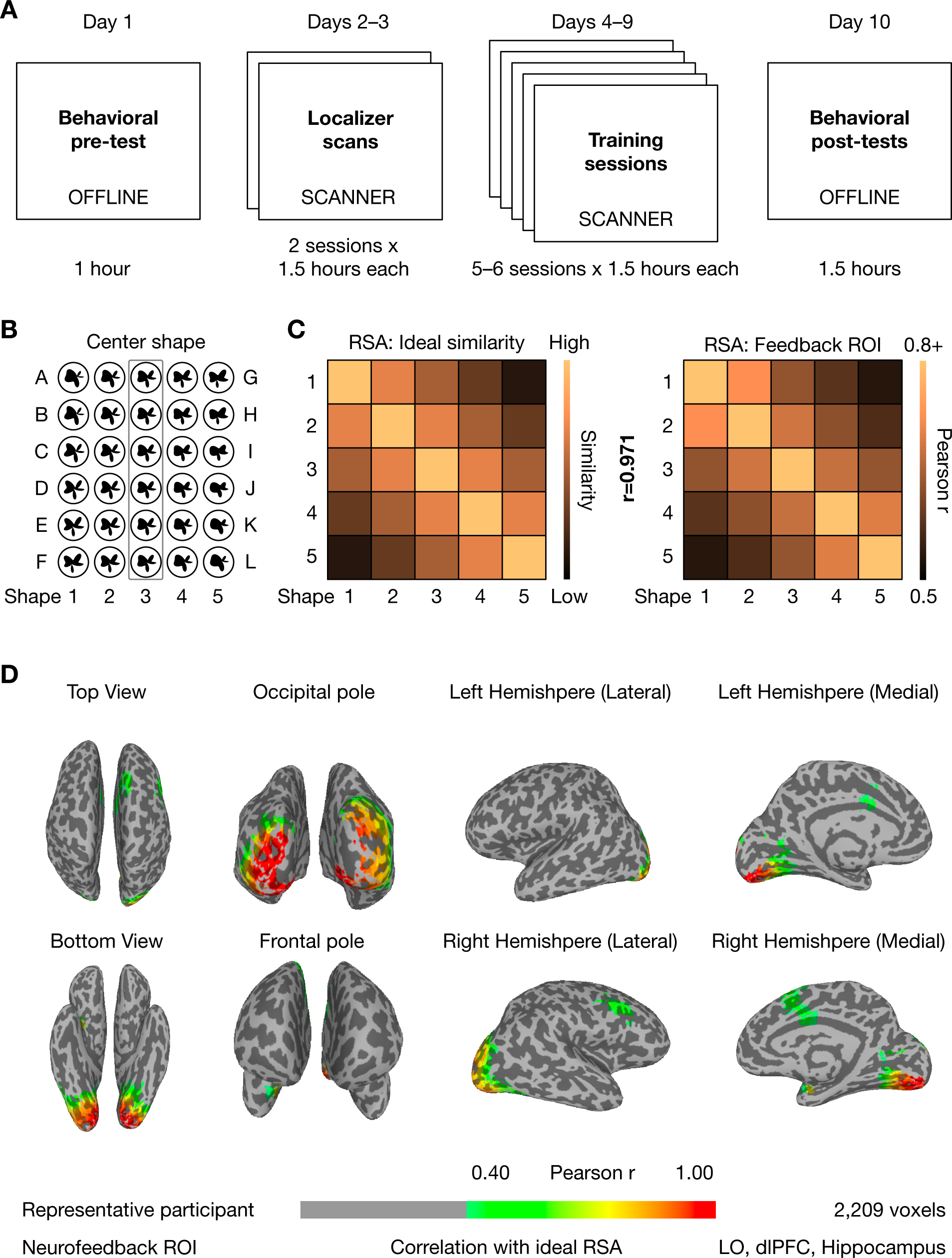
Experimental procedure and neurofeedback ROI computation. (A) The experiment extended over 9-10 days for each participant: one behavioral pre-test (2AFC along each diameter from Fig. 1), two localizer fMRI sessions, 5-6 real-time fMRI neurofeedback training sessions (5 days: n=2, 6 days: n=8), and a final behavioral post-test (repeat of first day). (B) 25 shapes shown during the localizer scans were used to define parametric neural representations of the shape space: 5 shapes along each diameter, −8, −4, 0, +4, +8 a.d.u. from the center shape common to all diameters. (C) Ideal representational similarity matrix (RSM) for a neural parametric representation (left) and average RSM for the 25 shapes in the final neurofeedback region of interest (ROI) of the training cohort (n=10, right). The correlation between the ideal and observed RSMs was very high (r=0.971) for the selected neurofeedback ROI across all participants, suggesting strong parametric representation of the shape space in these brain regions. (D) Cortical map of the neurofeedback ROI for an example participant encompassing several brain regions, including extrastriate visual cortex, parahippocampal gyrus, hippocampus, and medial frontal gyrus (see Methods for details of the selection procedure). Cortical maps for all participants (n=10) are shown in Figs. S3-S12.

### Using the Shape Space to Sculpt New Neural Categories of Objects in Multiple Brain Regions

We defined a region of interest (ROI) to target with neurofeedback by running two block-design fMRI localizer sessions (Days 2-3) in which we showed participants 12-15 repetitions of 49 shapes spanning the two-dimensional stimulus manifold. We constructed an ideal representational similarity matrix of a subset of 25 of these shapes (Fig. 2B) and performed a cortical searchlight analysis (*22*) to find all brain regions that represented the shape space parametrically (i.e., akin to a cognitive map (*23,24*), Fig. 2C,D). We defined the target neurofeedback ROI for each participant as the union of all their parametric representation voxels, excluding early visual cortex (since we were interested in high-level visual perception related to complex object categories, the pericalcarine region was defined anatomically using Freesurfer; (*25*)). The final neurofeedback ROIs comprised 780-2,209 voxels per participant (average 1,401±115 s.e.m. voxels; Figs. 2C & S2). A representative ROI is shown in Fig. 2D, all participants’ final ROIs are shown in Figs. S3—S12, the unions of all parametric voxels for each participant (including early visual cortex) are shown in Figs. S24—S33, and the group map for the average neurofeedback ROI across all participants after alignment to common MNI space is shown in Fig. S34. We hypothesized that if any portion of these ROIs (either individually or in concert) were causally related to the categorical perception of our stimuli, then sculpting novel neural categories across all of them collectively would maximize the chances of influencing participants’ perception after training.

Using this ROI and the fMRI data from Days 2-3, we selected an arbitrary diameter in the stimulus space as a category boundary for each participant by rolling a pink six-sided die (Fig. 3A). Participants were not informed that this was a study about visual categories, that the shapes were drawn from a continuous circular space bisected by diameters, nor about which diameter was randomly selected as the boundary. We built a model of the neural representations of the two resulting categories of shapes. Each category was modeled as a multivariate Gaussian distribution with shared covariance (Fig. 3B). The parameters of the category-specific distributions were computed using maximum-likelihood estimation for the top 100-200 principal components of the signal elicited by that category’s shapes in the neurofeedback ROI (we used a grid search to determine the optimal number of components and hemodynamic lag for each participant). To verify that our model could predict the distinction between shape categories in the brain, we inverted it into a discriminative log-likelihood-ratio-based pattern classifier. Leave-one-day-out decoding accuracy was high (chance = 50%): 73%-81% in the lateral occipital region (defined using Freesurfer; (*25*)), known to represent distinctions between closed contour objects (*26*), and 71%-79% in our neurofeedback ROI (Fig. 3C). Given that our ability to induce multivariate pattern-level neural plasticity using neurofeedback is predicated on the highly accurate decoding of shape category during training, this measure was also used as a selection criterion for inviting participants back for training after Day 3 (criterion: >70% decoding accuracy for both LO and neurofeedback ROI; 7 participants excluded; Table S1). This criterion also ensured that our participants rated highly on a strong predictor of the effectiveness of neurofeedback manipulation (neural signal strength as measured by baseline task-related decoding accuracy (*27*)) and, most importantly, that our neurofeedback models would have a high chance of providing accurate feedback to participants during the majority of the training trials, as our key outcomes depend on changes in multivariate representations that must be detectable during real-time engagement with the stimuli.

**Fig. 3.**
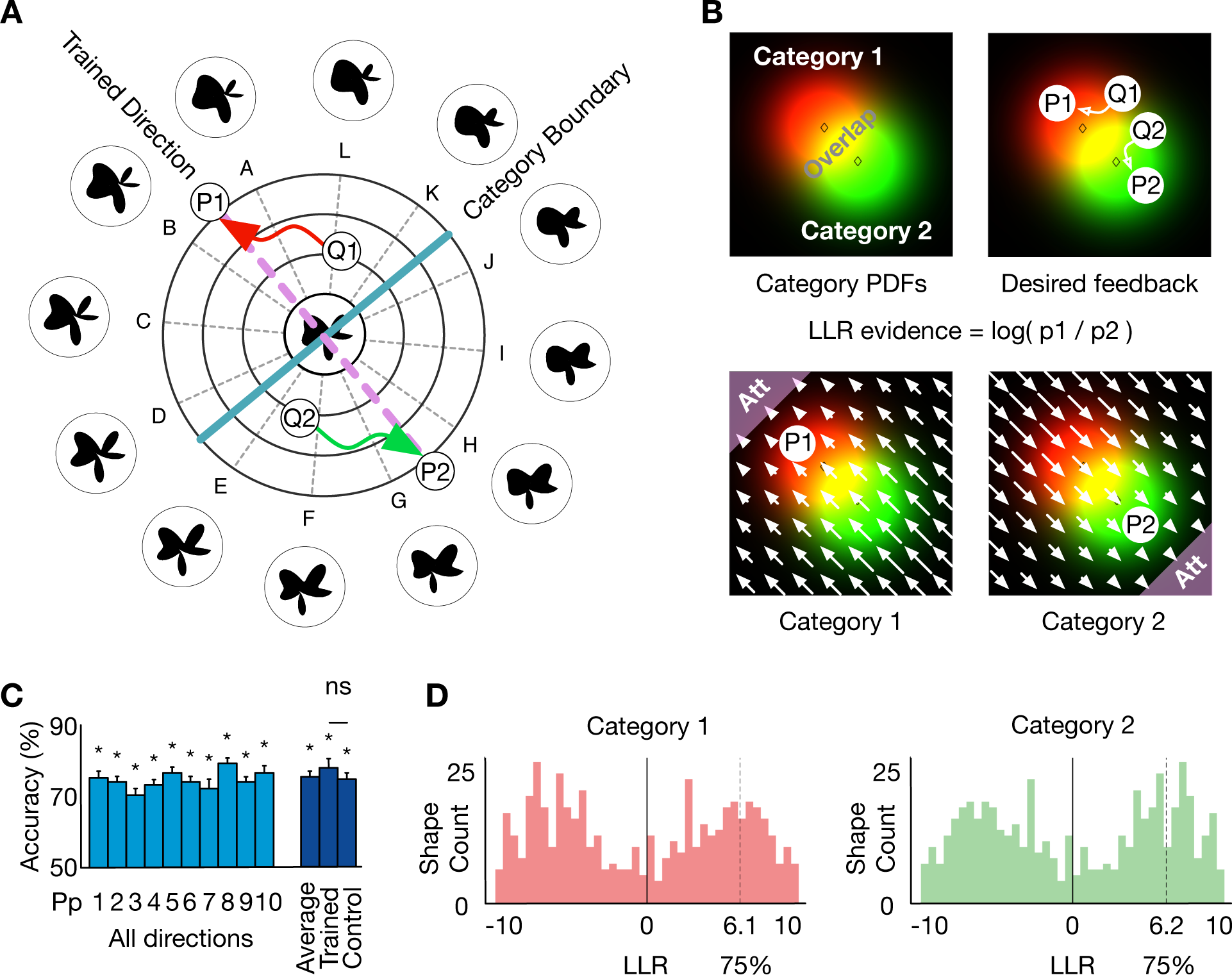
Using a neural model of the shape space to guide neurofeedback. (A) Unbeknownst to participants, we chose an arbitrary category boundary (blue) and sought to sculpt separate neural representations of shape categories along the perpendicular direction (purple). For example, the neural representations of shapes Q1 and Q2 were sculpted to be increasingly distant from the category boundary, illustrated by P1 and P2, respectively. (B) Two-dimensional schematic of neural representations of shape categories (the actual neural space comprised 100-200 dimensions). We sculpted the neural representations of shapes to become more similar to the corresponding attractor regions (Att) for categories 1 and 2. Arrows indicate relative change in log-likelihood ratio (LLR) required to reach the goal. (C) Gaussian model accurately predicted shape category in all participants during localizer scans (Days 2-3, leave-one-day-out decoding 75.7%±0.854% s.e.m.; chance=50%, *p<0.001). We observed no significant differences between decoding accuracy for the trained (sculpted) and control categories (trained: 78.1%±1.46% s.e.m.; control: 75.4%±1.38% s.e.m.; trained > control t(9)=1.12, p=0.293). The graph shows only participants who exceeded the 70% threshold and subsequently underwent training (also see Table S1). (D) Graphs show histogram of LLR values for neural representations elicited by all shapes presented during the localizer scans in the neurofeedback ROI for an example participant (LLR was computed using the cross-validated, optimal Gaussian model used for neurofeedback training). Feedback was given if neural evidence for the category of the shape on the screen (e.g., Q1) exceeded a given threshold in the distribution of LLR values for all shapes shown during the localizer scans. We note that the 75% threshold used for training may be different for each category under the same model. LLR histograms with 75% thresholds for all participants are shown in Fig. S13.

During neurofeedback training (Days 4-9), participants were told that during each trial they would see a continuously wobbling shape and had to “Generate a mental state that will make the shape wobble less or even stop!” Participants received additional instructions designed to accommodate for the hemodynamic lag inherent to the fMRI signal (e.g., that progress on the task is related to their mental state over the previous 8-10 seconds) and were incentivized with a monetary reward for each trial in which they made progress. However, participants were never told that a category boundary existed in the shape space or that feedback was based on the shape of the object, no less its relationship to the selected category boundary (see Methods for details). For each neurofeedback trial, we selected a random shape (seed) drawn from one of the two categories and generated a continuous oscillation in parameter space centered at that shape’s coordinates. Visually, participants saw shapes in the center of the screen (center of mass remained stationary) that morphed smoothly and gradually between the seed shape and other nearby, similar shapes (Movie S1). The apparent magnitude of the shape morph on the screen was manipulated via neurofeedback at every fMRI timepoint (TR=2s). For each individual participant, we computed the distribution of LLR values under our estimated Gaussian distributions for all shapes shown their localizer scans (520—800 trials, Fig. 3D) and positive feedback (less oscillation) was given during training if the neural evidence for the category of the current shape exceeded a given threshold in this distribution (Figs. 3D, S13, & S14, Movie S2). The initial threshold was selected based on pilot scans indicating that the neural model’s decoding performance decayed as a function of wobble intensity (Fig. S15). The threshold was continually adjusted during the experiment using an adaptive procedure designed to provide feedback on approximately 33% of the trials (Table S2). Each participant (n=10) completed 520–800 trials of closed-loop real-time neurofeedback training over the course of 5-6 daily sessions. A summary of feedback thresholds and performance for each participant, training session, and run is given Table S3.

### Neural Sculpting Generated New Visual Categories in the Brain and in Perception

After finishing the training sessions, we evaluated whether neural sculpting of the novel visual categories was successful; that is, whether the closed-loop real-time neurofeedback procedure induced multivariate pattern-level neural plasticity of the shape space representation. To measure changes in the neural representations of shape categories, we computed the difference between the average LLR of each trial (the log-likelihood ratio of the neural representation elicited by each shape in the neurofeedback ROI under the Gaussian model trained to differentiate the two trained categories of shapes) during the first two days of training versus the last two days of training (since participants underwent 5 or 6 days of training, there was no clearly defined first/second half of the experiment for everyone). We compared this score between the “trained” shape categories that we expected to have separated in the brain because they were bisected by the selected diameter, versus the “control” shape categories that would have been created by the perpendicular diameter, for which no separation was expected (Fig. 4A). We found strong positive neural sculpting effects for 6 out of 10 participants (between +0.342 and +1.204) and weak negative effects for 4 out of 10 participants (between −0.212 and −0.067) (Fig. 4D). From the first two days to last two days of training, we found a reliable difference in the LLR change between trained and control categories (Fig. 4B-D; +0.369±0.154 s.e.m; t(9)=2.27, p=0.049, Cohen’s d = 1.421). Within this interaction, LLR showed an increase for the trained categories (Fig. 4B; +0.234±0.077 s.e.m., t(9)=2.87, p=0.019) and a numeric, but not significant decrease for the control categories (Fig. 4B; −0.136±0.087 s.e.m., t(9)=1.48, p=0.174). Effects of neural sculpting on neural representations for each individual participant are shown in Fig. S16A. These results were obtained using the LLR for the first timepoint of each neurofeedback trial to avoid potential carry-over fMRI-adaptation effects for successively presented stimuli (*28,29*): since the shape stimuli oscillate around a fixed average shape throughout an entire trial, regardless of feedback-controlled amplitude, the activity level in brain regions that are involved in processing the stimulus (e.g., the neurofeedback ROI) may decrease as the trial progresses. Thus, the strongest measurement of the change in the neural representation of a stimulus is between the beginning of a trial at the start of neurofeedback training and the beginning of a trial at the end of neurofeedback training. This prediction is consistent with the results reported in Table S4: the neural effect of training is strongest for the first timepoint (TR) and remain high and significant but decrease slightly with each subsequent TR (up to a maximum possible of five) where adaptation may affect the neural representation in the neurofeedback ROI. Similar results were also observed using a ratio of the LLR for trained and control categories, instead of a difference score (Fig. S17). Together, these results suggest that our real-time fMRI neurofeedback neural sculpting procedure was successful in manipulating how the human brain represents complex objects from the multidimensional shape space we constructed.

**Fig. 4.**
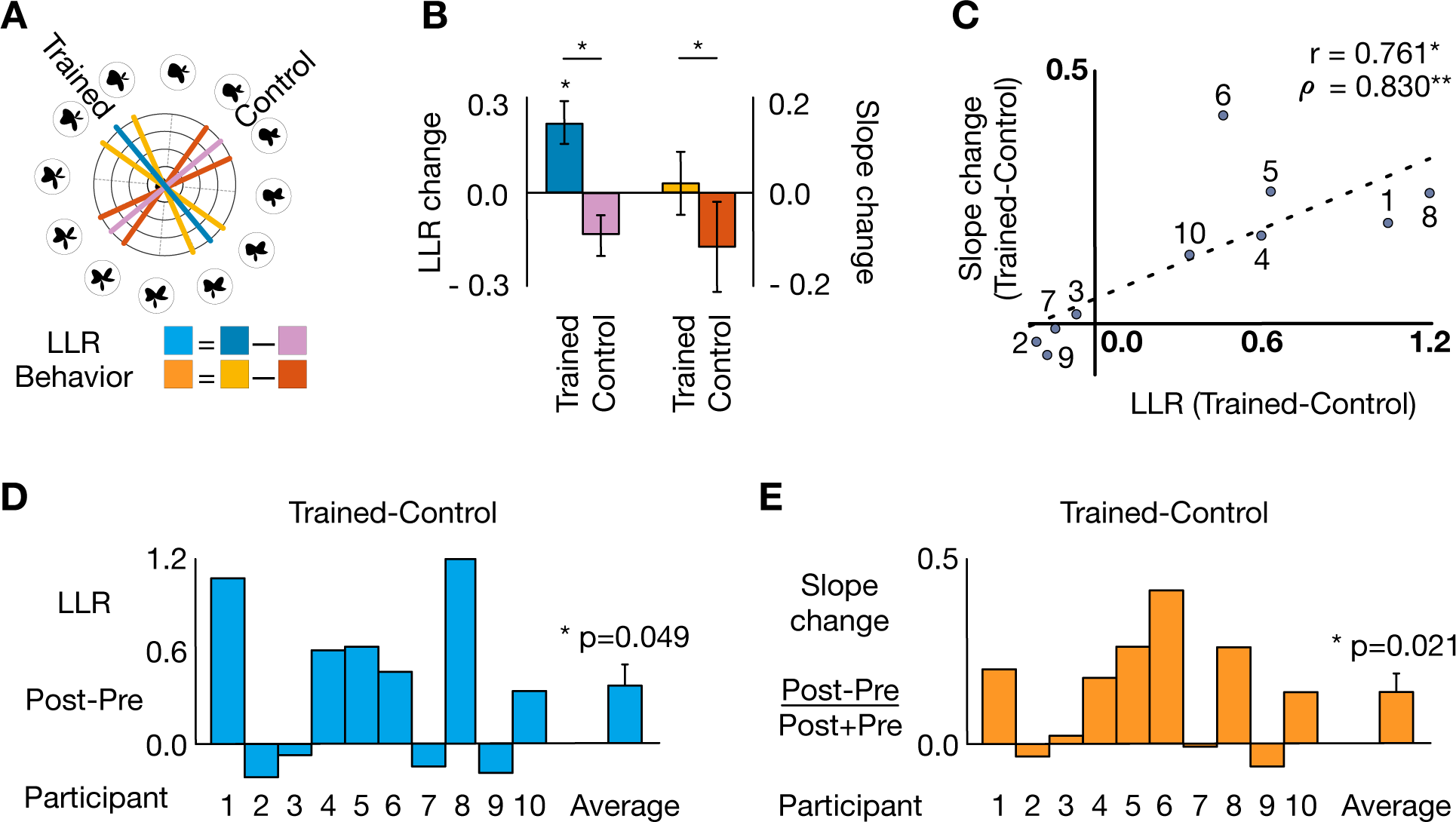
Neurofeedback successfully sculpted new visual categories in the human brain and altered perception. (A) Shape space with example category boundaries. Diameters for trained category distinction: LLR = blue, psychometric function slope = average of yellow lines. Diameters for control category distinction: LLR = purple, psychometric function slope = average of red lines. (B) Effects of neural sculpting on neural representations and perception for trained and control categories: differences in LLR and psychometric function slopes (colors as in subpanel A; *p<0.05). (C) Changes in the brain due to neural sculpting predict perceptual changes (Pearson r, *p=0.011; Spearman rho, **p=0.006). (D) Change in LLR between the last two days and the first two days of training for individual participants. Positive values indicate stronger neural boundaries in trained vs. control categories. (E) Change in psychometric function slope for trained vs. control categories between behavioral pre- and post-tests for individual participants. Positive values indicate stronger categorical perception for trained versus control categories.

Our goal was not only to modify the brain but also to test the hypothesis that neural sculpting is sufficient to alter behavior. Namely, we predicted that training would induce categorical perception whereby shapes close to the category boundary come to be perceived as clearer category members. Thus, we hypothesized that the slope of the psychometric function running perpendicular to the sculpted boundary would become steeper, compared to the psychometric function running alongside the sculpted boundary. To measure these perceptual changes, we estimated psychometric functions for the trained and control categories from the 2AFC behavioral task conducted on Day 10. Most participants reported fatigue with the experiment (after 5-6 consecutive days of fMRI scans) and we observed a tendency for some participants to perform worse at categorization across all directions in the post-test compared to the pre-test (Table S5 & Fig. S16B), as well as higher variability overall for psychometric function slopes across all directions in the post-test compared to the pre-test (average slope pre: 1.85±0.071 s.e.m.; average slope post: 1.98±0.339 s.e.m.; Table S5 & Fig. S23). We predicted that neural sculpting was successful at altering behavior for all participants, even if fatigue, decreased motivation, and/or disengaged reporting strategies during the post-test (*30*) potentially decreased the slope of psychometric functions across all directions for some participants. As such, we would observe a perceptual advantage for the trained direction after training, irrespective of overall categorization performance; that is, psychometric function slopes for all participants would relatively increase for the trained direction compared to the control direction, between the pre- and post-tests. To measure the relative change for the trained and control directions in the pre- and post-test measures across all participants, we calculated a normalized difference score between the psychometric function slopes from Day 10 and those on Day 1 (Fig. 4A). The discrimination slope significantly increased for the trained categories compared to the control categories (Fig. 4B,E; +0.137±0.049 s.e.m., t(9)=2.79, p=0.021, Cohen’s d = 0.505). We observed strong positive behavioral effects for 6 out of 10 participants (0.137-0.414), a weak positive effect for one participant (0.021) and weak negative effects for 3 out of 10 participants (−0.062, −0.033, −0.008). Estimated psychometric functions for all tests, directions, and participants are shown in Fig. S16B & Table S5. Overall, across our entire cohort, positive effects of neurofeedback manipulation (both behavioral and neural) tended to be substantial, whereas negative effects were exclusively negligible. Bootstrap resampling analyses (10,000 samples) showed that our results are extremely robust at the individual level (Fig. S21; neural: p=0.005, behavioral: p<0.001; correlation p=0.006).

In light of this evidence, we interpret this induced categorical perception change as possibly resulting from the neural sculpting of the categories. Accordingly, the increase in behavioral discrimination slope should be related to the increase in neural separation of the category representations. Indeed, across participants there was a significant positive relationship between training-related behavioral and neural changes: the relative increase in LLR between neural representations of shapes from distinct categories in the neurofeedback ROI was correlated with the relative strengthening of perceptual categorization in the trained direction compared to the control direction (Fig. 4C; Pearson r=0.761, p=0.011; Spearman rho=0.830, p=0.006); a similar result was obtained using the LLR ratio measure (Fig. S17D). We also verified that null distributions for the Pearson and Spearman correlation values for our main effect had means close to zero (Pearson r mean<0.001, Spearman rho mean=0.001) and that the correlation values observed for our main effect were highly significant compared to these null distributions (Pearson p=0.006; Spearman p=0.002; Fig. S35). Additionally, a linear mixed effects model analysis showed that behavioral changes in our cohort were significantly predicted by neural changes induced by neurofeedback, even when accounting for the effects of number of training days, baseline decoding accuracy during localizer scans, and the random category split chosen for each participant during training (p<0.001).

The success of our neural sculpting procedure may be related to participant age, which showed a trending correlation with the strength of the observed behavioral effects (r=0.600, p=0.067). One potential explanation for this result is that participant conscientiousness or their level of engagement with the experimental task (for both of which age may be a proxy) may play a critical role in the effectiveness of the neurofeedback manipulation. Additionally, we observed that individual variation in the ability to decode category information from neural activity patterns may also be a good predictor of the eventual success of our neural and behavioral manipulation procedure. Specifically, category decoding was the basis of neurofeedback and thus better performance would lead to more precise feedback. To test this relationship, we defined a baseline neural decoding measure for each participant as the model decoding accuracy in the neurofeedback ROI during the two localizer scans (Days 2-3). As expected, baseline decoding was high for all participants (75.7%±0.854% s.e.m., chance=50%, Fig. 3C), and no significant difference was observed at baseline between trained (78.1%±1.46% s.e.m.) and control (75.3%±0.875% s.e.m.) categories (t(9)=1.12, p=0.293). However, collapsing across categories, baseline decoding was highly correlated with the neural LLR change (r=0.720, p=0.019) and the behavioral slope change (r=0.777, p=0.008) from pre- to post-training. Interestingly, these correlations persisted (LLR, r=0.710, p=0.021; slope, r=0.729, p=0.017) even after excluding the trained direction from the baseline decoding average, suggesting that the precision of shape representations (or our ability to recover them) can be considered as a general property of individuals that can be used to predict training success. Moreover, these findings provide strong post-hoc support for our decision to focus neurofeedback training on participants with high baseline neural decoding.

To measure whether lapses in performance between the behavioral pre- (Day 1) and post-tests (Day 10) could potentially explain the effects of neurofeedback training on perception, we performed a separate joint estimate of the lapses (upper and lower), thresholds, and slopes for our psychometric functions, before and after training (parameter estimates: Table S6; psychometric function estimates: Fig. S36). The separate estimates of the slope change replicated our original results: the difference in slope between the trained and control directions increased significantly as a consequence of training (slope change = 0.122, t(9) = 2.678, p = 0.025) and the updated slope change was also significantly correlated with the LLR change induced via neurofeedback in our ROI, using both parametric and non-parametric tests (Pearson r = 0.725, p = 0.018; Spearman rho = 0.746, p = 0.018). Additionally, psychometric function thresholds and lapses did not change significantly as a consequence of neurofeedback training (thresholds p=0.941, upper lapses p=0.472, lower lapses p=0.989), and a linear mixed effects model showed that the effect of neurofeedback on the new slope change difference remained significant after controlling for threshold and lapses: t(18) = 2.403, p = 0.027 (Fig. S37).

Our results cannot readily be explained by differences in the amount of feedback received by each participant, either overall, or for each individual category. Because of our adaptive feedback procedure, each participant received a similar amount of feedback as a proportion of total experiment trials (28.7%±0.657% s.e.m.) and for each stimulus category (Category 1: 14.9%±1.74% s.e.m.; Category 2: 13.9%±1.43% s.e.m.; t(9)=0.305, p=0.767). Additionally, the distribution of feedback values received by participants was not different between Trained and Control category boundaries (t(9)=0.82, p=0.435; Kolmogorov-Smirnov test=0.50, p=0.111) and an ideal-observer analysis showed that, across our cohort, the difference in feedback received for the Trained and Control categories was not sufficient to predict the participants’ neural (r=-0.447, p=0.196) or behavioral changes (r=0.320, p=0.283) after training, suggesting that the distribution of feedback values during training does not predict neural sculpting effects or participant performance.

Furthermore, our results cannot be explained by participants explicitly learning the random category distinction enforced during the training. At the conclusion of the study (but before being fully debriefed), we informed participants that the stimuli came from an unspecified number of categories. When asked to guess freely, participants reported an average of 3.3 categories whose boundaries did not coincide with those of the categories randomly selected for each of them in the experiment (Fig. S18 & Table S7). Only when subsequently forced to split the stimulus space into 2 categories using a straight line, their performance suggested a trend toward having some crude information about the category boundary (Figs. S18, S19 & Table S7; trained=0°, average absolute displacement=31.5°±5.89° s.e.m; control=90°, average absolute displacement=58.5°±5.89° s.e.m.; random choice=45°, t(9)=2.18, p=0.058). However, a linear mixed effect model confirmed that the effect of neurofeedback training on perception remains significant after controlling for angle displacement from the correct boundary, as well as for participant age (t(18)=4.290, p<0.001). Additionally, any such information about categories cannot readily be explained by implicit learning or priming (*31,32*) or by any potential serial dependence between the stimuli (*33*), given that the training included an equal number of stimuli from both categories, these stimuli were presented in a random order during training, and there were no differences in the amount of feedback received for stimuli of each category. Additionally, a support vector machine-based ideal observer analysis also confirmed that the feedback values received during training were not sufficient to predict shape category (decoding accuracy = 54.2%±3.51% s.e.m., chance = 50%; accuracy > chance, t(9)=1.14, p=0.283). This further suggests that any (small) amount of information about the category boundary that may have been inferable by participants due to the feedback itself was unlikely to have significantly influenced their ability to (not be able to) discover the category boundary during the experiment.

## Discussion

Neural sculpting with real-time fMRI puts forward two key advances showcased by our proof-of-concept study involving novel visual category learning. First, prior correlational studies have linked distributed patterns or fMRI activity to the perception of objects from different categories (*34–36*). By sculpting new neural representations in the human brain that mimic those that emerge as a consequence category learning and inducing corresponding perceptual discrimination changes, we provide causal evidence that these representations are sufficient for categorical perception. Second, prior neurofeedback studies focused exclusively on reinforcing existing neural representations of visual features or categories (*11,12*). In contrast, here we use neurofeedback for the more radical goal of sculpting categories that did not previously exist in the brain. In doing so, we demonstrate a direct causal link between multivariate pattern representations in the human brain and perception of complex visual objects that is accessible and amenable to manipulation using non-invasive functional neuroimaging. Together, our findings broaden the possibility for non-invasive causal intervention in humans with neurofeedback fMRI, including the distant possibility of sculpting more extensive knowledge and/or complex categories or concepts in the human brain, bypassing experience and instruction.

Previous studies have shown that learning novel categories induces increased within-category similarity (*9,10*), as well as neural and perceptual suppression of task-irrelevant features (*37–40*). Here, we used neural sculpting to induce a change in participants’ neural representations of the shapes that was intended to mimic this learning effect. Specifically, we linked exemplars in our shape space to one of two different neural representations via neurofeedback, resulting in increased neural and behavioral discrimination across the classification boundary separating these target representations relative to other irrelevant boundaries matched in stimulus space. As such, our findings show that neural similarity is sufficient to produce categorical perception, a major advance over prior work that established categories by linking exemplars to a common behavioral response (*8–10,37,39,40*). Consistent with these findings, the observed neural differences between trained and control categories reflected significant increases in sensitivity for trained categories and numeric, but not significant decreases in sensitivity for control categories (LLR, Figs. 4B & S16). A numerical, but not significant, effect in this direction was also observed in the perceptual measure (psychometric function slope, Figs. 4B & S16); the discrepancy between behavioral and neural results may be, in part, explained by reported participant fatigue, decreased motivation, and/or may reflect disengaged reporting strategies (*30*). Additional work is required to disambiguate the contributions of the two opposite effects of feature enhancement and suppression and their interaction with perceptual change within the context of neural sculpting, including how they influence the discriminability of these features by the participants.

A key strength of real-time neurofeedback is the ability to identify which brain regions are, and are not, required to induce a human behavior (*11*). Here, we sculpted visual categories within a feedback ROI that comprised multiple disparate brain regions, tailored to the individual response of each participant during independent localizer scans. We demonstrate that these regions, encompassing areas of inferior temporal and lateral occipital cortices involved in the visual perception of objects and categories (*34,36*), as well as areas of prefrontal cortex involved in category judgments, are sufficient to induce a new visual category. By the same token, the exclusion of early visual cortex from the feedback ROI suggests that these areas are not necessary to induce a new category. Our results leave open the question of which individual brain region(s) within the feedback ROI are necessary and/or sufficient. The current experiment was not designed to fully answer this question, but rather as a proof-of-concept that some brain regions exist whose manipulation via neurofeedback is not only possible, but also sufficient for influencing categorical perception. Nevertheless, several post-hoc analyses investigating pre- and post-training category decoding (Fig. S22) suggest that more posterior (e.g., posterior LO) and/or more visually selective brain regions (e.g., the occipito-temporal part of the training ROI) are potentially undergoing the most change due to neurofeedback training. As such, it is possible that these regions outside of early visual cortex might be driving the categorization effects we observe in behavior, similarly to how induced changes in V1 drove perceptual effects in prior work (*11*). However, more work is needed to fully understand where the bulk of the changes are happening in the brain due to training. Similarly, learning new visual categories can have lasting effects in the brain and in behavior, ranging from at least a few days (*41*) to several months or more (*42*). Although our study provides a proof-of-concept for visual category learning via neurofeedback, the durability of this neural sculpting remains an important question for future work, especially for determining the utility of this method for clinical applications.

We found that participants who showed higher baseline levels of neural shape decoding also showed larger training effects, suggesting that differences in either the precision of the neurofeedback signal or the quality of the underlying shape space representation may affect training outcomes. Similar to prior work (*12,27,43*), we observed variable training outcomes across our cohort, with some participants being more susceptible to neurofeedback manipulation compared to others. This may be due to individual differences in factors such as attentional control (*12*), differences in the ability to access and modify neural activity under an external constraint (i.e., non-responders; *43*), or differential plasticity across brain regions (*44*). In our cohort, the only three participants who exhibited a decrease in category separation for the trained direction after neurofeedback manipulation also exhibited overall neural and behavioral effects close to zero. Conversely, the only other participant who exhibited a decrease in discrimination for the trained direction after training, also exhibited an even larger decrease in discrimination for the control direction, coupled with a strong positive neural effect of training. We interpret this result as suggesting that successful neural changes induced for the trained direction for this particular participant improved relative discriminability for the trained direction, but that fatigue and/or disengaged reporting strategies (*30*) may have played an outsized role during their post-test behavioral session (Day 10 of the study). Taken together, these results provide additional evidence that behavioral effects are extremely robust at the individual level and that they hinge on successful neural manipulation.

Given the open-ended nature of our instructions for the neurofeedback trials (‘*Generate a mental state that will make the shape wobble less*!’), it is also possible that differences in strategies employed by the participants may have played a role in achieving success. Consistent with this, participants reported extremely varied approaches to the task, the most common of which involved naming the shapes and/or focusing on local features (e.g., an indentation) instead of the entire shape. Although no consensus strategy could be identified, nor one that could explain either the neural or behavioral outcomes (Table S7), it is possible that further work could identify such strategies.

Our study represents a proof-of-concept that neural sculpting can change neural patterns in high-level visual cortex and also induce behavioral change in participants’ perception. To maximize the ability of our neurofeedback neural models to provide consistent high-quality, reliable feedback to participants throughout the training portion of the experiment, we employed a relatively stringent criterion (70% decoding accuracy for shape categories in LO and in the neurofeedback ROI) for selecting participants that would undergo the second half of the study. This resulted in 7 out of 17 participants not being invited back for the full study after the two localizer scans. Future work is needed to elucidate the relationship between this decoding accuracy criterion (>70%) and the speed and efficacy of the neurofeedback manipulation induced by neural sculpting in the human brain and, correspondingly, in behavior. For example, subsequent studies could forego or manipulate this threshold and ascertain how the initial baseline decoding accuracy influences the strength of the neurofeedback effect. Similarly, since the speed with which neural changes could be induced in human cortex as a result of neural sculpting remains unclear, we designed our study to include at least 5 fMRI neurofeedback sessions across separate days. While both of these features of our study may potentially increase the difficulty of using our method going forward, they also provide evidence that with strong neural signal and decoding accuracy, as well as a sufficient amount of neurofeedback training, our type of manipulation can indeed change neural patterns in high-level visual cortex and also induce behavioral change in participants’ perception. Our study was meant to provide crucial evidence of what can be achieved with current technology in the field of neural and behavioral manipulation via neurofeedback, and we strongly believe that expected advances in signal acquisition (e.g., improved multiband protocols) will help both significantly increase base levels of neural decoding across the population, and reduce the training timeline requirements for similar studies in the future.

Our findings suggest that neural sculpting may also be an effective tool for influencing and potentially enhancing the neural effects of other learning processes that involve neural differentiation. For example, in the visual domain, our technique may prove useful for enhancing neural differences associated with domain-level expertise for categorization or recognition (e.g., in the fusiform face area, FFA; (*44,45*)), with visual memory co-activation in the hippocampus (*46*), or with brain-based education initiatives that complement classical learning paradigms (*47*). Conversely, our method may provide novel avenues to reverse de-differentiation of neural patterns stemming from aging (*48*) or from disorders impairing the natural function of visual brain regions such as visual agnosia (*49*) or prosopagnosia (*50*). Beyond the visual domain, previous studies have shown that other forms of neurofeedback aimed at enhancing or suppressing activity or connectivity of regions of interest (*11,51*) can be used to treat various neuropsychiatric disorders, such as major depressive disorder (*52–56*) and autism spectrum disorder (*57*). Our work provides a promising avenue towards potentially enacting more complex interventions in patient populations that would attempt to sculpt specific patterns of brain activity within regions of interest, in order to neurally mimic (or increase alignment with) neural activity patterns of healthy controls. Similarly, our work invites possible future applications for novel neurorehabilitation approaches, including brain-machine interfaces (*58*) and neuroprosthetics (*59*) that rely on generating or maintaining a specific multivariate pattern of brain activity in real-time.

To conclude, the work presented here represents a proof-of-concept for a new, non-invasive approach to investigating the causal relationship between neural representations and behavior using fMRI. More than 2,100 years after Mnesarchus of Athens made his observations about the nature of human experience (*1*), we show that his philosophical insights may, in fact, adequately describe how perception arises from human neural representations: sculpt a concept in the neural clay of the brain and it may subsequently exist.

## Acknowledgments and Funding Sources

The authors would like to thank M. L. Nguyen, A. N. Hoskin, E. A. Piazza, N. Keshavarzian, E. A. McDevitt, J. W. Antony, N. Y. Kim, and A. R. Burton for help with data collection; A. C. Mennen for discussions about the infrastructure for real-time fMRI neurofeedback; N. DePinto, L. E. Nystrom, and G. McGrath for technical support; and A. Tompary for contributions to initial pilot studies. Funding: John Templeton Foundation, Intel Corporation, NIH Award R01 MH069456.

## Materials and Methods

### Building and Norming the Shape Stimulus Space

We generated complex visual shapes defined by seven radial frequency components (RFCs) (*19–21*) (Fig. 1A). To obtain each shape, sine waves determined by the seven RFCs were added together and the resulting wave was wrapped around a circle to obtain a closed contour which was then filled in to create a shape. We first ran a preliminary pilot experiment to measure how variation along 3 of these 7 components, alone or in combination, was categorically perceived by participants (Fig. S38) and selected the two-dimensional manifold where the pilot participants’ performance matched the parametric midpoint between shape endpoints most closely. We then created a two-dimensional shape space by independently varying the amplitude of the two RFCs from the original set of seven that corresponded to this two-dimensional manifold (from 12.6 to 36.6 for the 4.94 Hz component and from −6.0 to +42.0 for the 1.11 Hz component), while holding the amplitudes of all other five components constant. Distance in this two-dimensional space was computed via changes in amplitude on each axis independently: a displacement of 1 arbitrary distance unit (a.d.u.) corresponded to a change in amplitude of 6 a.d.u. for the 1.11 Hz component and a change in amplitude of 3 a.d.u. for the 4.94 Hz component. This procedure allowed us to generate novel visual shapes for any arbitrarily chosen point within this two-dimensional manifold.

To map the stimulus space perceptually we chose 9 shapes that sat at 0, ±2, ±4, ±6, and ±8 a.d.u. from a fixed center shape along 6 equally spaced radial directions (60 degrees; AG, BH, CI, DJ, EK, FL; Fig. 1A). We recruited 16 healthy individuals (‘norming cohort’) from the Princeton community with normal or corrected-to-normal vision who provided informed consent to a protocol approved by the Princeton University Institutional Review Board and were compensated ($12/hour) to participate in a self-paced two-alternative-forced-choice behavioral experiment, where they were shown 40 repetitions of all 9 shapes from each direction, counterbalanced across runs for left/right placement of the endpoints (e.g., AG vs. GA) and across participants for the order of directions shown. All shapes subtended 5 degrees of visual angle on the screen and the experiment was run using Matlab (2016a) and PsychToolbox (*60*). The endpoint shapes (e.g., A) were also included as catch-trials and we excluded from the analysis six participants who incorrectly categorized more than 10% of the endpoint shapes for any given direction or more than 5% of the endpoint shapes across all directions.

For the remaining 10 participants, for each radial direction, we computed the probabilities (*x*) of categorizing each individual shape as the line endpoints and fitted a corresponding psychometric function (estimated *slope* and *threshold*; Fig. 1B-G):

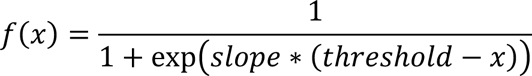

To check that the stimulus space was perceived similarly across all 6 directions, we performed a repeated measures ANOVA on the slopes of the resulting psychometric functions with ‘direction’ as a factor.

All participants in the main experiment (see below, n=10, ‘training cohort’) took part in an identical two-alternative-forced-choice behavioral experiment on Day 1 (see below) of the study. We performed a repeated measures ANOVA on the slopes of the resulting psychometric functions with ‘direction’ and ‘cohort’ as factors to verify that no significant differences existed between the norming and training cohorts (Fig. S1).

For additional details, see corresponding section in Supplementary Text.

### fMRI Localizer Scans to Identify Cognitive Map Brain Regions

The localizer scans (which were not analyzed in real time) involved showing participants sample shapes from the stimulus space in order to (a) test whether we can decode shape category information with high precision from neural patterns (participant exclusion criterion detailed below); (b) find all brain regions that represent the stimulus space as a cognitive map ((*23–24*), prerequisite for defining our neurofeedback target region of interest (ROI)); and (c) to build a neural model of the shape space to be used during real-time neurofeedback training to track how participants’ brains represent the stimuli presented on the screen in each experimental trial.

We performed two scan sessions per participant on separate days. Each scan comprised an anatomical scan (T1 MPRAGE), followed by 4-8 functional (echoplanar) runs (11m23s each, 12-15 total functional runs per participant). During each functional run, participants were shown 49 distinct stimuli from the shape space using a short block design (color: black, background: gray; visual angle: 5 degrees; 3s stimulus presentation) followed by a 10s ISI during which a countdown was displayed (font color: white; background: gray). Shape presentation order was randomized across runs and participants. Since z-scoring with respect to the full response of each run would not be possible during continuous, real-time fMRI acquisition, we included a 72s countdown (font color: white; background: gray) at the beginning of each run and instructed participants to keep their eyes open and stay alert. After eliminating the first 6 TRs to allow T1 equilibration and accounting for hemodynamic lag, this yielded 30 TRs of functional data per run, comparable between all runs (localizer and real-time), which could be used to normalize the signal acquired at each TR (see <u>fMRI Preprocessing for Localizer Scans</u>). Stimuli were displayed on a rear-projection screen (1024×768 resolution, 60 Hz refresh rate) using Matlab (2016a) and PsychToolbox (*60*). Participants viewed the visual display through a mirror mounted on the head coil. For each trial, participants were asked to indicate using an MR-compatible button box whenever a shape ‘oscillated’ (orthogonal task to ensure alertness; see Movie S1 for an example of shape ‘oscillation’ during the neurofeedback training runs). ‘Oscillation’ was defined as a parametric continuous perturbation of the shape in a random direction in the two-dimensional manifold, after which it returned it its original position with the same speed as it was perturbed (250 ms total). Each trial had either 0, 1, or 2 ‘oscillations’, with an average of 1 ‘oscillation’ per trial, randomized across all trials in a run.

For additional details, see corresponding section in Supplementary Text.

### fMRI Acquisition

All structural and functional MRI data were collected on a 3T Siemens Skyra scanner with a 64-channel head coil. Functional images were acquired using an echoplanar imaging sequence (TR, 2000 ms; TE, 28 ms; 36 transversal slices; voxel size, 3×3×3 mm; 64° flip angle; IPAT factor 2), which produced a full brain volume for each participant. Anatomical images were acquired using a T1-weighted MPRAGE sequence, using a GRAPPA acceleration factor of 2 (TR, 2530 ms; TE, 3.3 ms; voxel size, 1×1×1 mm; 176 transversal slices; 7° flip angle). Given the time constraints of real-time fMRI processing and our goal of matching acquisition parameters for classic and real-time scans, we did not correct for susceptibility-induced distortions in any of our echoplanar images.

### fMRI Preprocessing for Localizer Scans

Images were preprocessed using custom AFNI (*61*), Freesurfer (*25*), and bash scripts. All analyses were performed in participants’ native space with no smoothing. The first six volumes of each run were discarded to allow T1 equilibration. For each run, the remaining functional images were spatially realigned to correct for head motion and registered to the participants’ structural T1 image, using boundary-based registration implemented in AFNI’s afni_proc.py. We then performed polynomial trend correction using AFNI’s 3dDeconvolve and simultaneously regressed out 6 degrees of head motion (x, y, z, roll, pitch, yaw). We used FreeSurfer’s recon-all tool to estimate the boundaries of the gray matter for each participant and anatomically defined lateral occipital (LO) and early visual cortex (EVC) regions. We used AFNI’s 3dSurf2Vol and 3dAllineate to obtain corresponding volume masks of the gray matter, LO, and EVC aligned to each participant’s functional data. For each functional run, each voxel was z-scored using the mean and s.d. of its response during the countdown at the beginning of the run (30 TRs).

### Neural Cognitive Map of the Shape Stimulus Space and Neurofeedback Region of Interest

To measure whether object-selective cortex (LO) represents the shape space as a cognitive map ((*23,24*), i.e., similar to how it’s built parametrically and perceived), we computed a representational similarity matrix (RSM) describing the relative relationships between five of the nine shapes in each direction (Fig. 2B; center=0 a.d.u., ±4 a.d.u., ±8 a.d.u.). We compared this RSM using Pearson correlation with an ideal RSM that assumed a parametric linear relationship between the neural activity of the shapes along that direction (Figs. 2C & S2).

To construct our neurofeedback region of interest (ROI), we ran a searchlight analysis (*22*) to find all brain regions for each participant that represented the stimulus space parametrically. First, to increase the probability that potential sources of top-down control (e.g., prefrontal cortex, parietal cortex) would be subsumed by the neurofeedback ROI, we ensured that it must include at least one non-connected cluster of 50+ voxels outside of visual cortex. Second, to ensure that ROI sizes are similar across participants despite natural variability in signal strength, we restricted the final ROI size to 750—2,250 voxels across all participants. Third, to provide evidence that object- and category-selective brain regions in inferior temporal and lateral occipital cortex are causally related to the perception of our shape categories, beyond V1 which we know from prior work can be causally manipulated with neurofeedback (*11*), we excluded an anatomically defined region of interest for early visual cortex defined using FreeSurfer that encompassed V1. To satisfy all three of these constraints, we used an iterative cluster selection procedure to select the optimal threshold for each participant’s searchlight map. Thresholds and neurofeedback ROI sizes for each participant are shown in Table S1. Surface maps of ROIs for each participant are shown in Figs. S3-S12.

For additional details, see corresponding section in Supplementary Text.

### Neural Model of Shape Stimulus Space

To build a neural model of shape categories in the neurofeedback ROI, we modeled each category (12 total) resulting from the partitions in the shape space defined by the 6 radial directions (e.g., AG) as a multivariate Gaussian distribution (Fig. 3B) whose parameters were obtained using maximum likelihood estimation. We used a grid search of number of principal components of the data (100, 150, 200) and hemodynamic lags (4s, 6s) to select the optimal projection and delay (*62*) from stimulus onset that yielded optimal decoding performance for each participant and each ROI.

To account for the small number of training examples, we estimated a shared covariance matrix (Σ_!_) for the two shape categories in each partition:

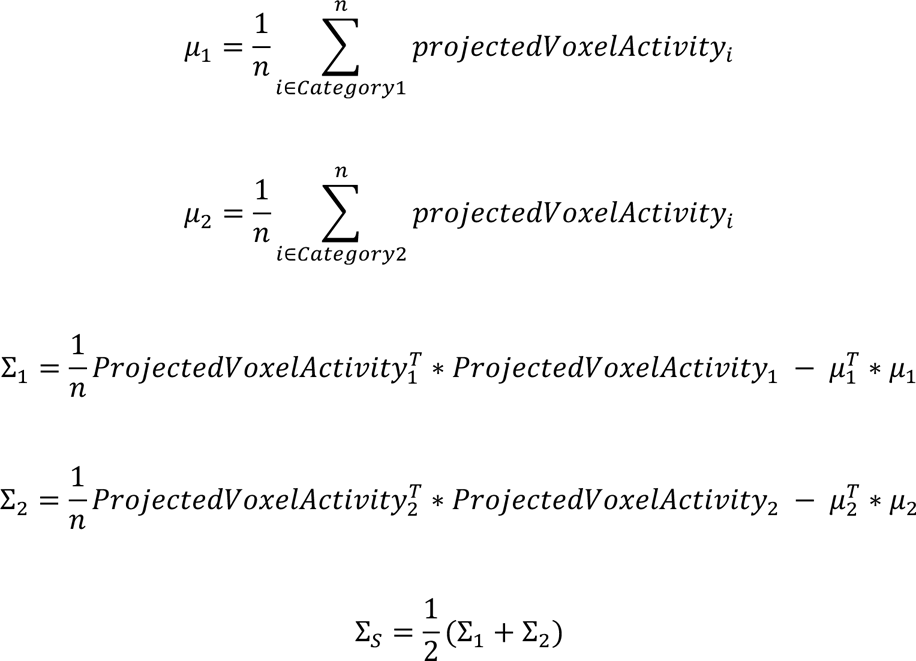

We used our model as a linearly discriminative log-likelihood-ratio-based pattern classifier for neural category representations by computing the probability of a new point *x* (*n*-dimensional neural representation of a shape) under the two estimated category distributions in a given partition:

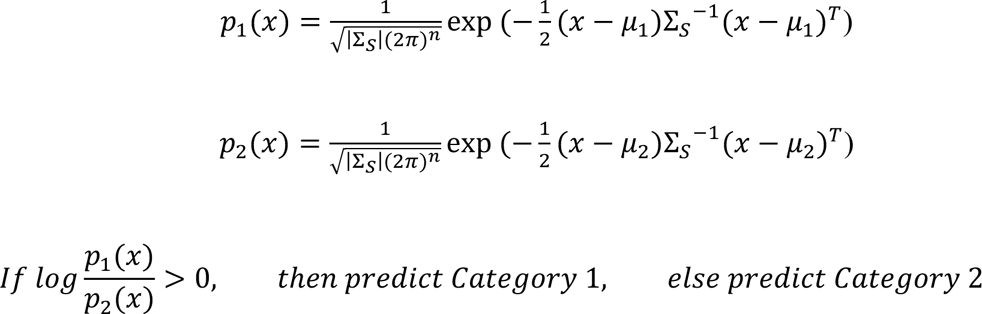

Using our classifier, participants whose average leave-one-run-out cross-validated decoding accuracy across all 6 category partitions was below 70% (chance=50%) in either LO or the neurofeedback ROI were excluded from the neurofeedback training part of the experiment. This exclusion criterion was employed because (1) neurofeedback neural models require high specificity to be able to provide correct feedback during training; and (2) we sought to maximally select for the strongest known predictor of the effectiveness of neurofeedback manipulation across our cohort: neural signal strength as measured by baseline visual category decoding accuracy (*27*). As such, participants whose decoding reliability was below this threshold (70%) in either LO or the neurofeedback ROI were considered to have too low signal to afford reliable real-time neurofeedback training in subsequent sessions of the experiment, as the key outcomes of our study depend on changes in multivariate representations that must be detectable during the training sessions. Decoding results for LO and neurofeedback ROIs and final parameter combination choices for each trained participant are shown in Table S1.

For additional details, see corresponding section in Supplementary Text.

### Neurofeedback fMRI Experiment Design

The full neurofeedback experiment comprised ten total sessions per participant (Fig. 2A):

Session 1: Behavioral 2AFC pre-test

Sessions 2-3: Neural localizer scans / Model training data acquisition

Sessions 4-9: Real-time fMRI neurofeedback training

Session 10: Behavioral 2AFC post-test (identical to pre-test, counterbalanced presentation order)

We designed our experiment with a planned sample size of 10 participants, which is comparable or exceeds similar real-time neurofeedback studies (*11-18*). We recruited and trained participants sequentially until 10 of them both met the initial decoding criterion listed in section <u>Neural Cognitive Map of the Shape Stimulus Space</u> and finished the training and behavioral post-test portion of the study in its entirety (i.e., without dropping out of the study voluntarily). As such, we recruited a total of twenty-five healthy participants over the course of 17 months (May 2018-October 2019; 13 female, 23 right-handed, ages 18-35, mean age 22.4) from the Princeton community with normal or corrected-to-normal vision who provided informed consent to a protocol approved by the Princeton University Institutional Review Board and who were compensated for their participation ($12/h for behavioral sessions, $20/h for fMRI sessions, retention incentives: $5/session cumulative bonus for each fMRI session beyond the first, $0.10 per successful neurofeedback trial, $100 for completing the full experiment. If the participant had satisfactory performance on the behavioral test (using criteria listed in section <u>Building and</u> <u>Norming the Shape Stimulus Space</u>) and if a neural model with high decoding accuracy could be built using the neural data from the two localizer scans (using criteria listed in section <u>Neural</u> <u>Cognitive Map of the Shape Stimulus Space</u>), then the participant was invited to take part in the rest of the experiment. Of our initial set of twenty-five participants, two were rejected for not meeting MRI safety criteria, eight were rejected for poor behavioral and/or neural decoding performance during sessions 1—3, and five voluntarily withdrew their participation after 1—3 sessions. Ten participants (7 female, all right-handed, ages 20-35, mean age 24.0) took part in the full experiment, two of which were trained for 5 days and eight of which were trained for 6 days. Participant performance during the localizer sessions and reasons for exclusion are shown in Table S1.

Each session of the experiment was run on a separate day. To minimize fatigue and afford potential benefits from overnight memory consolidation (*63*), we sought to have participants come into the lab for at most 2 hours each day on consecutive days during the training part of the experiment. Participants performed a variable number of trials per training day, depending on the amount of time the scanner setup and participant pre-scan procedures took (minimum 20 trials, maximum 140 trials per day). We ensured that participants performed at least 500 total neurofeedback training trials before the final behavioral post-test. To minimize potential effects of over-training and to guarantee a comparable amount of training across our entire cohort, we stopped the training procedure once a participant performed 800 trials (two participants). Three participants were allowed one 48h gap each during the experiment for emergencies and/or personal reasons.

### Real-Time fMRI Neurofeedback Procedure

For each participant, we randomly choose one of the 6 radial category partitions by rolling a pink six-sided die to become their training category boundary (information hidden from participants).

We performed 5—6 neurofeedback training scan sessions per participant (Table S3), each comprising an anatomical scan (T1 MPRAGE), a functional (echoplanar) localizer run (11m37s, identical to Days 2—3), and 1—7 functional (echoplanar) neurofeedback training runs (10m32s each) for a total of 26-40 neurofeedback runs per participant (in-scanner training time: 4.6h-7.0h per participant). The training runs used a block design with an initial 72s countdown (analogous to the localizer scans), followed by twenty trials (16s=8TRs stimulus presentation, 12s=6TRs ITI). Stimuli (color: black; background: gray; visual angle: 5 degrees) were displayed on a rear-projection screen (1024×768 resolution, 60 Hz refresh rate) using Matlab (version 2016a) and PsychToolbox and were viewed through a mirror that was mounted on the head coil. Each trial comprised a random shape (chosen i.i.d. from the shape space), ten total from each category, shown in random order. Since preliminary tests showed that the decoding performance of the neural shape model deteriorated for shapes close to the category boundary (Fig. S15), we chose shapes at least 1 a.d.u. away from both the category boundary and from the perpendicular direction in shape space. Shapes oscillated continuously at a rate of 500ms (4 independent oscillations per 2s TR, Movie S1).

The radius of the oscillations in each TR was the main method by which neurofeedback was given to study participants. They were told that during each trial they would see a continuously oscillating shape and given the following set of instructions:

1. “Generate a mental state that’s going to make each shape oscillate / wobble less or even stop!”;
2. “Different shapes may require different strategies.”; and
3. “When you are successful in slowing down the shapes, it has to do with your mental state over the past 8-10 seconds; progress is not instantaneous.”

The first instruction encouraged participants to generate mental state variability that may help shift representations in the neurofeedback ROI. The second instruction addressed the possibility that shifting representations in neural space in different (opposing) directions of a cognitive map may require different types of top-down influences; while it may have alerted participants to the possibility that multiple categories of shapes exist, the high variability of the shape space was designed to make boundaries between such categories difficult to intuit. The final instruction was necessary due to hemodynamic lag inherent to the fMRI signal, which together with its temporal autocorrelation properties would invariably influence the visual feedback. As participants attempted to generate such a target mental state during each TR (2s), we preprocessed their functional data (see <u>fMRI Preprocessing for Real-Time Scans</u>), and used our neural model to identify how far from the neural category boundary the shape was represented in the participants’ neurofeedback ROI; if the shape was strongly represented as a member of its ground truth category (known to the experimenter, but not to the participant), then positive feedback was given to the participant by shrinking the radius of the oscillation (causing it to look less extreme, Movie S1, trial 2), otherwise no feedback was given and the participant saw the oscillation continue with the same amplitude (see <u>Neurofeedback fMRI Data Analysis</u>). Each shape was presented for a total of 8 TRs. During the first 3 TRs, no feedback was given since hemodynamic lag prevented it. For the remaining 5 TRs, participants received potential feedback on a TR-by-TR basis. Feedback was cumulative. Participants also received a monetary reward ($0.10) with positive feedback and a bonus of $0.25 if they stopped the oscillation completely (positive feedback for 3+ TRs per trial).

For additional details, see corresponding section in Supplementary Text.

### fMRI Preprocessing for Real-Time Scans

fMRI data were collected similarly to the localizer scans (equipment, parameters, etc.) and the images were preprocessed analogously. We used AFNI’s afni_proc.py script to align each functional run to the anatomical scan matching both the offline neural shape model and the neurofeedback ROI mask. This allowed fast recovery of the shape-specific functional data from each acquired volume and ensured that the neural model could be evaluated in real time.

Given the constraints of the real-time environment, classical preprocessing techniques and normalization procedures could not be implemented (e.g., polynomial trend regression, motion parameter regression, z-scoring with respect to the entire timecourse). Instead, the functional data acquired up to a given TR were temporally high-pass filtered (e.g., on TR 45, the first 45 TRs were filtered together) with a 52s (26 TRs) period cutoff (the length of two consecutive trials) using a fast custom script written in C++. During each TR of a functional run, each voxel’s timecourse was z-scored using the mean and s.d. of its response during the countdown at the beginning of the run (30 TRs).

For additional details, see corresponding section in Supplementary Text.

### Neurofeedback Online fMRI Data Analysis

For each TR, the neural model was used to estimate the log-likelihood ratio (LLR) of the neural activity elicited by the current stimulus under that participant’s individual Trained category boundary. At the beginning of each trial, the default oscillation radius (see <u>Real-Time fMRI</u> <u>Neurofeedback Procedure</u>) was set to 1.875 a.d.u. If the LLR for a given TR was above a particular threshold (described below), then the amplitude of the oscillation was reduced during the subsequent TR by 0.625 a.d.u., otherwise the amplitude was kept unchanged. Feedback was cumulative (Fig. S14). The LLR threshold for the first training run was initially set to the 70^th^ percentile of the distribution of localizer data for that participant. In subsequent runs, the threshold was adjusted using an adaptive procedure based on participant performance (how many trials generated positive feedback) on all previous runs since the beginning of the training day as detailed in Table S2. To avoid potentially random feedback being given due to classifier uncertainty near the category boundary (Fig. S15), the threshold was never lowered below 60% of the localizer scan LLR distribution, regardless of participant performance. The threshold was also kept unchanged between the last run of a training day and the first run of the subsequent day, regardless of performance during the former. The LLR distribution for a representative participant and a sample threshold (75%) are shown in Fig. 3D. Distributions and sample thresholds for all participants are shown in Fig. S13.

For additional details, see corresponding section in Supplementary Text.

### Post-Experiment fMRI Data Analysis

To measure relative neural change as a consequence of training, we computed a participant-level summary statistic of the effect of neurofeedback training by averaging the LLR across the first two days (pre) and the last two days (post), separately for the Trained (e.g., ABCDEF vs. GHIJKL) and Control (orthogonal, e.g., DEFGHI vs. JKLABC) category boundaries.

First, we used a difference score to compute the LLR change (Fig. 4A):

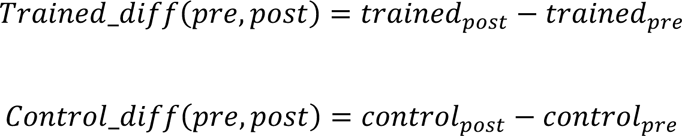

We then used a paired t-test to investigate whether training induced different changes for the Trained and Control category distinctions and computed Cohen’s d to measure the effect size of this change between Trained and Control categories. We also used two separate statistical tests to verify that the main neural effect values were normally distributed: Shapiro-Wilk p=0.256, Epps-Pulley p=0.097. To maximize sensitivity of this measure given potential fMRI adaptation effects (*28, 29*), we focused on LLR values from the first TR of each trial only as the most reliable measure of LLR across the neurofeedback training runs. Results using the first 2, 3, 4, and 5 TRs of each trial were highly similar (Table S4).

Second, we computed the ratio between Trained and Control LLR for the pre and post conditions:

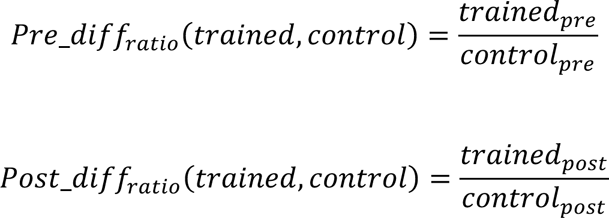

We then used a paired t-test to investigate whether the LLR ratio changed as a consequence of training between the first two days and the last two days of the experiment (Fig. S17).

To measure potential changes in neural category patterns across our experiment, we computed Pearson correlations with the ideal parametric similarity matrix and with the ideal categorical similarity matrix (Figs. 2C & S22) for patterns of activity elicited in LO and in the neurofeedback ROI during the single functional localizer run from the beginning of each in each neurofeedback fMRI training session (see <u>Real-Time fMRI Neurofeedback Procedure</u>), for the first two vs. last two days of training (1 run per participant per day).

To measure the strength of category representations within sub-regions of LO and of the neurofeedback ROI, we (a) split LO into equal sized anterior (aLO) and posterior (pLO) halves using the Z coordinate of its Freesurfer map; and (b) split the neurofeedback ROI into “visual” (vis-ROI, all voxels within occipito-temporal cortex) and “high-level” (high-ROI, all remaining ROI voxels outside occipito-temporal cortex) components. We then trained new category decoding models within these sub-regions for the data collected during the localizer scans. To quantify changes in decoding accuracy due to training, we computed the decoding accuracy of these models for the Trained and Control directions, across the calibration runs collected during first two days and the last two days of training. We also computed the difference of differences for this latter quantity (Fig. S22).

### Behavioral Tests, Debrief, and Post-Experiment Data Analysis

To evaluate the influence of training on perception, we conducted two 2AFC experiments (procedure identical to <u>Building and Norming the Shape Stimulus Space</u>) during the first and last days of the experiment and we estimated psychometric function slopes for each of the 6 radial directions in our stimulus space separately before and after training. To compare the strength of category separation for each participant between the pre- and post-tests, we computed the average slope for the two directions most perpendicular to the training category boundary (Trained, e.g., for categories ABCDEF and GHIJKL, the average slope of lines CI and DJ) and the average slope for the two directions most parallel to the training category boundary (Control, e.g., for categories ABCDEF and GHIJKL, the average slope of lines AG and FL) (Fig. 4A). We also computed this change for the average of the two directions that were neither parallel, nor perpendicular to the trained category boundary (Neutral, e.g., for categories ABCDEF and GHIJKL, the average slope of lines BH and EK) (Fig. S20). We used a normalized difference score to compute the relative change in psychometric function slope before and after training:

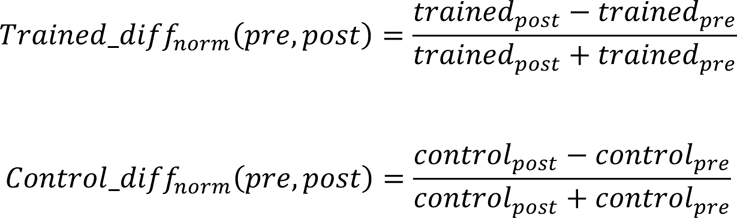

We computed a two-sided paired t-test between Trained and Control slopes to evaluate whether our neurofeedback training procedure induced a significant behavioral change consistent with our hypothesis and we computed Cohen’s d to measure the effect size of this behavioral change. We also used two separate statistical tests to verify that the main behavioral effect values were normally distributed: Shapiro-Wilk p=0.386, Epps-Pulley p=0.076. Estimated psychometric functions for all tests, directions, and participants are shown in Table S5.

To measure whether a relationship exists between neural change and perception, we computed the Pearson and Spearman correlation between the LLR change and the behavioral change, i.e., how well does LLR change in the neurofeedback ROI predict psychometric function change for the Trained vs. Control direction in behavior.

To measure whether lapses in performance between the pre- (Day 1) and post-tests (Day 10) could potentially explain the effects of neurofeedback training on perception, we also performed a separate joint estimate of the lapses (upper and lower), thresholds, and slopes for our psychometric functions (parameter estimates: Table S6; psychometric functions: Fig. S36):

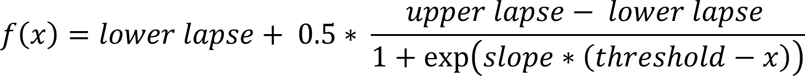

We also used a linear mixed effects model to measure whether the effect of neurofeedback on the separate slope change difference estimate remained significant after controlling for threshold and lapses.

To measure whether our results are robust in individuals and at the group level with our sample size (n=10), we performed bootstrap resampling analyses for (1) the neural change elicited by the neurofeedback manipulation; (2) the behavioral change observed between Day 1 and Day 10 of the experiment; and (3) the correlation between neural change and behavioral change. We used a sample-with-replacement procedure to generate 10,000 random draws of 10 samples (i.e., 10 participants) from the distributions of observed neural and behavioral effects (Fig. 4), and subsequently computed the Pearson correlation of each matched sample pair. Histograms of the sample values for the neural, behavioral, and correlations are shown in Fig. S21. To further investigate the robustness of our observed correlation values between neural and behavioral effects of training, we computed null distributions for the Pearson and Spearman correlation values by keeping the neural effects fixed and randomizing the order of the behavioral effects 10,000 times. Histograms of the null distributions are shown in Fig. S35.

To investigate if participant training outcomes may be related to individual differences in access to neural information via computational analysis of fMRI data, we defined the baseline neural decoding measure for each participant as the classifier decoding accuracy in the neurofeedback ROI during the two localizer scans (Days 2—3), averaged across all six directions of the stimulus space (Fig. 3C). We then computed the correlation between baseline decoding for each participant and their LLR neural changes, as well as their behavioral outcomes.

After the two-alternative-forced-choice behavioral post-test was completed (but before full debrief), we asked participants to fill out a questionnaire about their experience, including whether they suspected there were multiple categories of shapes; if so, how many and which ones; and asking them to comment on strategies they used to perform the task. The questionnaire and a summary of the answers are shown in Table S7. The free guess category boundaries and the 2-category-forced-choice boundaries are shown in Figs. S18 & S19.

We also performed a linear mixed effects model analysis to measure whether behavioral changes observed in our cohort were significantly predicted by neural changes when accounting for the effect of number of training days (5 or 6), the baseline decoding accuracy during the localizer scans, the random category split assigned to each participant, the guessed angle displacement from the correct category boundary reported at the end of the study, and the participants’ ages.

To measure whether the distribution of positive feedback trials throughout the experiment could explain our results, we also performed ideal observer analyses to measure whether the feedback values received during training would be sufficient to predict the participants’ neural or behavioral changes or to explain the participants’ forced choice guesses about the category boundary at the end of the experiment. First, we computed the proportion of feedback points that fell into the two sets of putative categories defined by the Trained and Control boundaries, and we normalized the amount of feedback to a proportion for each category in each condition. Then, we computed D1 as the difference in the proportion of positive feedback received on the two sides of the Trained boundary and D2 as the difference in the proportion of positive feedback received on the two sides of the Control (perpendicular) boundary. Using a two-sided t-test and a Kolmogorov-Smirnov test, we measured whether the proportion of feedback differs between the Trained and Control boundaries (D1 vs. D2). We also computed the Pearson correlation between the difference of differences (D3=D2-D1) and the neural and behavioral changes observed after training. Second, to further test whether an ideal observer model could have used the feedback received during training to guess the correct shape category boundary, we trained a linear SVM classifier to predict shape category based on the feedback values received during training.

### Data and Materials Availability

fMRI data for all 78 scanning sessions of the study is publicly available in BIDS format in the following NIH repository: https://nda.nih.gov/study.html?id=1098. Behavioral data and experiment code, including for real-time display and processing of fMRI data, are publicly available in the following GitHub repository: https://github.com/NatCogLab/neuralsculpting.

