## Supporting Information for "Sculpting New Visual Categories into the Human Brain"

**This PDF includes:**

Supporting Text

Supplementary Figures S1-S38

Supplementary Tables S1-S7

Legends for Supplementary Movies S1-S2

Supporting Information References

**Other supporting materials for this manuscript include the following:**

Supplementary Movies S1-S2

### Supporting Information Text

Building and Norming the Shape Stimulus Space

To map the stimulus space perceptually, we chose shapes that sat at ±2, ±4, ±6, and ±8 arbitrary distance units from a fixed center shape along 6 equally spaced radial directions (60 degrees; AG, BH, CI, DJ, EK, FL; Fig. 1A). This yielded 9 equally spaced shapes along each of 6 radial directions, including the common center shape (e.g., A and G are the 8 a.d.u. shapes from center for the AG direction). We then recruited 16 healthy individuals ('norming cohort') from the Princeton community with normal or corrected-to-normal vision who provided informed consent to a protocol approved by the Princeton University Institutional Review Board and were compensated ($12 per hour) to participate in a self-paced two-alternative-forced-choice behavioral experiment. For each radial direction, participants were told that the endpoints (e.g., shapes A and G, 8-distance-units away from center, shown continuously on the left and right of the screen) correspond to two different categories of shapes and they were instructed to use this information to categorize new shapes that appeared in the center of the screen. New shapes were drawn randomly from the 9 shapes chosen for that particular radial direction (see above) such that 40 repetitions of all 9 shapes were shown in each run of the experiment. To ensure that categorization effects were not influenced by the left/right placement of the endpoints on the screen, each participant performed two separate runs for each radial direction (e.g., AG vs. GA) and the order of the runs and assignment of endpoints to each run was counterbalanced across participants. The endpoint shapes (identical to one of the 'categories') were also included as catch-trials and we excluded from the analysis six participants who incorrectly categorized more than 10% of the endpoint shapes for any given direction (criterion: >16 out of 160; 2 participants excluded: 29/160 and 67/160) or more than 5% of the endpoint shapes overall across all directions (criterion: >48 out of 960; 4 participants excluded: 77/960, 84/960, 93/960, and 456/960).

fMRI Localizer Scans to Identify Cognitive Map Regions

We performed two scan sessions per participant on separate days. To help minimize in-scanner motion and encourage participants to self-regulate this potential source of noise during the scans, we chose to avoid the use of rigid restraints (for comfort, given the length and number of sessions), but did use masking tape to both gently tether the participants' heads to the coil, and to provide physical feedback whenever they attempted to move during the experiment. Each scan comprised an anatomical scan (T1 MPRAGE acquisition), followed by 4-8 functional (echoplanar) runs (11m23s each) for a total of 12-15 total functional runs collected per participant across the two scan days.

For each localizer run, similarly to the shape space norming experiment, we chose shapes that sat at ±2, ±4, ±6, and ±8 arbitrary distance units from a fixed center shape along 6 equally spaced radial directions (60 degrees; AG, BH, CI, DJ, EK, FL; Fig. 1A). This yielded 9 equally spaced shapes along each of 6 radial directions, including the common center shape (e.g., A and G are the 8 a.d.u. shapes from center for the AG direction), for a total of 49 unique shapes. The presentation order was randomized across runs and across participants. During each unique shape presentation (3s), participants were asked to indicate using an MR-compatible button box whenever a shape 'oscillated' (orthogonal task to ensure alertness). An 'oscillation' was defined as a parametric continuous perturbation of the shape in a random direction in the parametric manifold, after which it returned it its original position with the same speed that it was perturbed

(250 ms total, i.e., we picked a random point in parameter space near the original shape chosen by drawing an i.i.d. sample from the circumference of a circle of radius 2 arbitrary distance units centered at the shape, then we created an animated morph between the original shape and the one defined by the point on the circle, and back to the original shape). Each trial had either 0, 1, or 2 'oscillations', with an average of 1 'oscillation' per trial, randomized across all trials in a run. The absolute position of the center of mass of the shapes was always held constant at the center of the screen; the oscillation in parameter space simply created the visual impression that the current shape morphed into a similar shape and then reverted to the original (see Movie S1 for a demo of continuous oscillation).

Neural Cognitive Map of the Shape Stimulus Space and Neurofeedback Region of Interest

To construct our neurofeedback region of interest (ROI), we sought all brain regions in each participant's brain that represented the stimulus space parametrically and ran a searchlight analysis (Kriegeskorte et al., 2006) (cube of side length 7 voxels intersected with gray matter volume, minimum 172 voxels, distance between cube centers = 2 voxels) across their entire gray matter mask. In each voxel cube, we computed the Pearson correlation with the ideal RSM (Fig. 2D). First, to increase the probability that potential sources of top-down control (e.g., prefrontal cortex, parietal cortex, etc.) would be subsumed by the neurofeedback ROI, we ensured that it must include at least one non-connected cluster of 50 or more voxels outside of visual cortex (i.e., not connected to LO). Thus, if such voxels would be amenable to neurofeedback manipulation, and if changes in these voxels were causally related to perception, then including them in our neurofeedback ROI would maximize our chances to elicit a behavioral effect in our participants. Second, to ensure that ROI sizes are relatively similar across participants despite natural variability in signal strength, we restricted the final ROI size to between 750 and 2,250 voxels across all participants. Third, to provide evidence that object- and category-selective brain regions in inferior temporal and lateral occipital cortex are causally related to the perception of our shape categories, beyond V1 which we know from prior work can be causally manipulated with neurofeedback (Shibata et al., 2011), we excluded an anatomically defined region of interest for early visual cortex (EVC) defined using FreeSurfer that encompassed V1. To satisfy all three of these constraints, we started by selecting all clusters of 20 or more voxels that had an $r>=0.50$ with the ideal RSM, excluding the EVC. If the conditions were met, this became our neurofeedback ROI. Otherwise, we either relaxed or increased the r threshold stepwise using values from the set {0.25, 0.33, 0.40, 0.50, 0.60, 0.66} until both of the conditions were met for each individual participant.

Neural Model of Shape Stimulus Space

To measure how the shape space was categorically represented in the neurofeedback ROI, we split the stimulus space into groups of two categories each comprising half of the shape space circle (Fig. 3A). To maximize the number of training examples available, the category boundaries were chosen halfway between the diameters defined in Fig. 1A, e.g., halfway between lines AG and BH, halfway between lines BH and CI, etc. This yielded 6 categorical partitions of the shape space, each associated with 24 out of 49 unique trials per localizer experiment run (24 shapes shown belonged to one category partition, 24 belonged to the other partition, and the center shape was on the dividing category boundary line, and thus was not used for estimating a category model). For each trial in each functional localizer run, we selected two TRs following each short block onset (accommodating for optimal activation given hemodynamic lag, see below) and considered them as independent training points in order to maximize the amount of training data available. This procedure yielded 48 training points per category per functional run, with a total of 576—720 training points per category per run for each participant and each of the 6 categorical partitions (i.e., ABCDEF vs. GHIJKL, BCDEFG vs. HIJKLA, CDEFGHI vs. JKLABC, etc.). To build a neural model, we modeled each categorical partition as a multivariate Gaussian distribution. The parameters of these category-specific distributions were computed using maximum likelihood estimation using the localizer scan data associated with each category partition (schematic shown in Fig. 3B). Given the high dimensionality of the training data (750—2,250 voxels per participant) and the relatively small number of training examples ($n$=576—720 per category per participant), we used a dimensionality reduction procedure to project the training data (voxel activations) onto its first 100, 150, and 200 principal components ($ProjectedVoxelActivity_k=\{projectedVoxelActivity_i\},\ k\in\{1,2\}$) and estimated a shared covariance matrix ($\Sigma_S$) for both shape categories in each partition.

Additionally, given potential variability in hemodynamic lag between brain regions and across participants (Handwerker et al., 2012), we also selected two potential lags: the training data comprised either the 3rd and 4th TRs following short block onset (4s lag) or the 4th and 5th TRs following short block onset (6s lag). We built neural category models for each direction in each participant using all six combinations of these two parameters (PC dimensions = 100, 150, 200 x hemodynamic lag = 4s, 6s) in both LO and the neurofeedback ROI. To verify that our model could correctly predict the distinction between shape categories in the brain, we used it as a linearly discriminative log-likelihood-ratio-based pattern classifier.

Using our classifier, participants whose average leave-one-run-out cross-validated decoding accuracy across all 6 category partitions was below 70% (chance=50%) in either LO or the neurofeedback ROI using all possible parameter combinations were excluded from participating in the training portion of the experiment. For all participants that met the threshold, we selected the parameter combination (PC dimensions x hemodynamic lag) that gave the highest average decoding accuracy across all 6 categorical partitions in the neurofeedback ROI (Fig. 3C), regardless of performance in LO. Localizer sessions decoding results for LO and neurofeedback ROIs, as well as final parameter combination choices for each trained participant are shown in Table S1.

### Real-Time fMRI Neurofeedback Procedure

For each participant, we randomly choose one of the 6 radial category partitions (by rolling a pink six-sided die) to become their training category boundary (information hidden from participants), while ensuring that neural decoding accuracy was at least 70% for this direction and that it hadn't been chosen before for another participant (after we trained 6 participants on all 6 distinct partitions, we reset this constraint).

We performed five to six neurofeedback training scan sessions per participant on separate days. The radius of the oscillations in each TR was the main method by which neurofeedback was given to study participants. Each shape was presented for a total of 8 TRs. During the first 3 TRs, no feedback was given since hemodynamic lag prevented it. Afterwards, for the remaining 5 TRs of the block, participants received appropriate (or no) feedback on a TR-by-TR basis. Feedback was cumulative, such that if the participant maintained a mental state for the shape that took it away from the category boundary for 3 TRs out of the 5 TRs in which feedback is given, then the shape actually stopped oscillating altogether (radius of oscillation decreased to zero). In practice, this was extremely rare, but did occur sometimes, especially towards the end of the experiment. We limited each training trial block to 16s (8 TRs) to eschew potential fMRI adaptation effects that might diminish our ability to apply our neural model and/or decode the category of the shape on the screen (Grill-Spector & Malach, 2001; Aguirre, 2007).

fMRI Preprocessing for Real-Time Scans

Structural and functional MRI data were collected in an identical fashion to the localizer scans (coil, equipment, parameters, etc.). The images were preprocessed using custom AFNI (Cox, 1996), Matlab (version 2016a), C++, and bash scripts. All analyses were performed in participants' native space and no smoothing was applied. The first six volumes of each run were discarded to allow T1 equilibration. We used AFNI's afni_proc.py script to preprocess the localizer run from each training day scan (analogous to fMRI Preprocessing for Localizer Scans), as well as align this functional run to the anatomical scan matching the Sessions 2-3 data used to generate the offline neural shape model. This allowed us to generate a template image (the 50$^{th}$ volume of the localizer EPI run) to which we would be able to align new volumes acquired during the neurofeedback training runs in a reasonable amount of time (aligning to an anatomical scan or to a full functional run usually requires tens of seconds and is not useable in a real-time setting but aligning a single volume to another single volume requires <500ms on average). This template volume was also aligned to the neurofeedback ROI mask, which allowed fast recovery of the relevant functional data from each real-time acquired volume after fast single-volume-to-single-volume alignment, such that the neural model could subsequently be evaluated in real time.

The real-time functional scans were transferred via a fast network connection from the scanning console (Siemens) memory directly to the hard drive of a high-powered server on which all analyses were performed. Each real-time volume was aligned to the template volume (see above) using AFNI's 3dvolreg and the neurofeedback ROI mask was used to subsequently extract the relevant functional data.

Neurofeedback Online fMRI Data Analysis

The functional data from the neurofeedback ROI was projected into the dimensionality-reduced space defined during the procedure used to select the optimal number of PC dimensions and hemodynamic lag for shape category decoding (see Neural Model of Shape Stimulus Space). The multivariate Gaussian neural model was then used to estimate the log-likelihood ratio (LLR) of the current data point (representing the neural activity elicited by the current stimulus being shown on the screen in the scanner) for each of the two categories of shapes (e.g., ABCDEF vs. GHIJKL) selected for training for the current participant. At the beginning of each trial, the default oscillation radius (see Real-Time fMRI Neurofeedback Procedure) was set to 1.875 a.d.u. based on a trade-off between the visual salience of the perturbation and pilot results measuring how the amplitude degraded decoding (Fig. S15). If the LLR for a given TR was above a particular threshold (described below), then the amplitude of the oscillation was reduced during the subsequent TR by 0.625 a.d.u., otherwise the amplitude was kept unchanged. Feedback was cumulative, allowing the amplitude to become zero (thus causing the shape to become static) after

3 cumulative (but not necessarily consecutive) TRs of positive feedback during a trial (Fig. S14). The LLR threshold for the first training run was initially set to the 70$^{th}$ percentile of the distribution of all data points (1,152—1,440 per participant) collected during the localizer scans for that participant under the offline Gaussian neural model. After the first training run, the threshold was potentially adjusted using an adaptive procedure for two reasons. First, since the only way in which participants can learn to shift their neural representations is through receiving feedback via the shape oscillation radius, we sought to converge on giving feedback for approximately 1/3 of the twenty trials in each run. Second, given the high variability of conditions elicited by fMRI scanning across multiple days in terms of signal strength and sources of external noise, we expected our ability to decode using a static neural model to vary across the training week, as well as across training runs. As such, an adaptable threshold can recover from being too stringent (and never providing feedback) or too lax (providing feedback on almost every trial), both of which would be undesirable outcomes of the training procedure. Thus, the threshold was adjusted given participant performance (how many trials generated positive feedback) on all previous runs since the beginning of the training day as detailed in Table S2. By design, positive feedback could only be given on trials where the model correctly guessed the ground truth category of the shape being shown.

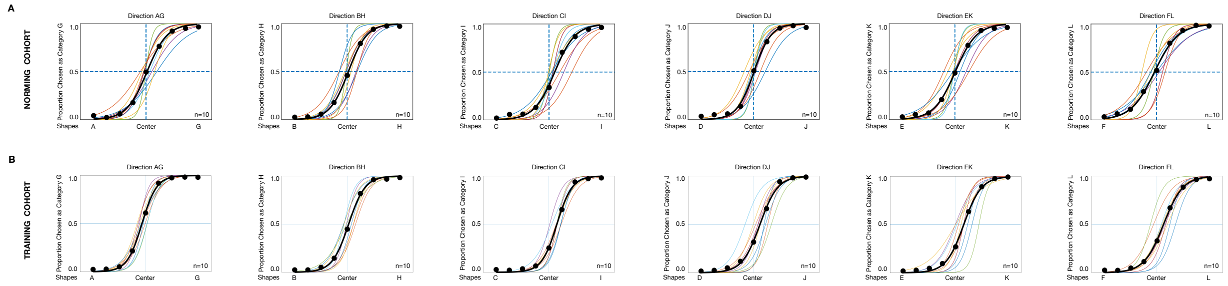


**Fig. S1. Stimulus space was perceived identically by norming and training cohorts of participants.**

Psychometric functions estimated for each of the six directions in the stimulus space (AG, BH, CI, DJ, EK, FL) for the norming cohort (A, n=10, replicated from Fig. 1B-G for convenience) and the training cohort (B, n=10, bottom). Color lines indicate individual participants and bolded line indicates average. No significant differences were observed between the two cohorts or between the six directions (psychometric function slope, repeated measures ANOVA, factors='direction', 'cohort'; interaction: $F(5,108)=0.19$, $p=0.964$; main effect of direction: $F(5,108)=0.57$, $p=0.722$; main effect of cohort: $F(1,108)=1.71$, $p=0.194$).

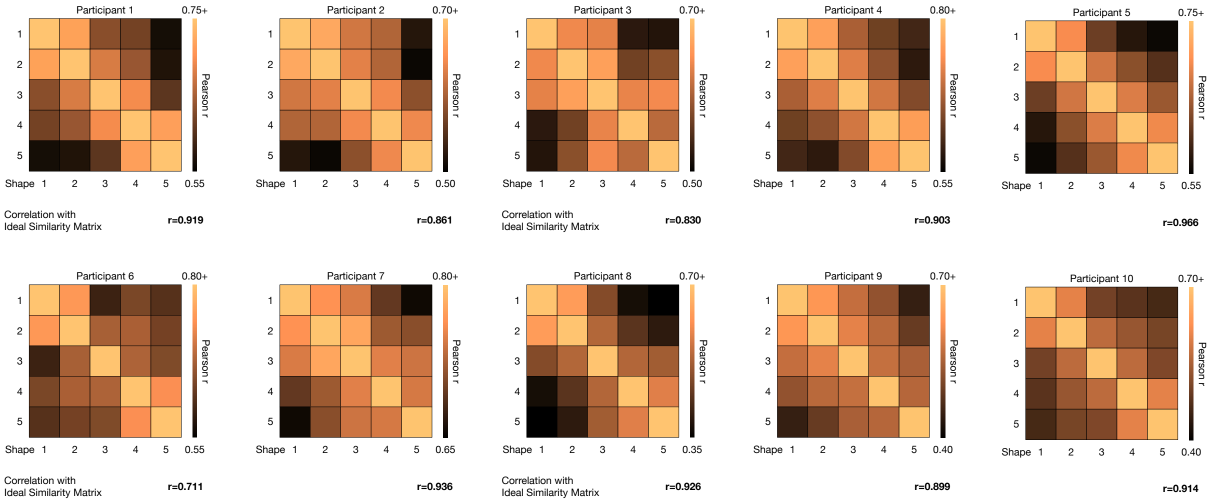


**Fig. S2. Shape space is represented parametrically in neurofeedback ROI.**

Representational similarity matrices (RSM) obtained by computing the Pearson correlation between neural activity elicited by pairs of shapes spanning the stimulus space (Fig. 2B) in the neurofeedback ROI of each participant during the localizer scans (Days 2-3). Correlation with the ideal representational similarity matrix RSM (Fig. 2C) is shown below each participant RSM.

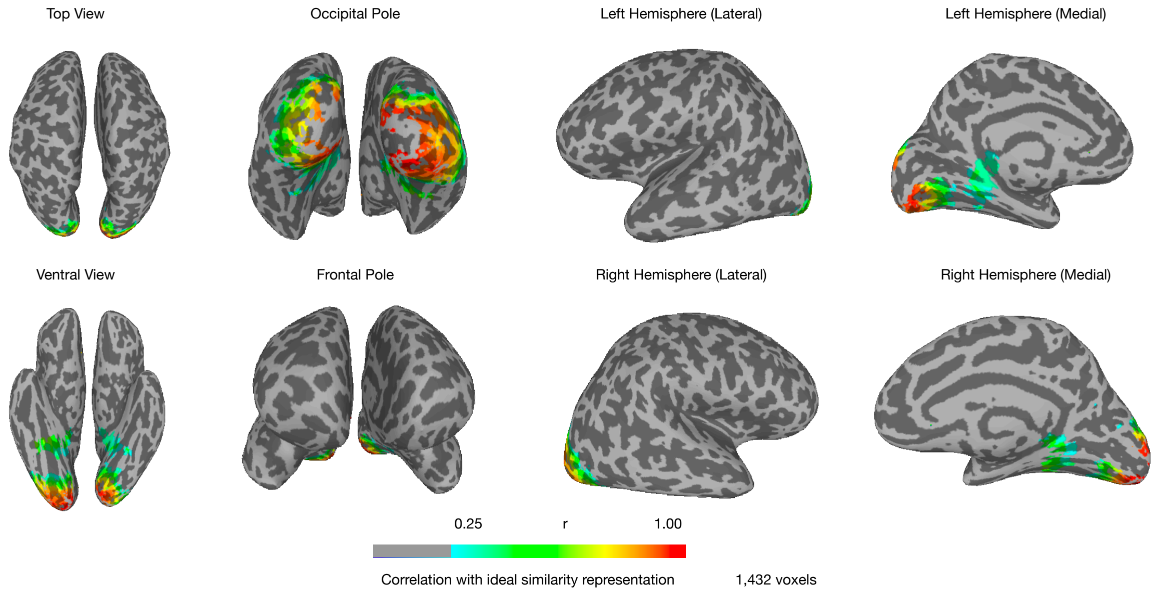


**Fig. S3. Neurofeedback ROI for participant #1.**

Searchlight analysis indicating all of participant #1's brain regions that represented the shape space parametrically, including extrastriate visual cortex, parahippocampal gyrus, and hippocampus. See Methods for details of the searchlight procedure and threshold selection.

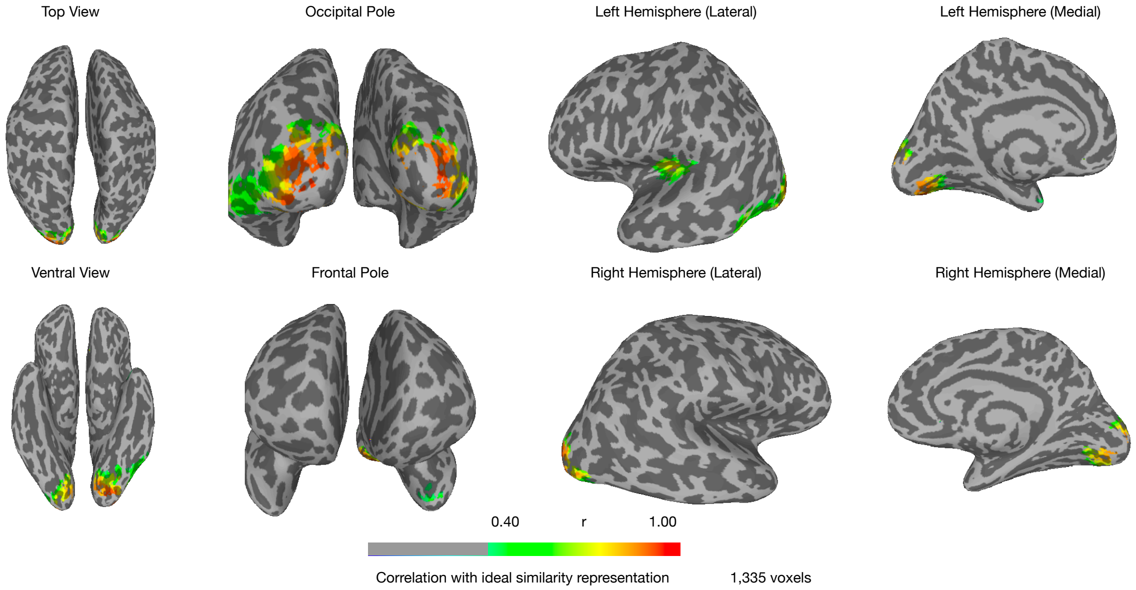


**Fig. S4. Neurofeedback ROI for participant #2.**

Searchlight analysis indicating all of participant #2's brain regions that represented the shape space parametrically, including extrastriate visual cortex, inferior temporal cortex, anterior temporal cortex, and superior temporal sulcus. See Methods for details of the searchlight procedure and threshold selection.

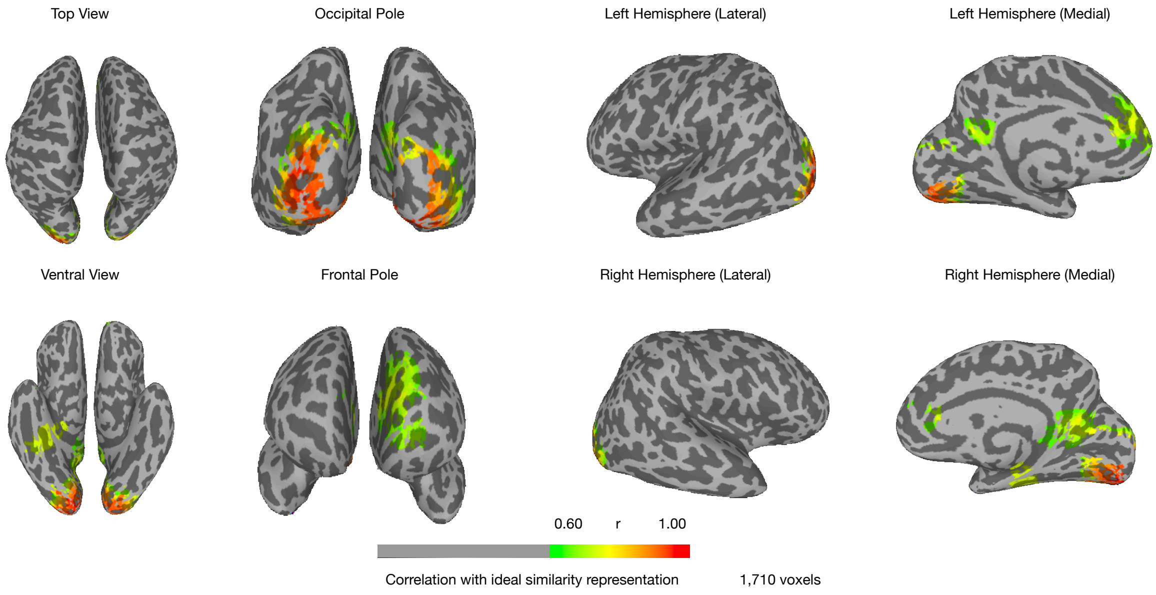


**Fig. S5. Neurofeedback ROI for participant #3.**

Searchlight analysis indicating all of participant #3's brain regions that represented the shape space parametrically, including extrastriate visual cortex, parahippocampal gyrus, hippocampus, and medial frontal gyrus. See Methods for details of the searchlight procedure and threshold selection.

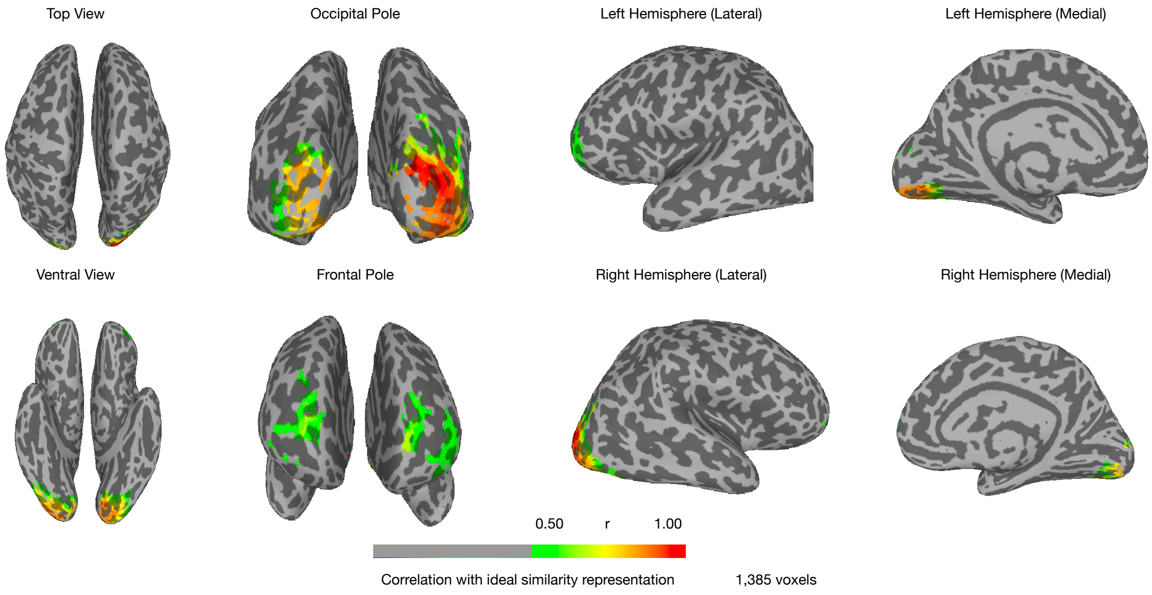


**Fig. S6. Neurofeedback ROI for participant #4.**

Searchlight analysis indicating all of participant #4's brain regions that represented the shape space parametrically, including extrastriate visual cortex, transverse occipital sulcus, and dorsolateral prefrontal cortex. See Methods for details of the searchlight procedure and threshold selection.

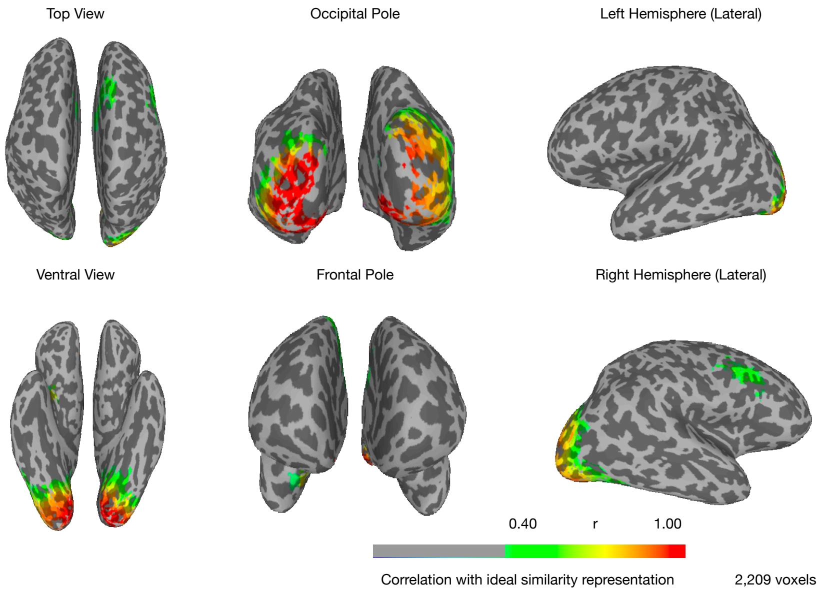


**Fig. S7. Neurofeedback ROI for participant #5.**

Searchlight analysis indicating all of participant #5's brain regions that represented the shape space parametrically, including extrastriate visual cortex, inferior temporal cortex, parahippocampal gyrus, and dorsolateral prefrontal cortex. See Methods for details of the searchlight procedure and threshold selection.

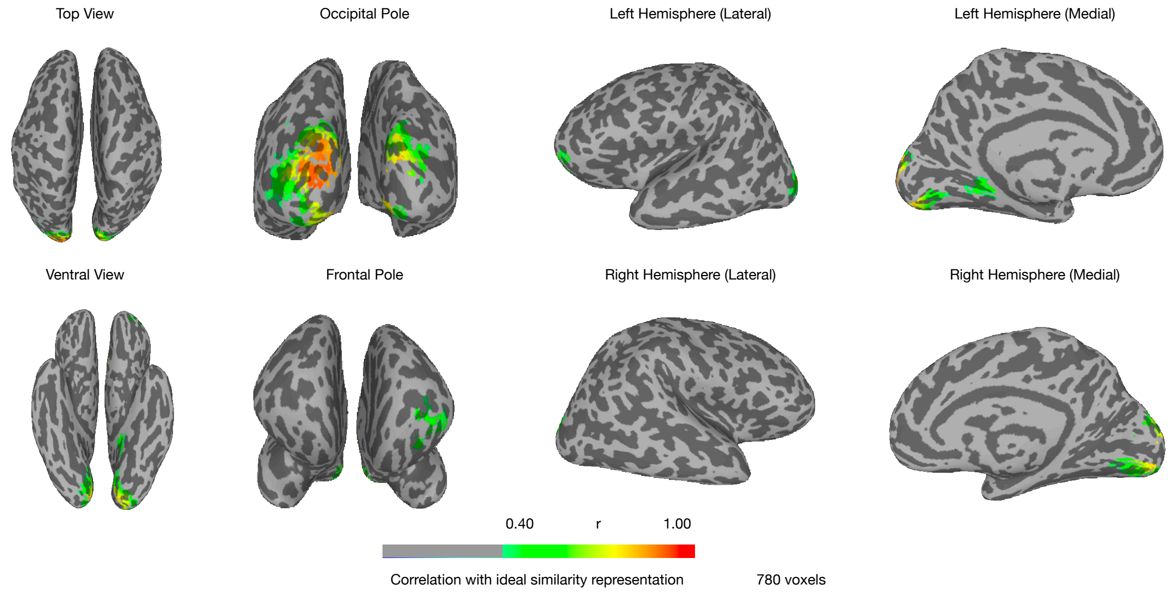


**Fig. S8. Neurofeedback ROI for participant #6.**

Searchlight analysis indicating all of participant #6's brain regions that represented the shape space parametrically, including extrastriate visual cortex and orbitofrontal cortex. See Methods for details of the searchlight procedure and threshold selection.

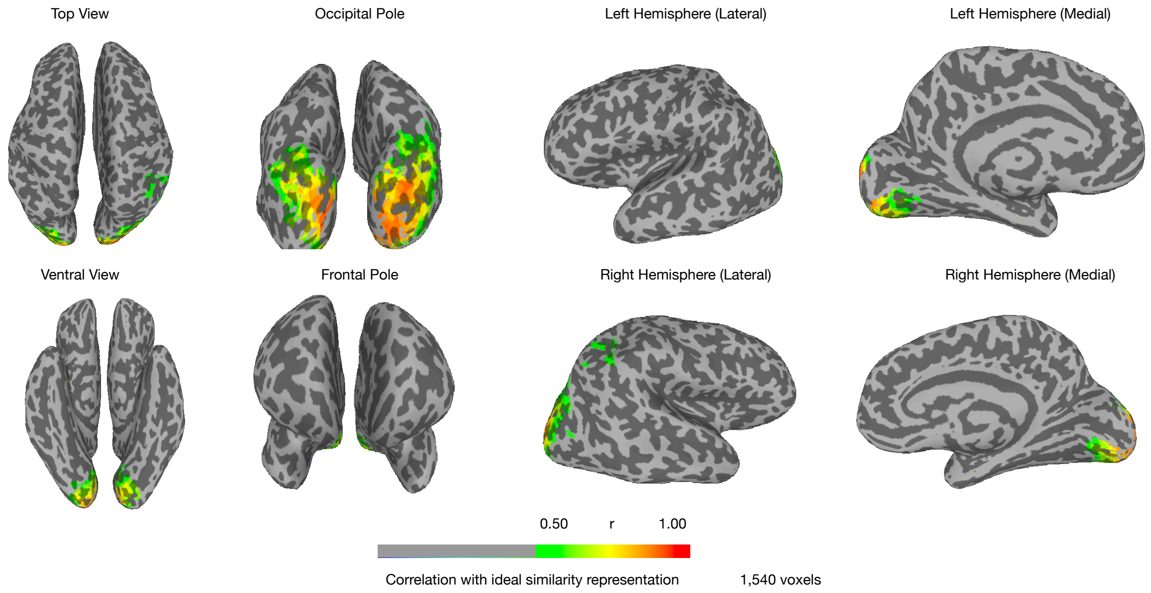


**Fig. S9. Neurofeedback ROI for participant #7.**

Searchlight analysis indicating all of participant #7's brain regions that represented the shape space parametrically, including extrastriate visual cortex and parietal cortex. See Methods for details of the searchlight procedure and threshold selection.

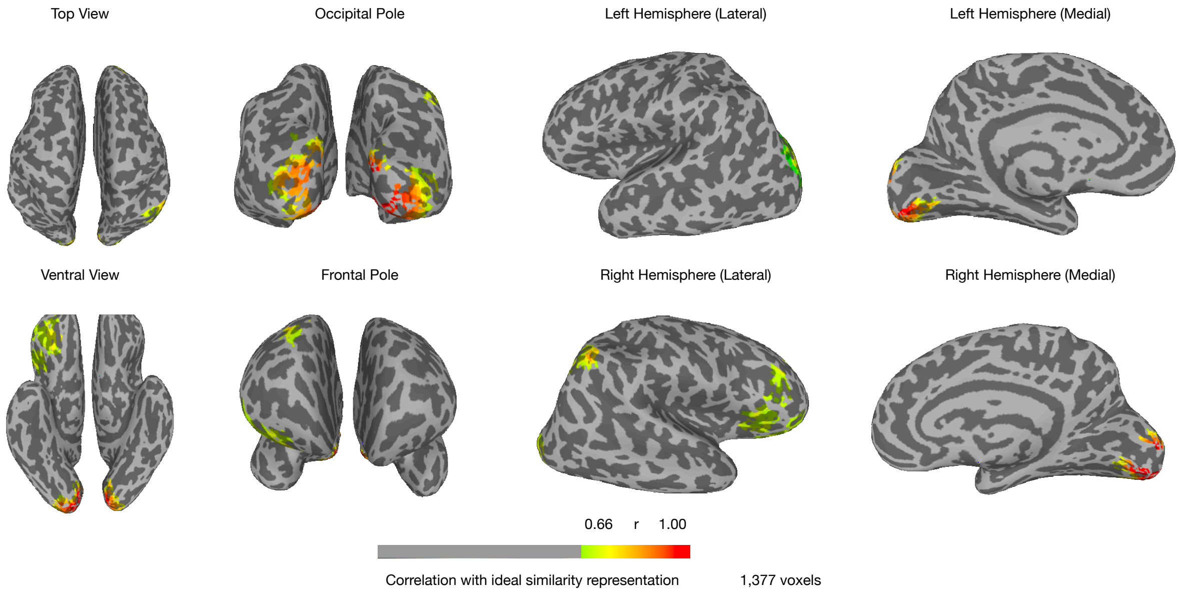


**Fig. S10. Neurofeedback ROI for participant #8.**

Searchlight analysis indicating all of participant #8's brain regions that represented the shape space parametrically, including extrastriate visual cortex, parietal cortex, and dorsolateral prefrontal cortex. See Methods for details of the searchlight procedure and threshold selection.

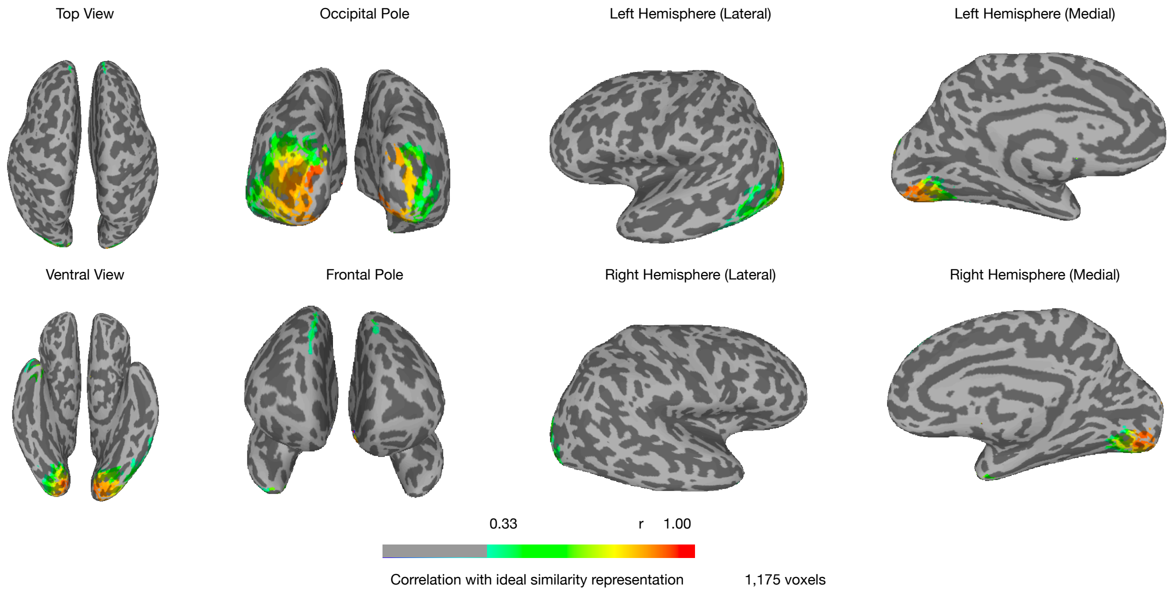


**Fig. S11. Neurofeedback ROI for participant #9.**

Searchlight analysis indicating all of participant #9's brain regions that represented the shape space parametrically, including extrastriate visual cortex, transverse occipital sulcus, inferior parietal cortex, and dorsolateral prefrontal cortex. See Methods for details of the searchlight procedure and threshold selection.

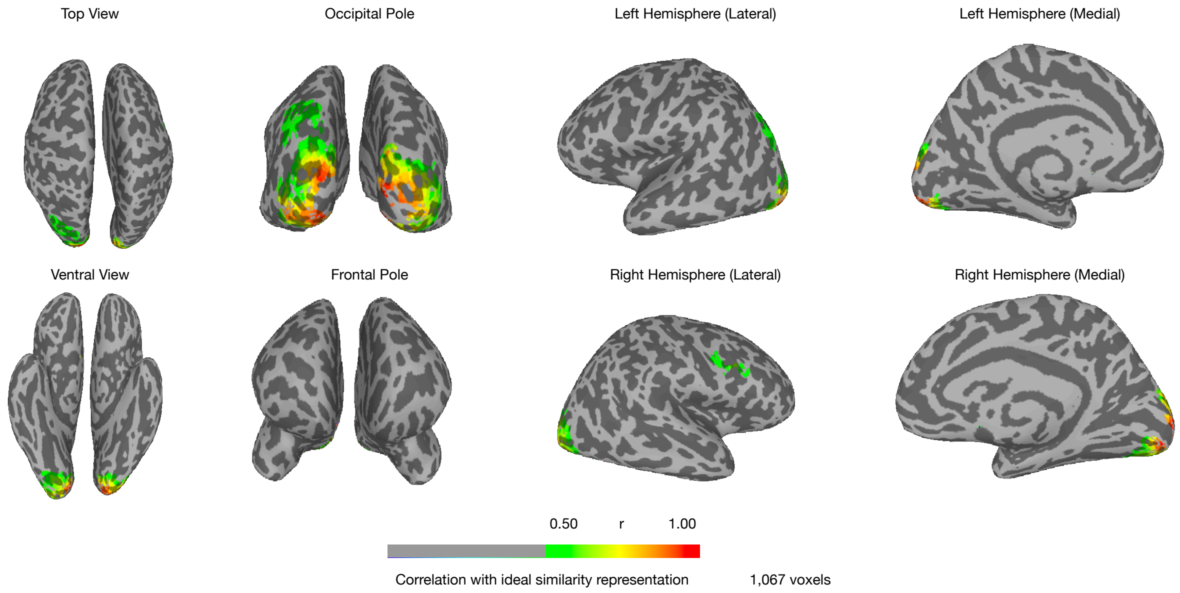


**Fig. S12. Neurofeedback ROI for participant #10.**

Searchlight analysis indicating all of participant #10's brain regions that represented the shape space parametrically, including extrastriate visual cortex, inferior temporal cortex, anterior temporal cortex, and dorsolateral prefrontal cortex. See Methods for details of the searchlight procedure and threshold selection.

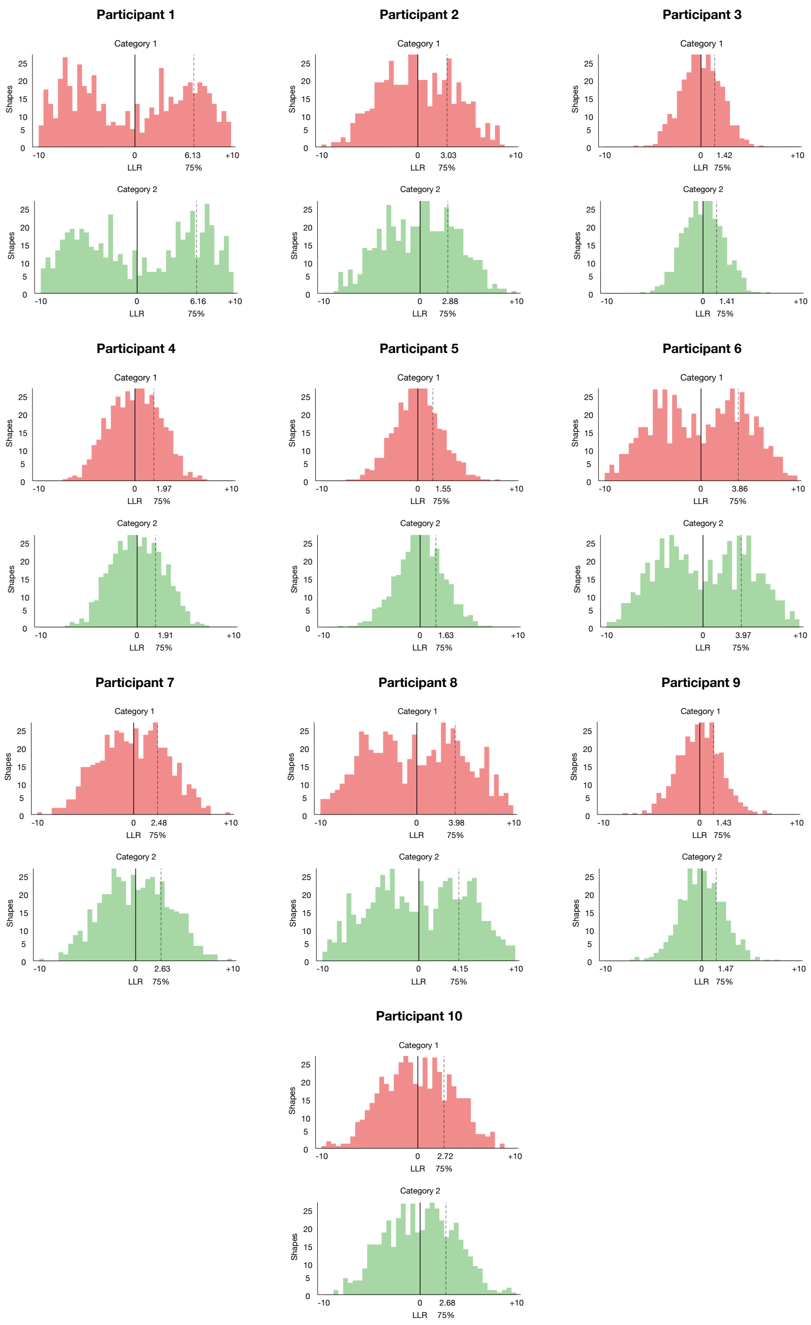


**Fig. S13. LLR histograms for shapes shown during localizer scans (days 2-3).**

We computed a distribution of log-likelihood ratios (LLR) using the estimated neural model Gaussian distributions for 48 out of the 49 shapes shown to participants (all except the ambiguous center shape) during each fMRI run scanned during the Days 2-3 fMRI sessions (12-15 runs per participant, 576-720 examples per category per participant). The full histogram of the LLR distributions are shown for each participant and each category (by definition, LLR1 = - LLR2), together with a representative threshold of 75% of the distribution used during the neurofeedback trials to assess performance and give feedback to the participants for each category.

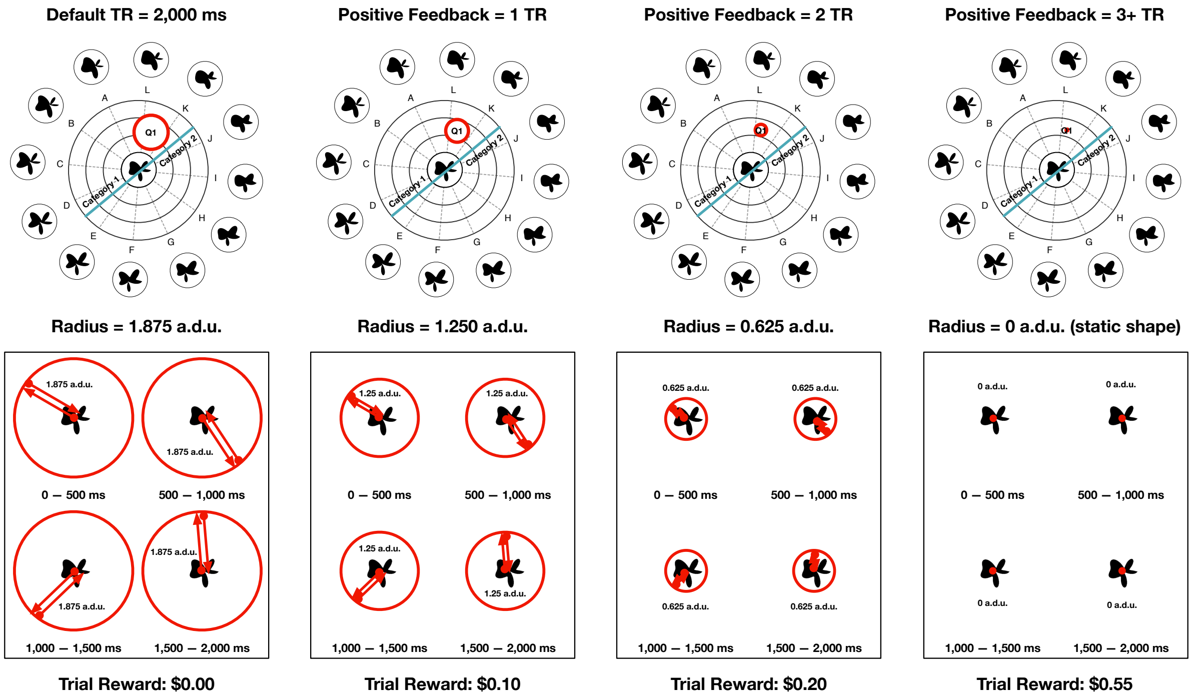


**Fig. S14. Neurofeedback trials with varying amounts of feedback.**

The four types of TRs shown to participants during the neurofeedback training. At the beginning of each trial, a random shape was selected and made to oscillate with a radius of 1.875 a.d.u. (default TR) (the shape position was static on the screen, but the shape identity morphed gradually according to the trajectory shown in parameter space). If a participant accumulated positive feedback within that trial (cf. Fig. 3D), the radius of the oscillation was reduced in increments of 0.625 a.d.u., otherwise it was left unchanged from the previous TR. Each 2s TR comprised four independent shape oscillations of 500 ms each (see Movie S1 for two example trials). Participants also received a monetary reward for each trial in which they were able to reduce the radius of the oscillation.

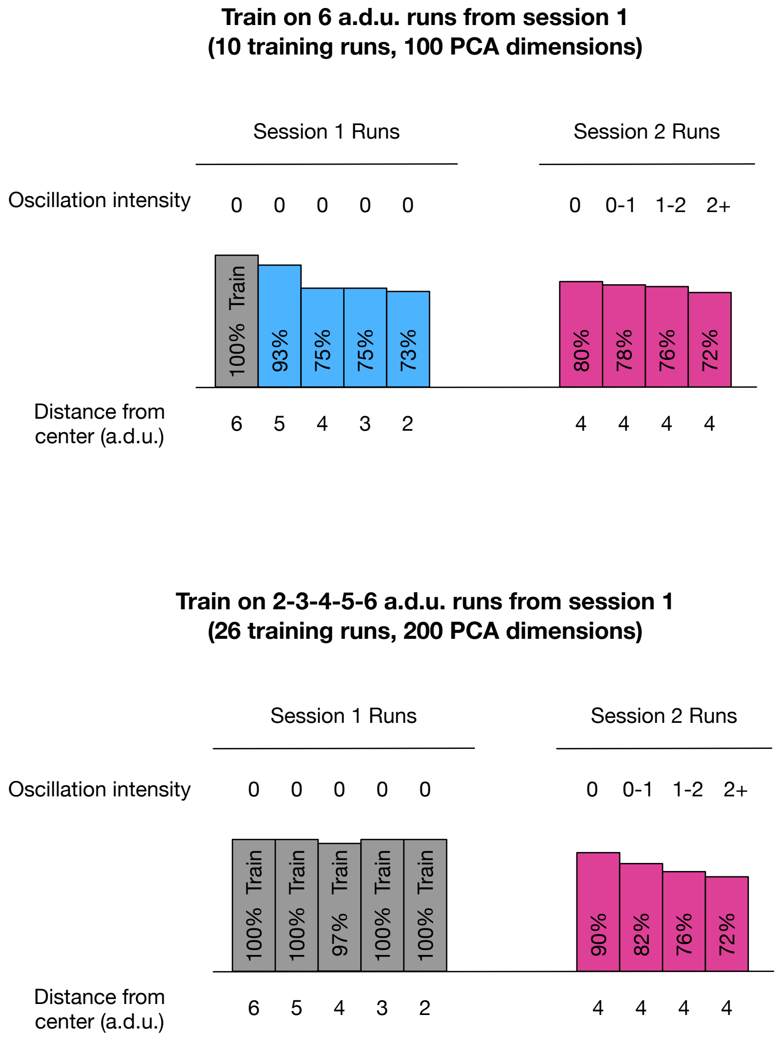


**Fig. S15. Gaussian model decoding performance decays closer to the category boundary and with increased oscillation radius.**

We collected two fMRI sessions for a pilot participant who performed the localizer task during the first session (four scans on separate days, 26 runs, 20 shapes of varying distance from center per run) and a center dot color change task during the fifth session (8 runs of 20 shapes similar to neurofeedback design, of varying levels of oscillation radius). Gray bars show classifier training accuracy. Blue bars show classifier testing accuracy for runs with varying distance from stimulus space center in session 1. Pink bars show classifier testing accuracy for runs with varying oscillation radius from session 2. Performance of the model decayed for shapes closer to the category boundary (top left) and when the model was trained on static shapes (e.g., localizer runs) and tested on oscillating shapes (e.g., neurofeedback runs). Based on these data, we chose an upper limit of 1.875 a.d.u. for the shape oscillation during neurofeedback trials and did not train shapes that were less than 1 a.d.u. from the category boundary, as the performance of the model was not reliable for combinations of parameters outside of these bounds.

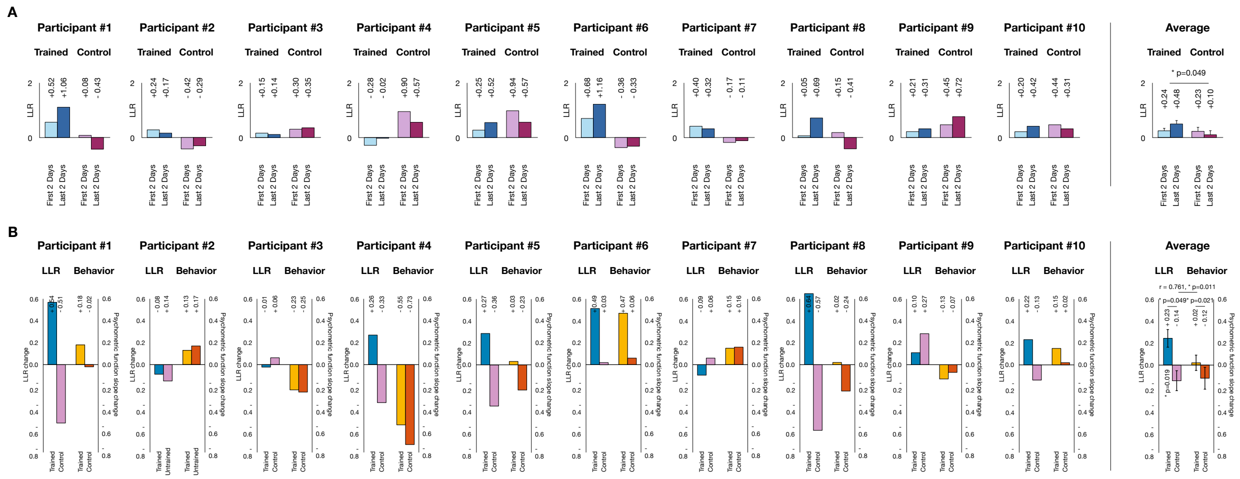


**Fig. S16. Neural and perceptual effects of neural sculpting for individual participants.**

(A) LLR for each participant's trained (blue) and control (purple) categories during the first two days (light color) and last two days (dark color). We observed a significant interaction effect between training and category (ANOVA; $F(1,76)=4.92$, $p=0.030$); (B) LLR for the trained (blue) and control (purple) categories, plotted together with the psychometric function changes for the trained (light orange) and control (dark orange) categories. Difference between LLR bars yields graph in Fig. 4D and difference between Behavior bars yields graph in Fig. 4E.

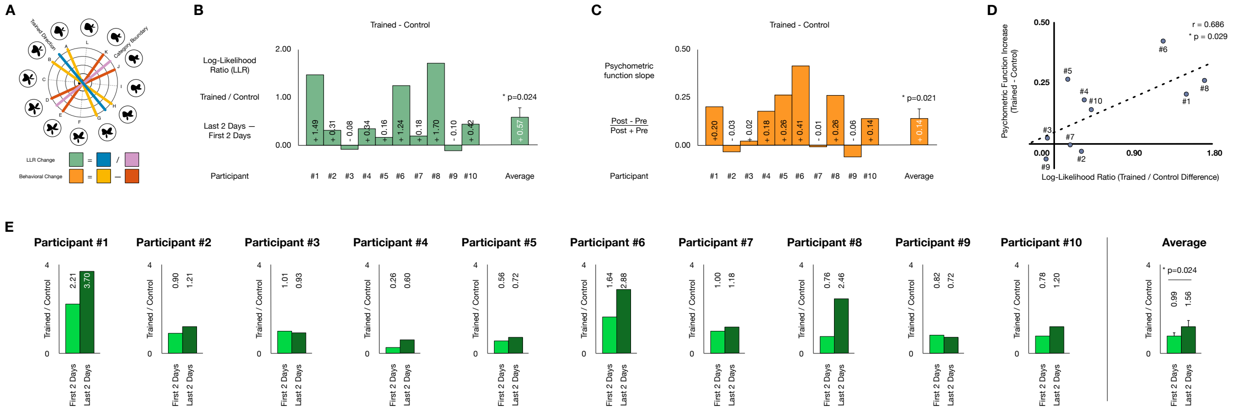

**Fig. S17. Neural sculpting effects and correlation with behavior using LLR ratio are similar to those using a difference score.**

(A) Diagram of stimulus space for example participant with emphasized trained direction, category boundary (for LLR computation), and directions used to compute psychometric function slope changes in the behavioral experiments. LLR Ratio (green) represents the LLR in the Trained direction divided by the LLR in the control direction. (B) Main effect of LLR Ratio change as a consequence of neurofeedback training (LLR Ratio difference between first two days and last two days of training). (C) Main behavioral effect of neurofeedback training (replicated from Fig. 4E for convenience). (D) Correlation plot of LLR ratio main effect vs. behavioral main effect. Similar to the LLR difference score measure, we observed a strong correlation between the neural change and perceptual change (r=0.686, p=0.029). (E) LLR Ratio for each participant during their first two days and last two days of neurofeedback training. We observed a significant increase of LLR ratio at the end of the experiment compared to the beginning (p=0.024).

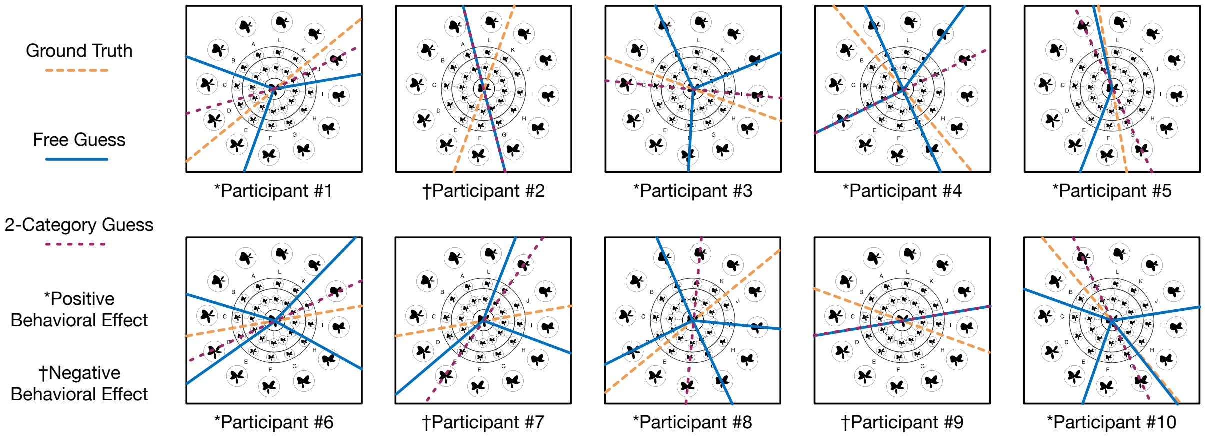


**Fig. S18. Category boundary guesses before final debrief.**

At the conclusion of the experiment (but before being fully debriefed), participants were told that the shapes they had seen during training belonged to an unspecified number of categories. They were instructed to draw their best guess for the category boundary/boundaries between them. Participants reported an average of 3.3 categories, whose boundaries (blue lines) did not coincide with the category boundaries randomly selected for each of them in the experiment (orange lines). When subsequently forced to split the stimulus space into 2 categories using a straight line, their performance suggested a trend towards being able to access some information about the category boundary given an explicit external constraint (purple line, trained=0°, average absolute displacement=31.5° ± 5.89° s.e.m,; random choice=45°, $t(9)=2.18$, $p=0.058$).

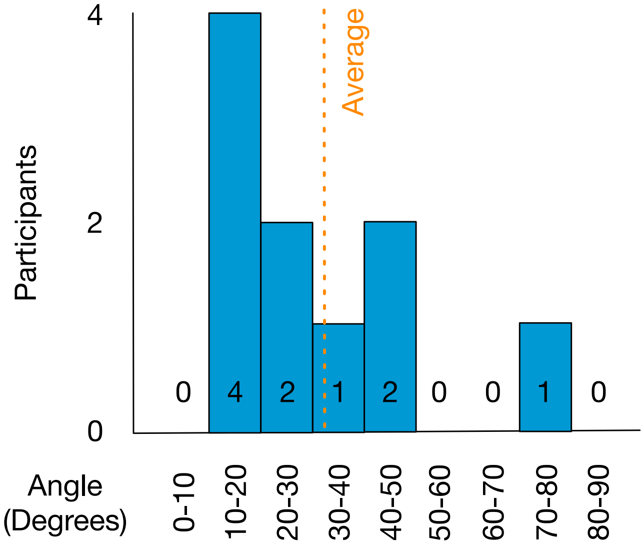


**Fig. S19. Histogram of forced 2-category boundary guesses.**

Histogram of angle displacement from correct category boundary for forced 2-category guesses in Fig. S18 (trained=0°, average absolute displacement=31.5° ± 5.89°; control=90°, average absolute displacement=58.5° ± 5.89°). Participants' performance suggests a trend towards having some crude information about the category boundary given an explicit external constraint (random choice=45°, $t(9)=2.18$, $p=0.058$).

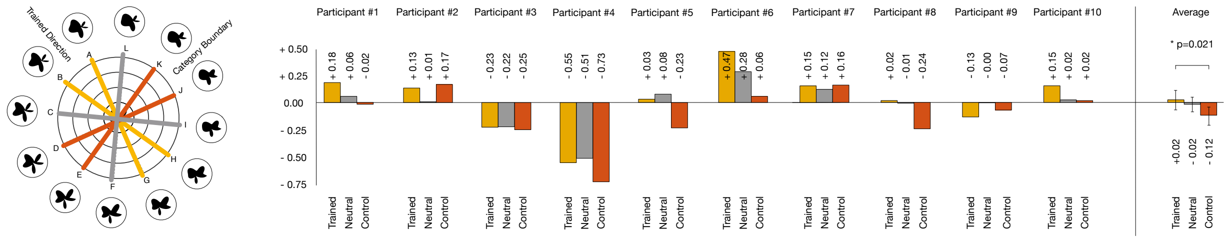


**Fig. S20. Neutral direction in shape space was not strongly affected by neurofeedback training.**

As a control analysis, we computed the psychometric function slope change for the average of the two directions that were midway between parallel and perpendicular to the trained category boundary (in gray, neutral, e.g., for categories ABCDEF and GHIJKL, the average slope of lines BH and EK). The behavioral change in the neutral direction was almost absent (-0.017±0.064 s.e.m., $t(9)=0.257$, $p=0.803$) and this change was not significantly different from that observed in the trained or control directions, albeit potentially more similar to the former than the latter (neutral<trained $t(9)=1.26$, $p=0.239$; neutral>control $t(9)=2.08$, $p=0.067$). The trained and control bar plots are replicated from Fig.S16B for convenience.

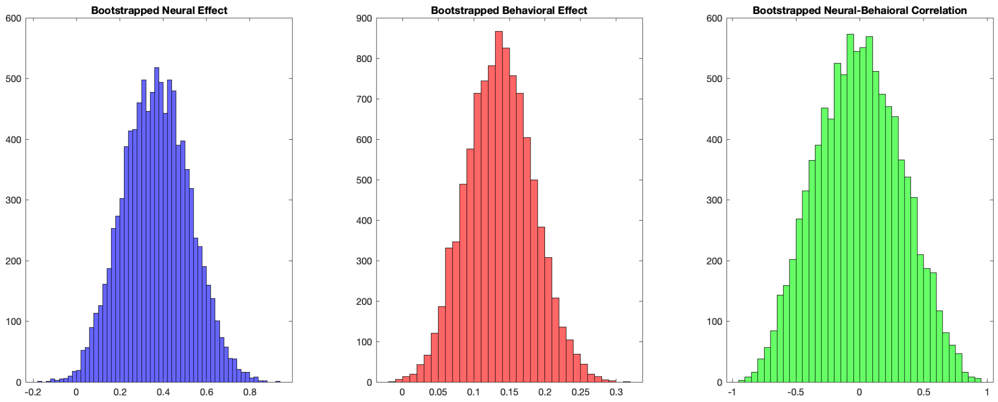


**Fig. S21. Bootstrap resampling analysis showed that our results were robust at the individual level.**

A sample-with-replacement procedure was used to generate 10,000 random draws of 10 samples (i.e., 10 participants) from the distributions of observed neural and behavioral effects (i.e., LLR difference values and psychometric function normalized difference scores reported in Fig. 4). The histograms of mean observed effects across all samples are shown in blue (neural LLR change), red (behavioral effect), and green (correlation values between each of the neural and behavioral samples). The probability of observing a positive neural effect by chance given our cohort is p(neural>0) = 0.005; the probability of observing a positive behavioral effect by chance given our cohort is p(behavioral>0) < 0.001; the probability of observing a correlation of equivalent magnitude to our reported effect (r=0.761) by chance given the distributions of neural and behavioral effects in our cohort is p(correlation observed>sampled correlation) = 0.006.

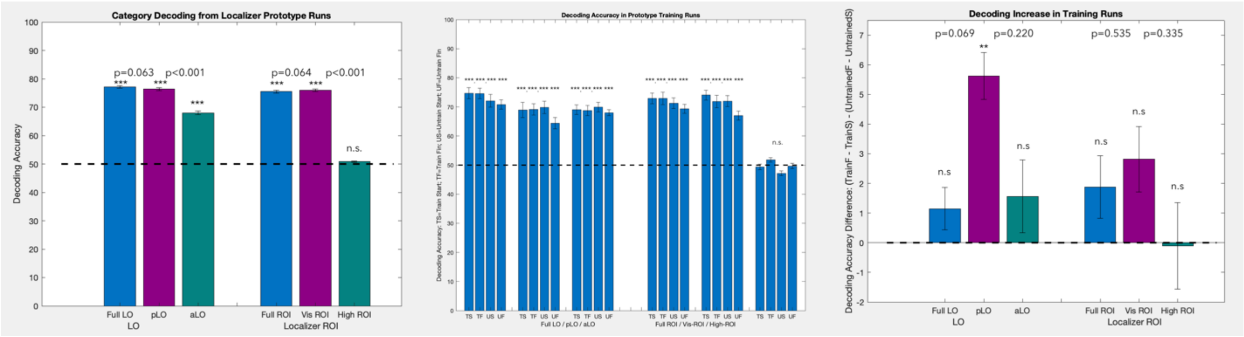


**Fig. S22. Split ROI analyses: anterior vs. posterior LO and visual vs. "high-level" ROI.** To measure the strength of category representations within sub-regions of our ROIs, we (1) split LO into equal sized anterior (aLO) and posterior (pLO) halves using the Z coordinate of the Freesurfer map from which the ROI was defined; and (2) split the neurofeedback ROI into "visual" (vis-ROI, all voxels within occipito-temporal cortex) and "high-level" (high-ROI, all remaining ROI voxels outside occipito-temporal cortex) components. We trained new category decoding models within these pairs of sub-ROIs for the runs collected during the two localizer scans (Days 2—3, 12-15 runs per participant). (Left) Decoding accuracy in aLO was markedly lower compared to pLO or to the full LO ROI (p<0.001). Furthermore, in the neurofeedback ROI, we could decode categories only in vis-ROI, but not in high-ROI. (Center) To quantify changes in decoding accuracy due to training, we computed the decoding accuracy of these models for the trained (T) and control (U) directions, across the calibration runs collected during first two days (TS, US) and the last two days (TF, UF) of training. In the trained direction, numerically, the decoding accuracy usually remained the same in all regions between the beginning and the end of training. In the control direction, however, accuracy numerically dropped after training, which is consistent with the main results, where we saw feature suppression in the control direction, rather than feature enhancement of the trained direction. While all accuracies were significantly higher than chance individually (except for high-ROI,) they were not significantly different from each other between TS vs. TF or US vs. UF. The only two exceptions were US vs. UF in pLO and in vis-ROI, which is consistent with the main results. (Right) We computed the difference of differences for the data shown in (Center). Numerically, we observed a trend for more posterior sub-regions to undergo more change due to training (i.e., pLO and vis-ROI). However, the only significant effect of decoding accuracy increase after training was observed in pLO (p=0.004) and no significant changes were present between any of the regions (although a trend, p=0.069 was observed between pLO and the full LO region).

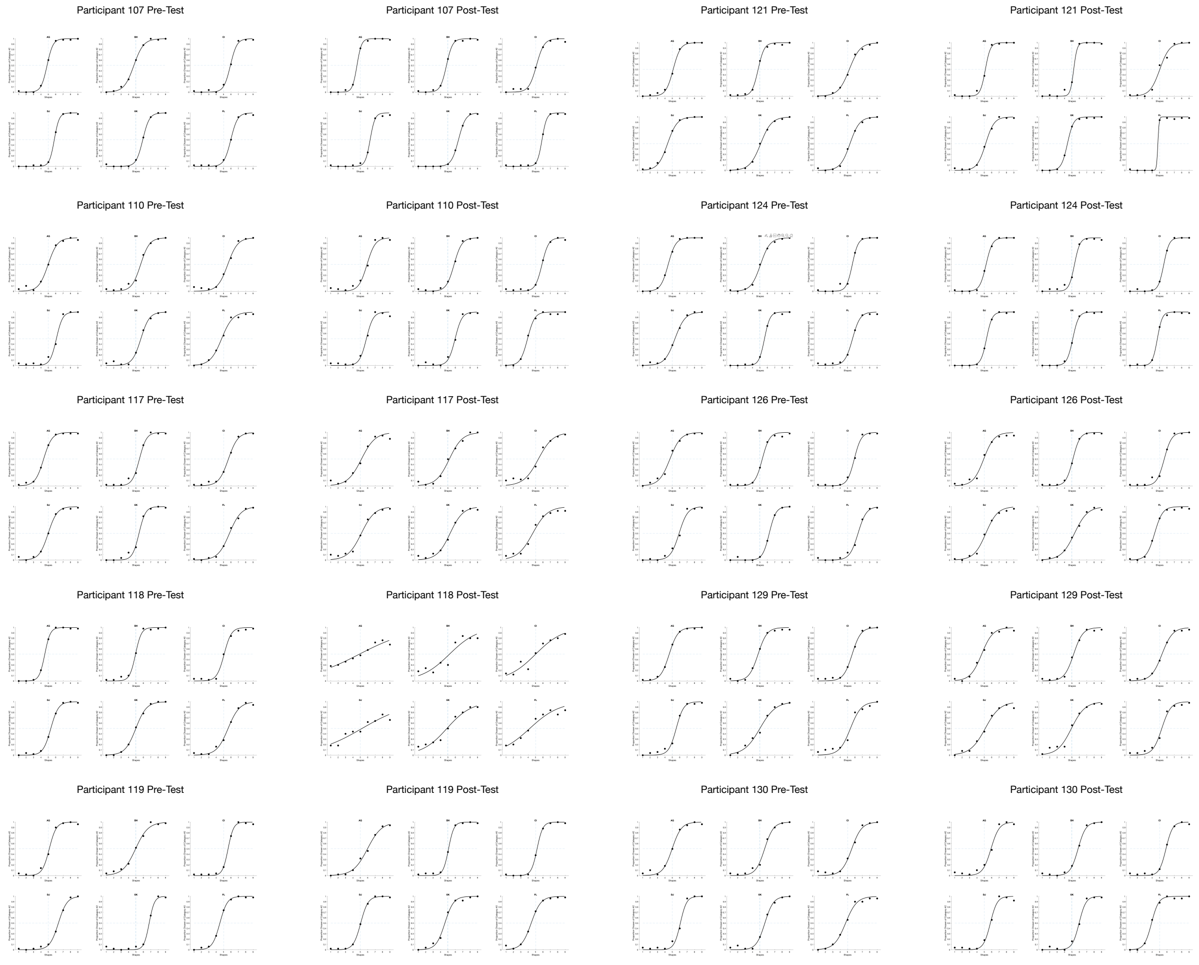


**Fig, S23. Psychometric functions for all participants and directions.**

To show that no categorization biases exist not just at the cohort level (Fig. S1), but also at the individual participant and direction level, we computed the psychometric functions for each individual direction tested in the shape space (AG, BH, CI, DJ, EK, FL) for each individual participant. No biases were observed for the trained of the control directions in the stimulus space at the individual level.

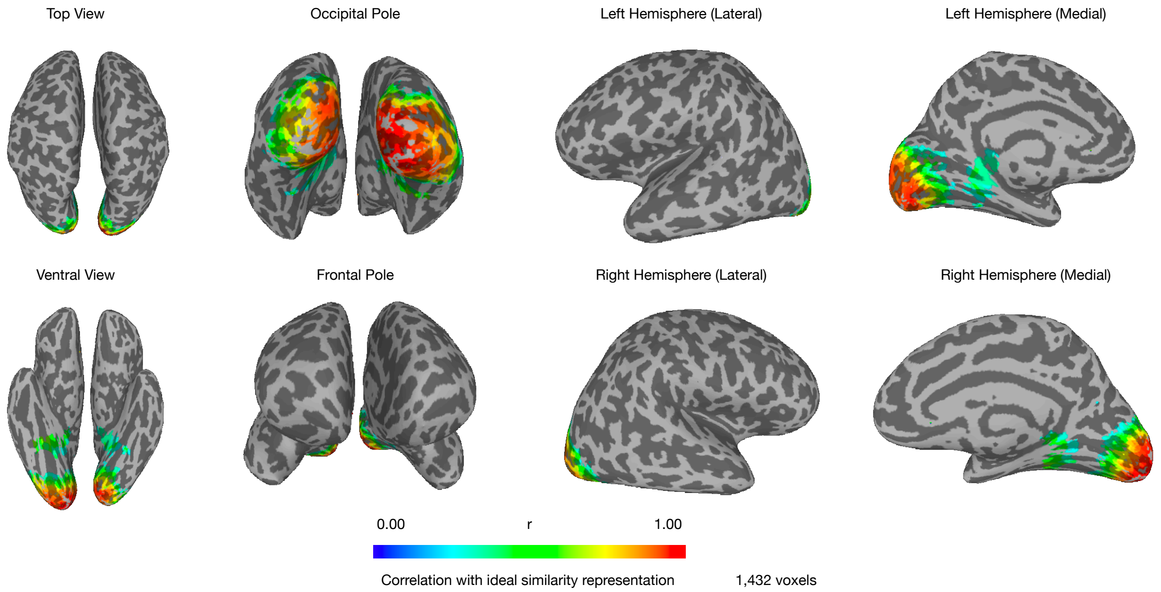


**Fig. S24. Neurofeedback ROI for participant #1 (including early visual cortex).**
Searchlight analysis indicating all of participant #1's brain regions that represented the shape space parametrically. See Methods for details of the searchlight procedure and threshold selection. (Fig. S3 shows the same brain regions, except without early visual cortex).

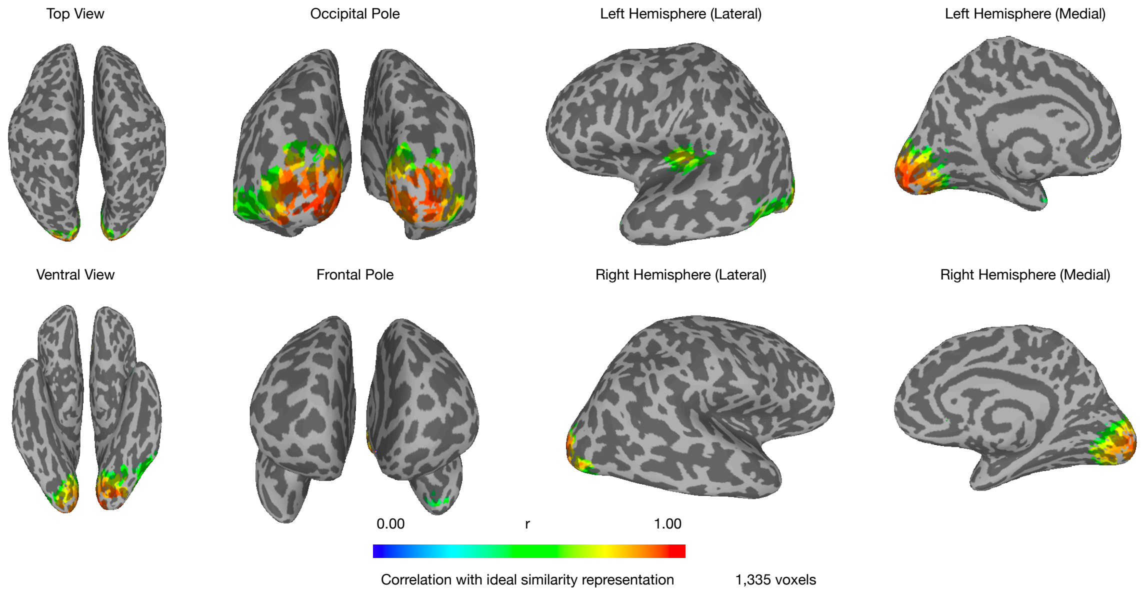


**Fig. S25. Neurofeedback ROI for participant #2 (including early visual cortex).**
Searchlight analysis indicating all of participant #2's brain regions that represented the shape space parametrically. See Methods for details of the searchlight procedure and threshold selection. (Fig. S4 shows the same brain regions, except without early visual cortex).

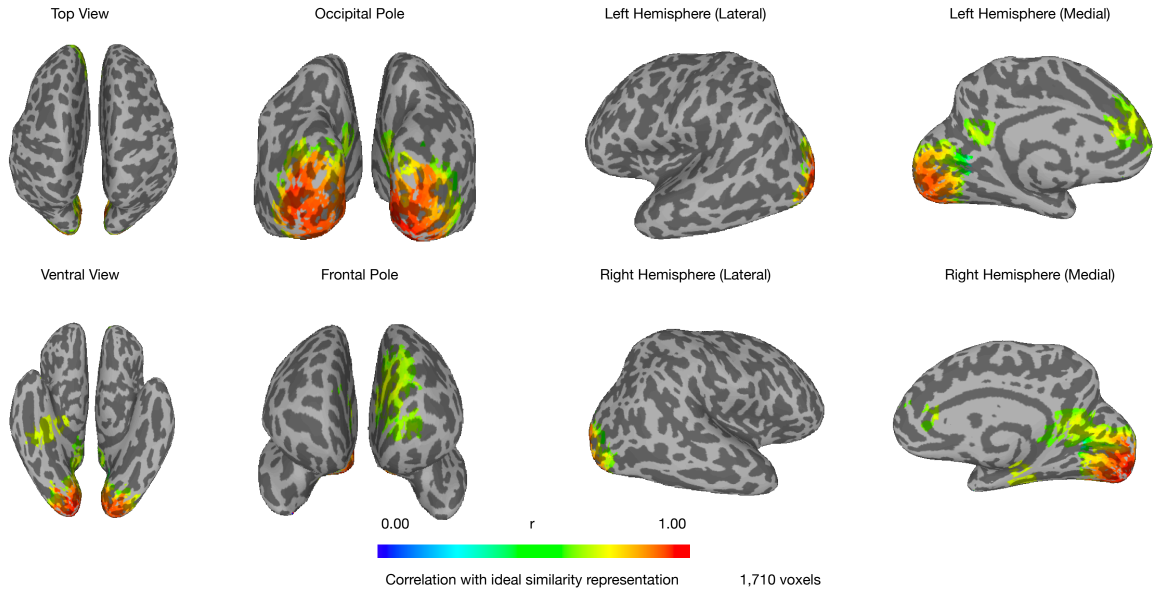


**Fig. S26. Neurofeedback ROI for participant #3 (including early visual cortex).**
Searchlight analysis indicating all of participant #3's brain regions that represented the shape space parametrically. See Methods for details of the searchlight procedure and threshold selection. (Fig. S5 shows the same brain regions, except without early visual cortex).

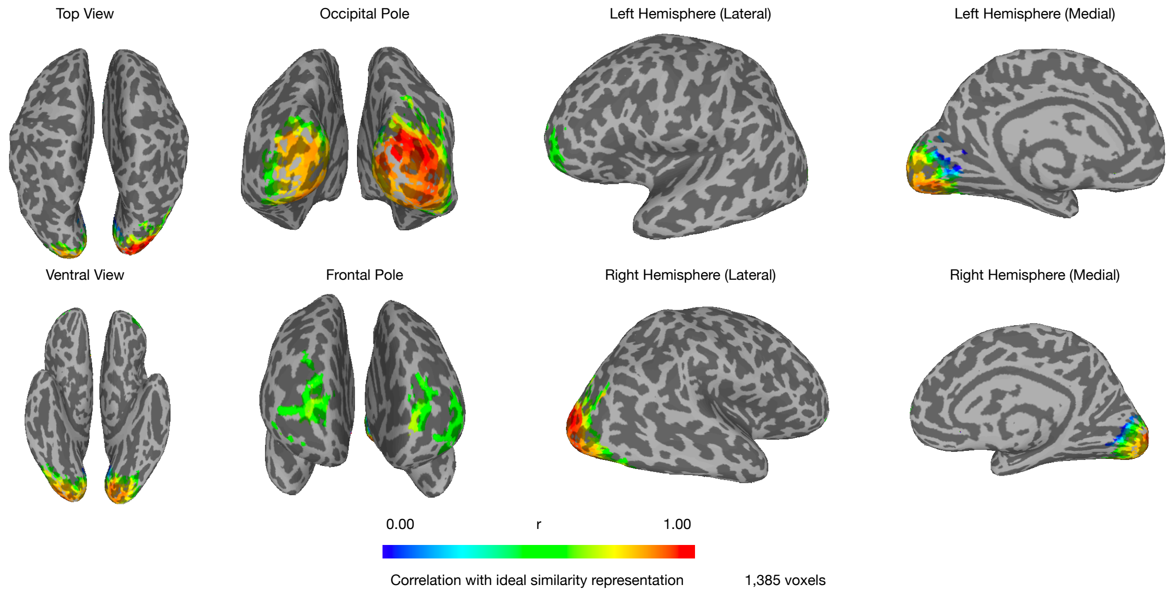


**Fig. S27. Neurofeedback ROI for participant #4 (including early visual cortex).**
Searchlight analysis indicating all of participant #4's brain regions that represented the shape space parametrically. See Methods for details of the searchlight procedure and threshold selection. (Fig. S6 shows the same brain regions, except without early visual cortex).

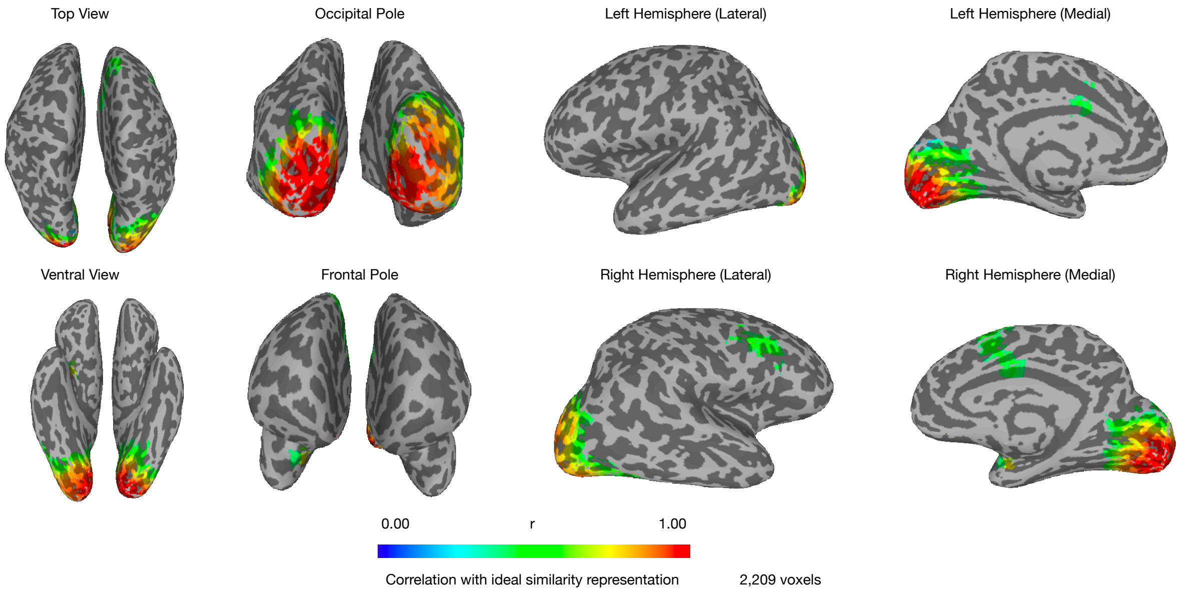


**Fig. S28. Neurofeedback ROI for participant #5 (including early visual cortex).**
Searchlight analysis indicating all of participant #5's brain regions that represented the shape space parametrically. See Methods for details of the searchlight procedure and threshold selection. (Fig. S7 shows the same brain regions, except without early visual cortex).

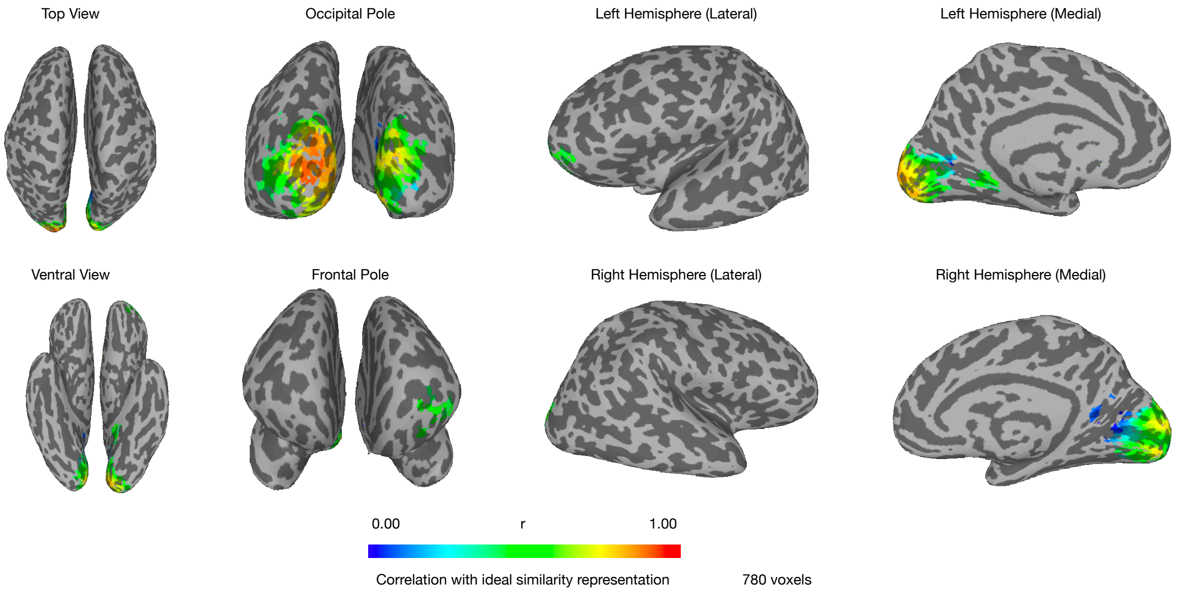


**Fig. S29. Neurofeedback ROI for participant #6 (including early visual cortex).**

Searchlight analysis indicating all of participant #6's brain regions that represented the shape space parametrically. See Methods for details of the searchlight procedure and threshold selection. (Fig. S8 shows the same brain regions, except without early visual cortex).

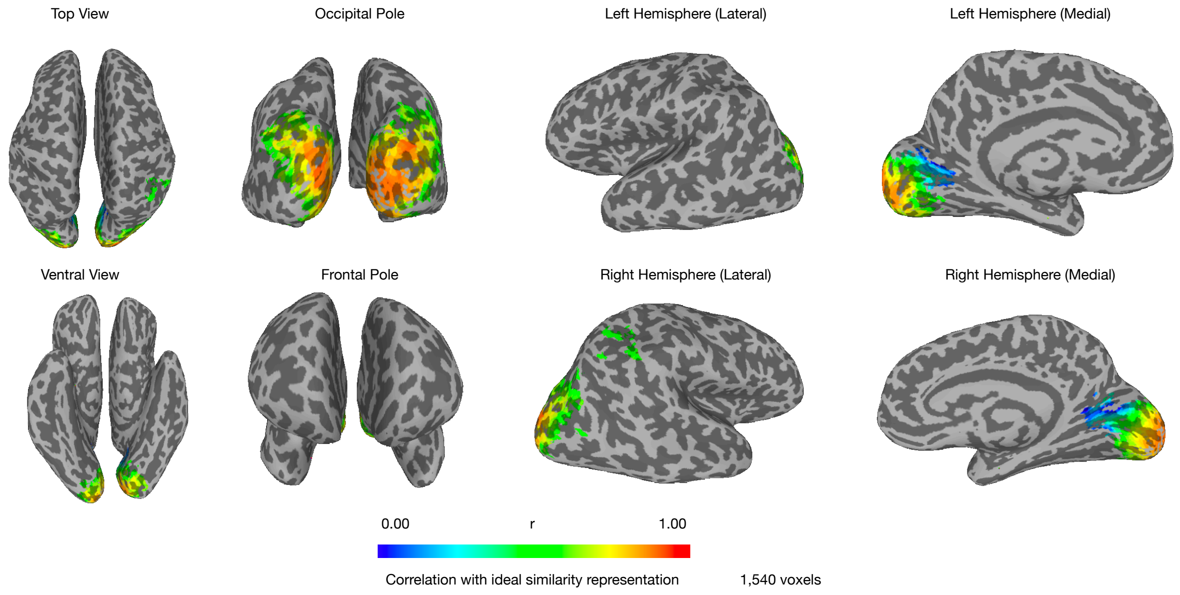


**Fig. S30. Neurofeedback ROI for participant #7 (including early visual cortex).**

Searchlight analysis indicating all of participant #7's brain regions that represented the shape space parametrically. See Methods for details of the searchlight procedure and threshold selection. (Fig. S9 shows the same brain regions, except without early visual cortex).

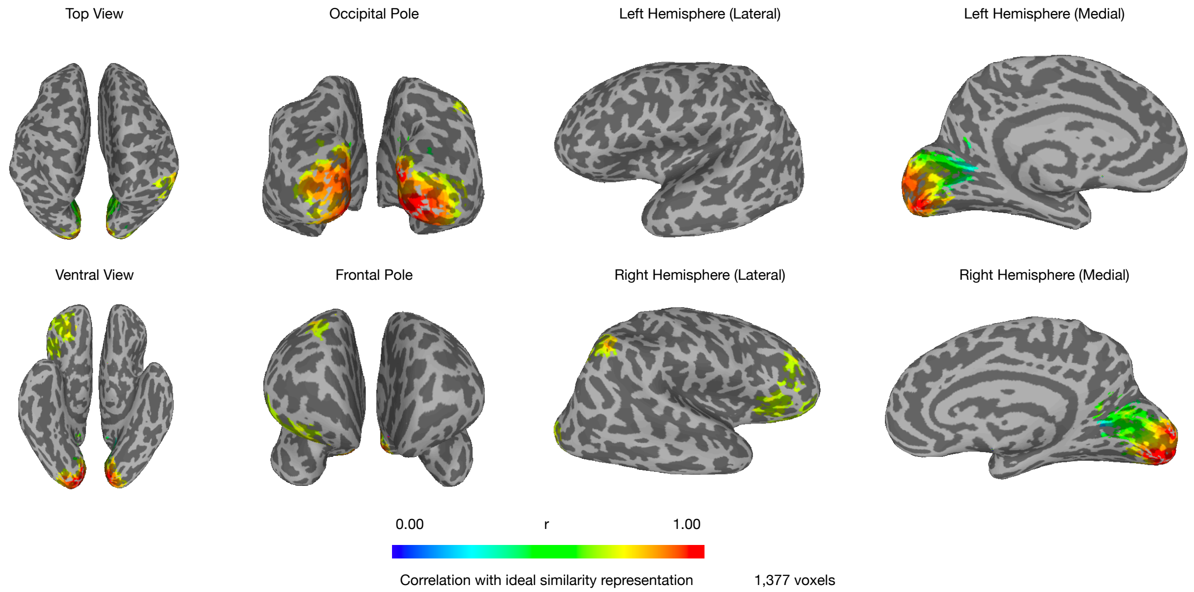


**Fig. S31. Neurofeedback ROI for participant #8 (including early visual cortex).**
Searchlight analysis indicating all of participant #8's brain regions that represented the shape space parametrically. See Methods for details of the searchlight procedure and threshold selection. (Fig. S10 shows the same brain regions, except without early visual cortex).

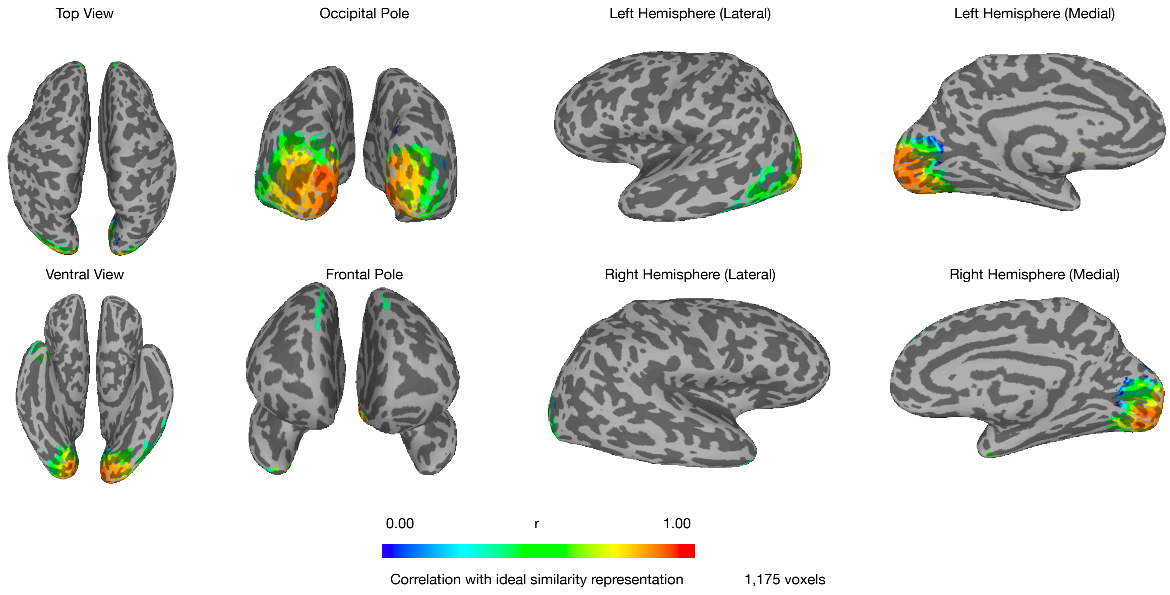


**Fig. S32. Neurofeedback ROI for participant #9 (including early visual cortex).**
Searchlight analysis indicating all of participant #9's brain regions that represented the shape space parametrically. See Methods for details of the searchlight procedure and threshold selection. (Fig. S11 shows the same brain regions, except without early visual cortex).

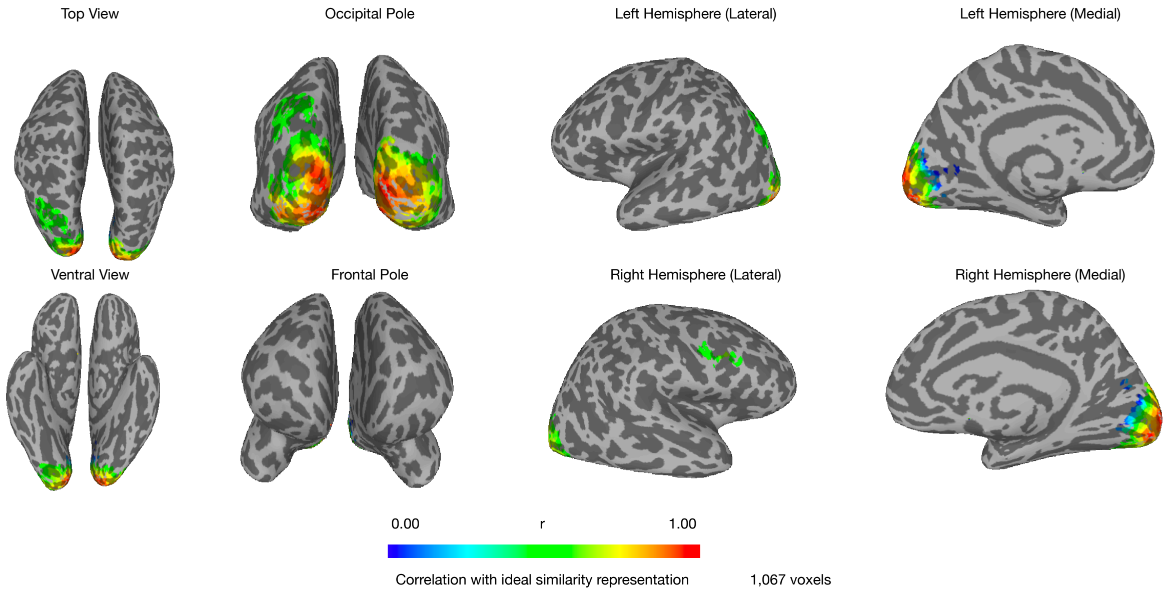


**Fig. S33. Neurofeedback ROI for participant #10 (including early visual cortex).**
Searchlight analysis indicating all of participant #10's brain regions that represented the shape space parametrically. See Methods for details of the searchlight procedure and threshold selection. (Fig. S12 shows the same brain regions, except without early visual cortex).

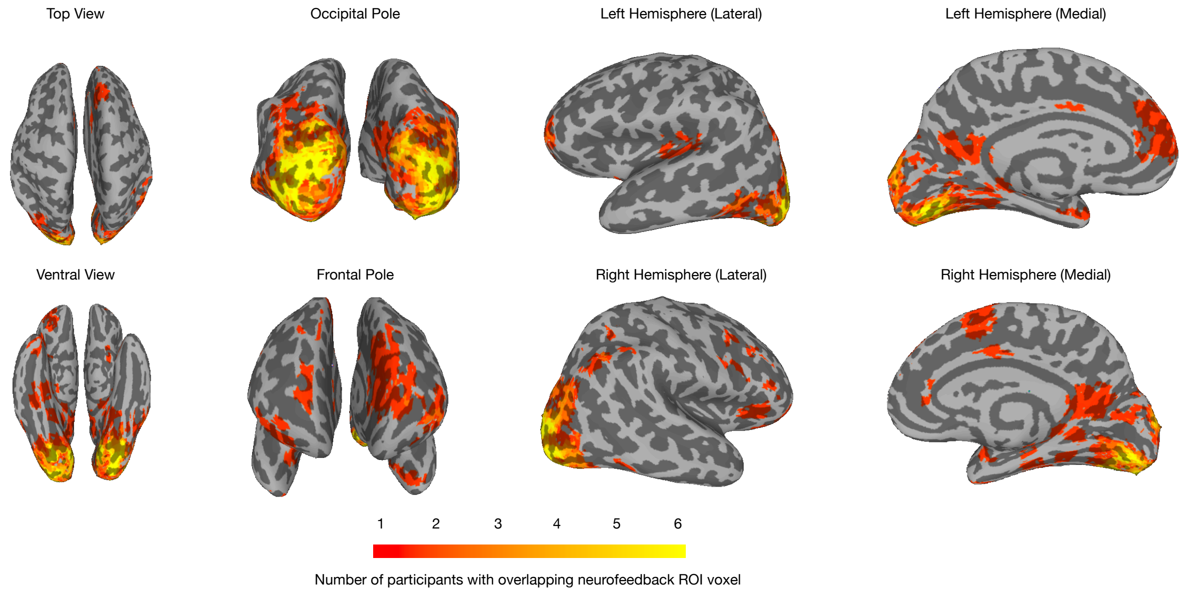


**Fig. S34. Group map for neurofeedback ROI.**

Group map of neurofeedback ROI overlap for all 10 participants (MNI registration of maps shown in Figs. S3—S12). A high degree of overlap was observed in extrastriate visual cortex (common ROI for up to 6 participants) and, to a smaller degree, in lateral temporal cortex, parahippocampal cortex, and prefrontal cortex (common ROI for up to 3 participants). We note that alignment to a common space is known to introduce biases in the spatial extent of maps, especially for highly specialized ROIs, such as the ones we defined based on an independent decoding accuracy criterion.

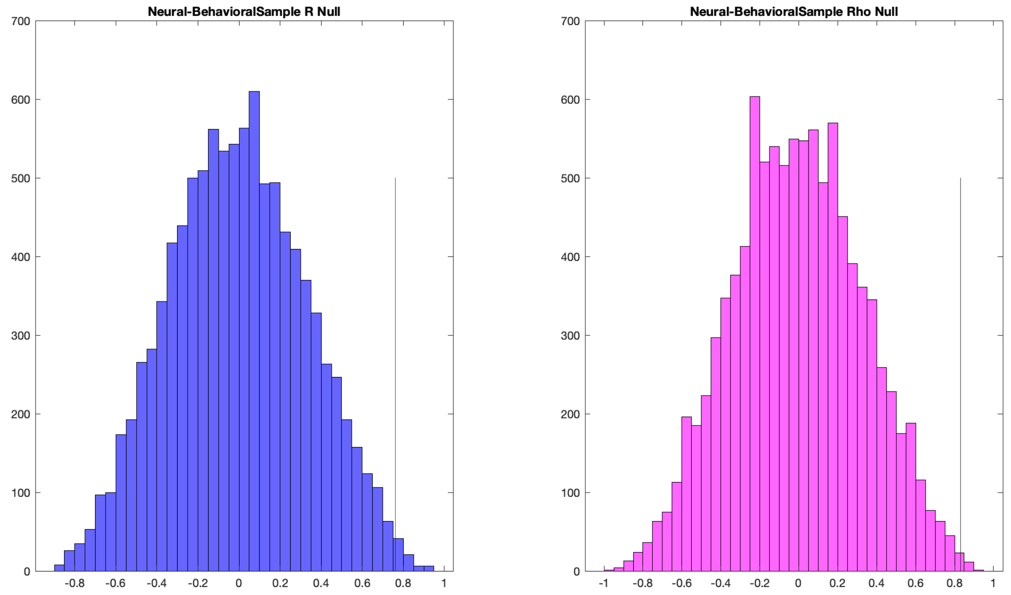


**Fig, S35. Null distributions for neural-behavioral correlations.**

We computed null distributions for the Pearson (left) and Spearman (right) correlation values corresponding to our main neural-behavior effect by keeping the neural effect values fixed and randomizing the order of the behavioral effect values 10,000 times. Both distributions had means close to zero (Pearson r mean<0.001, Spearman rho mean=0.001) and the correlation values observed for the unpermuted data were highly significant compared to these null distributions (Pearson r=0.761, p=0.006; Spearman rho=0.830, p=0.002; vertical lines in each graph).

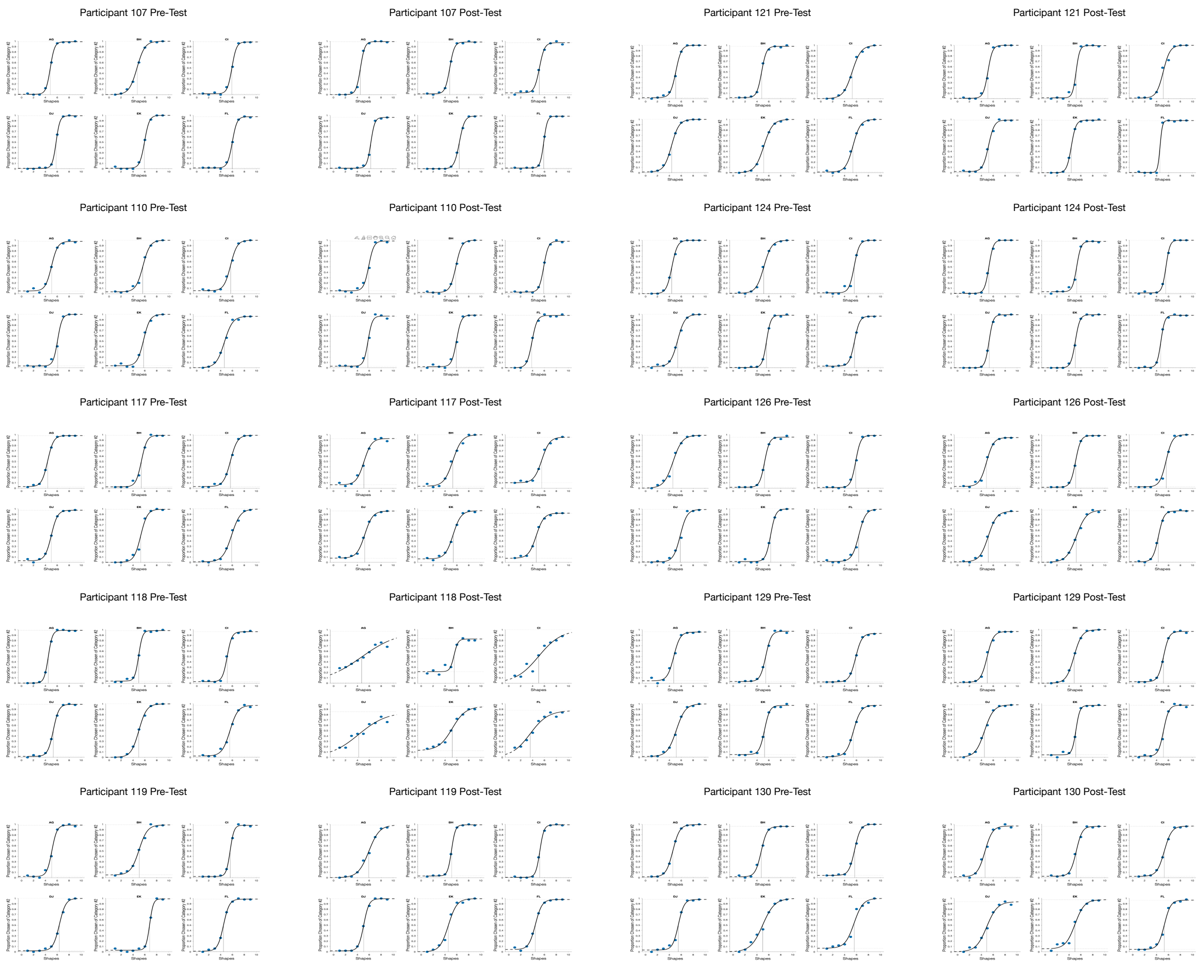


**Fig. S36. Alternate psychometric functions for all participants and directions.**

Psychometric function estimates for pre- and post-training behavioral tests that include explicit estimations of the lapses and thresholds (corresponding parameter estimates listed in Table S6).

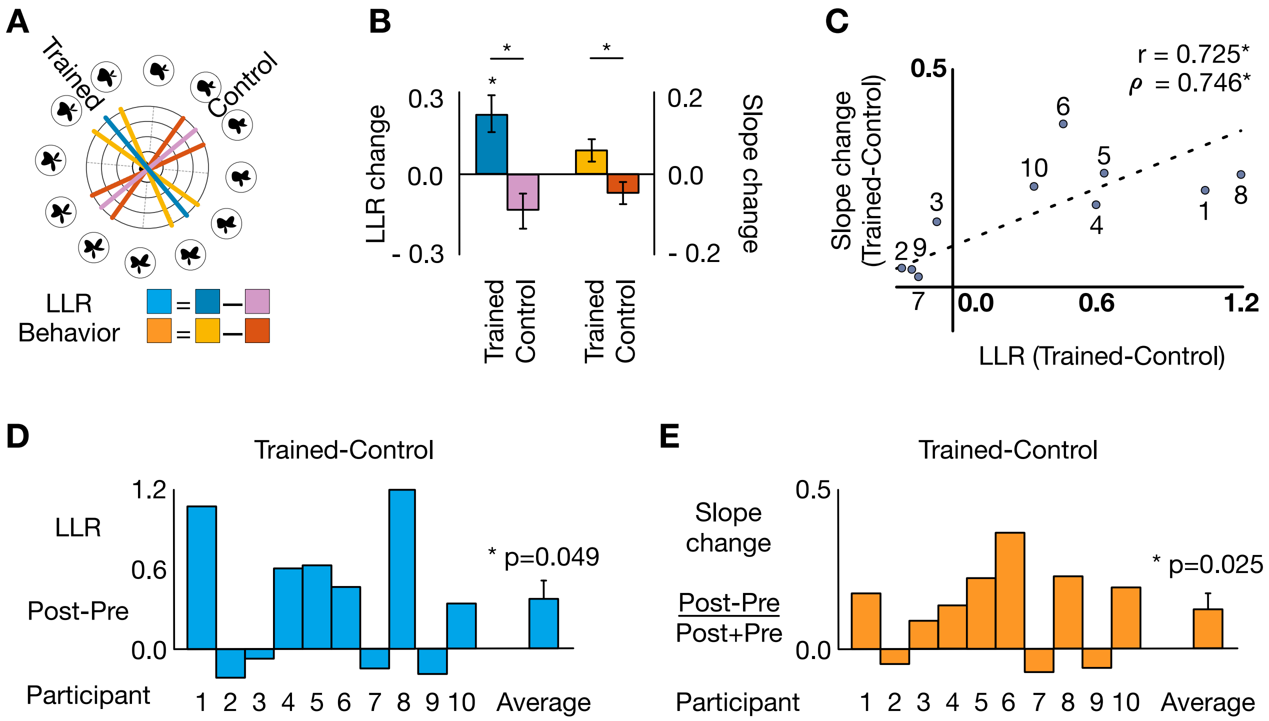


**Fig, S37. Behavioral effects and correlation with behavior using alternate psychometric function estimates are similar to original estimates (Fig. 4).**

(A) Shape space with example category boundaries. Diameters for trained category distinction: LLR = blue, psychometric function slope = average of yellow lines. Diameters for control category distinction: LLR = purple, psychometric function slope = average of red lines. (B) Effects of neural sculpting on neural representations and perception for trained and control categories: differences in LLR and psychometric function slopes (colors as in subpanel A; *p<0.05). (C) Changes in the brain due to neural sculpting predict perceptual changes (Pearson r, *p=0.018; Spearman rho, *p=0.018). (D) Change in LLR between the last two days and the first two days of training for individual participants. Positive values indicate stronger neural boundaries in trained vs. control categories. (E) Change in psychometric function slope for trained vs. control categories between behavioral pre- and post-tests for individual participants. Positive values indicate stronger categorical perception for trained versus control categories.

.

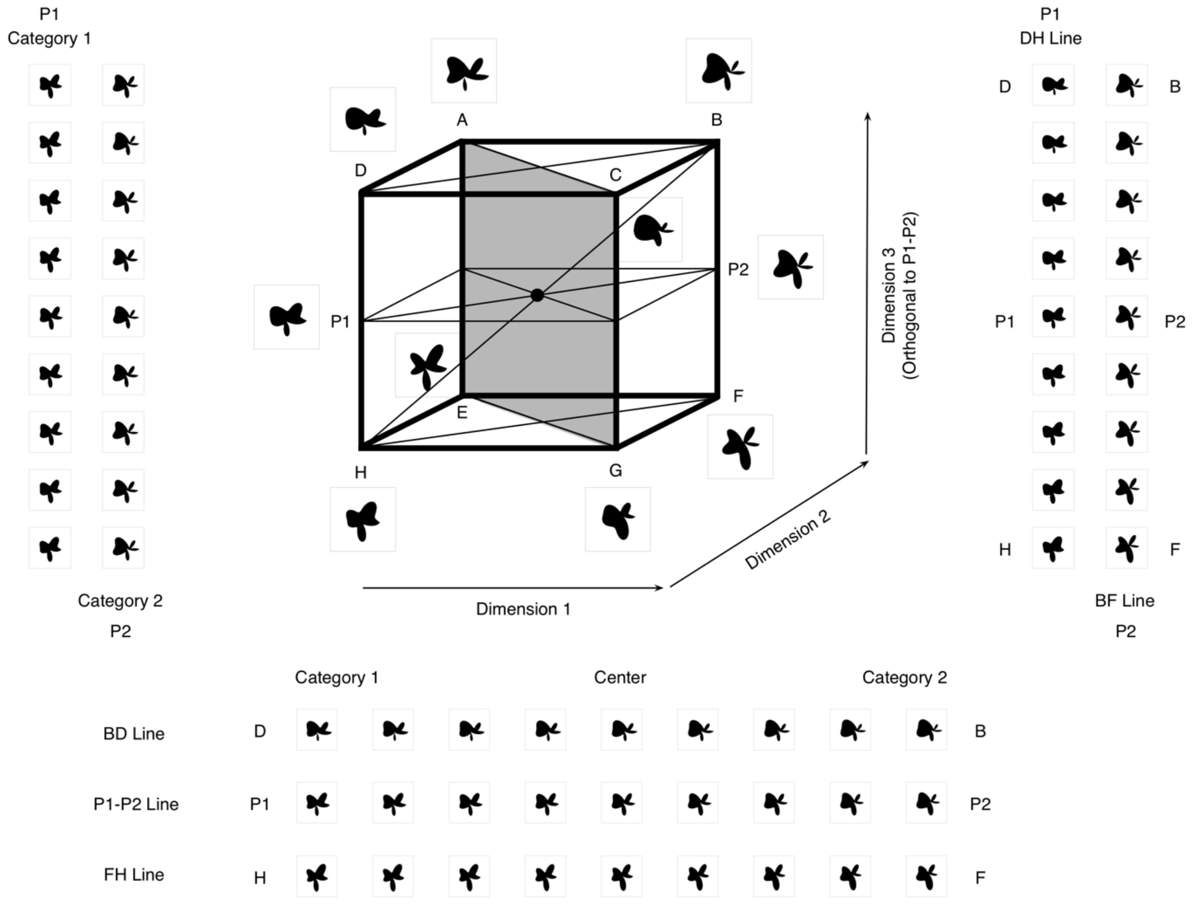


**Fig, S38. Preliminary shape space manifold: varying 3 dimensions out of a possible 7.**

Our initial pilot experiment selected 3 of 7 possible dimensions of variation in the shape space based on prior work (Op de Beeck et al., 2001; Kok et al., 2018) to measure how each of them, alone, or in combination, were categorically perceived by participants. This yielded a shape space that spanned this 3-dimensional cube in parameter space. We then ran versions of our 2AFC experiments for 7 lines in this 3-dimensional space and selected the shaded 2-dimensional manifold in the cube space for which the parametric midpoint between shape endpoints matched participants' performance most closely.

| | STATUS | PARTICIPANT ID | LOCALIZER RUNS | LATERAL OCCIPITAL ROI AVG ACCURACY | BEST PARAMS | ACCURACY % PER PARTITION | SEARCHLIGHT COGNITIVE MAP ROI R THRESHOLD | # VOXELS | AVG ACCURACY | BEST PARAMS | ACCURACY % PER PARTITION | TRAINED PARAMS | PARTITION | NOTES |
|---|---|---|---|---|---|---|---|---|---|---|---|---|---|---|
| 1 | REJECTED (PERF) | **106** | 13 | 67% | LAG 6, DIM 100 | 63-63-70-74-68-62 | | | | | | | | Rejected due to poor performance |
| 2 | FINISHED 30/06/2018 | **107** | 12 | 78% | LAG 6, DIM 100 | 78-79-78-77-77-81 | 0.25 | 1,432 | 76% | LAG 6, DIM 150 | 75-77-73-75-**76**-77 | LAG 6, DIM 150 | EFGHIJ vs. KLABCD | Finished! |
| 3 | REJECTED (PERF) | **108** | 12 | 56% | LAG 6, DIM 150 | 56-56-55-54-57-55 | | | | | | | | Rejected due to poor performance |
| 4 | WITHDRAWN (3) / REJECTED (PERF) | **109** | 14 | 73% | LAG 6, DIM 100 | 66-73-77-65-81-72 | 0.33 | 783 | 68% | LAG 6, DIM 150 | 65-69-72-63-77-64 | | | Quit right before could be rejected due to poor performance |
| 5 | FINISHED 17/10/2018 | **110** | 14 | 79% | LAG 4, DIM 200 | 75-81-81-71-88-79 | 0.40 | 1,335 | 74% | LAG 4, DIM 200 | 68-77-77-67-85-**70** | LAG 4, DIM 200 | FGHIJK vs. LABCDE | Finished! |
| 6 | WITHDRAWN (3) | **111** | 12 | 74% | LAG 6, DIM 150 | 74-73-76-76-74-73 | 0.50 | 1,755 | 70% | LAG 6, DIM 150 | 72-69-68-69-71-72 | | | Scheduled for training, then cancelled x2 |
| 7 | WITHDRAWN (3) | **112** | 13 | 75% | LAG 6, DIM 150 | 71-74-78-79-76-72 | 0.50 | 954 | 72% | LAG 6, DIM 200 | 69-67-73-78-75-67 | | | No email response when invited to return for training |
| 8 | WITHDRAWN (1) | **113** | | | | | | | | | | | | Quit after behavior pre-test |
| 9 | REJECTED (MRI) | **114** | | | | | | | | | | | | Eye surgery revealed when filling out screening form 3rd time |
| 10 | WITHDRAWN (2) | **115** | | | | | | | | | | | | Quit during 3rd run of first localizer scan |
| 11 | REJECTED (MRI) | **116** | | | | | | | | | | | | MRI-unsafe device revealed when filling out screening form 2nd time |
| 12 | FINISHED 26/03/2019 | **117** | 14 | 73% | LAG 6, DIM 100 | 66-73-79-63-78-77 | 0.60 | 1,710 | 71% | LAG 6, DIM 100 | 66-70-**78**-61-76-74 | LAG 6, DIM 100 | CDEFGH vs. IJKLAB | Finished! |
| 13 | FINISHED 30/04/2019 | **118** | 15 | 80% | LAG 4, DIM 150 | 76-85-74-83-76-85 | 0.50 | 1,385 | 79% | LAG 4, DIM 150 | 75-**85**-73-83-73-85 | LAG 4, DIM 150 | BCDEFG vs. HIJKLA | Finished! |
| 14 | FINISHED 27/06/2019 | **119** | 15 | 76% | LAG 6, DIM 150 | 72-80-68-83-67-83 | 0.40 | 2,209 | 76% | LAG 4, DIM 100 | **73**-81-70-81-70-83 | LAG 4, DIM 100 | ABCDEF vs. GHIJKL | Finished! |
| 15 | REJECTED (PERF) | **120** | 14 | 66% | LAG 6, DIM 200 | 62-66-70-62-69-67 | | | | | | | | Rejected due to poor performance (fell asleep during localizer scans) |
| 16 | FINISHED 20/07/2019 | **121** | 15 | 80% | LAG 6, DIM 150 | 75-88-75-84-75-83 | 0.40 | 780 | 79% | LAG 6, DIM 100 | 74-88-71-**85**-70-85 | LAG 6, DIM 100 | DEFGHI vs. JKLABC | Finished! |
| 17 | REJECTED (PERF) | **122** | 13 | 70% | LAG 6, DIM 100 | 68-63-71-78-74-65 | 0.33 | 1,444 | 67% | LAG 4, DIM 150 | 63-61-71-75-70-63 | | | Rejected due to poor performance |
| 18 | WITHDRAWN (1) | **123** | | | | | | | | | | | | Quit after behavior pre-test |
| 19 | FINISHED 29/08/2019 | **124** | 15 | 75% | LAG 4, DIM 150 | 71-79-70-79-70-80 | 0.50 | 1,540 | 73% | LAG 4, DIM 200 | 69-76-70-**76**-68-76 | LAG 4, DIM 200 | DEFGHI vs. JKLABC | Finished! |
| 20 | REJECTED (PERF) | **125** | 14 | 57% | LAG 4, DIM 150 | 52-59-60-54-61-54 | | | | | | | | Rejected due to poor performance |
| 21 | FINISHED 04/09/2019 | **126** | 14 | 81% | LAG 6, DIM 150 | 80-79-87-73-85-85 | 0.66 | 1,377 | 79% | LAG 6, DIM 100 | 78-76-83-75-**83**-82 | LAG 6, DIM 100 | EFGHIJ vs. KLABCD | Finished! |
| 22 | REJECTED (PERF) | **127** | 13 | 70% | LAG 4, DIM 200 | 66-67-76-79-70-64 | 0.25 | 886 | 65% | LAG 4, DIM 200 | 62-59-67-75-66-59 | | | Rejected due to poor performance |
| 23 | REJECTED (PERF) | **128** | 15 | 75% | LAG 6, DIM 150 | 66-84-67-83-70-83 | 0.25 | 790 | 67% | LAG 6, DIM 200 | 59-77-58-74-59-73 | | | Rejected due to poor performance |
| 24 | FINISHED 17/10/2019 | **129** | 14 | 77% | LAG 6, DIM 100 | 71-78-84-67-85-77 | 0.33 | 1,175 | 77% | LAG 6, DIM 150 | 74-78-80-69-84-**75** | LAG 6, DIM 150 | CDEFGH vs. IJKLAB | Finished! |
| 25 | FINISHED 08/10/2019 | **130** | 15 | 75% | LAG 6, DIM 200 | 70-81-68-80-70-79 | 0.50 | 1,067 | 74% | LAG 6, DIM 150 | 72-**79**-70-78-70-78 | LAG 6, DIM 150 | BCDEFG vs. HIJKLA | Finished! |

**LEGEND**

| | | |
|---|---|---|
| STATUS | Finished | Finished full experiment on listed date |
| | Rejected (Perf) | Did not meet performance criteria on localizer scans |
| | Rejected (MRI) | Revealed to be incompatible with MRI scan safety criteria after participating in behavior pre-test, but before first localizer scan |
| | Dropped (N) | Withdrew participation in the study after N sessions |
| PARTICIPANT ID | Unique identifier assigned to each participant recruited for the study between May 2018 and October 2019 | |
| LOCALIZER RUNS | 12–15 | Number of fMRI localizer runs participant underwent during first two scans (11m37s per run) |
| LO FREESURFER ROI | Avg Accuracy | Average decoding accuracy across all 6 partitions of the category space (LORO cross-validation across LOCALIZER RUNS) |
| | Best Params | Combination of parameters (PC dimensions x hemodynamic lag) that yielded the highest Avg Accuracy |
| | Accuracy % Per Partitior | Decoding accuracy for each of the 6 partitions: ABCDEF vs. GHIJKL, BCDEFG vs. HIJKLA, etc. |
| SEARCHLIGHT COGNITIVE MAP ROI | R Threshold | Cognitive map ROI represents all voxels that correlated with ideal RSA at r>R (see Methods for full description of how R was determined) |
| | # Voxels | Number of voxels in neurofeedback ROI |
| | Avg Accuracy | Average decoding accuracy across all 6 partitions of the category space (LORO cross-validation across LOCALIZER RUNS) |
| | Best Params | Combination of parameters (PC dimensions x hemodynamic lag) that yielded the highest Avg Accuracy |
| | Accuracy % Per Partitior | Decoding accuracy for each of the 6 partitions, bolded value indicates partition chosen for neurofeedback training (see Methods for description of how the partition was chosen) |
| TRAINED | Params | Parameters used for building final neurofeedback training neural model in neurofeedback ROI (identical to SEARCHLIGHT COGNITIVE MAP ROI -> Best Params) |
| | Partition | Shorthand description of which half of the shape space circle corresponds to the partition used for neurofeedback training, e.g., shapes closest to ABCDEF represent category 1, while shapes closest to GHIJKL represent category 2 |
| NOTES | Additional details about each participant who was rejected and/or withdrew their participation from the study | |

**Table S1. Participant fMRI run details, model decoding accuracy, exclusion criteria, neurofeedback ROI statistics, and training parameters.**

We recruited 25 participants for the study, of which 6 withdrew before completing the study, 2 were rejected due to not meeting MRI scanning safety criteria, 7 were rejected due to poor performance during the localizer scans, and 10 finished the full experiment. The embedded table legend provides detailed explanations for each column and field.

| PROCEDURE | | | | | |
|---|---|---|---|---|---|
| IF FIRST NEUROFEEDBACK TRAINING RUN OF SESSION 1 | | | | THEN | THRESHOLD = 70% |
| ELSE IF FIRST NEUROFEEDBACK TRAINING RUN OF DAY N | | | | | THRESHOLD = LAST THRESHOLD FROM DAY N-1 |
| ELSE IF PROGRESS WAS | 0-1 | FOR THE PREVIOUS | 1 RUN | | DECREASE THRESHOLD BY 5% (MIN. 60%) |
| | 2-3 | | 3 RUNS | | |
| | 4-5 | | 5 RUNS | | |
| | 6 | | ANY RUNS | | KEEP THRESHOLD UNCHANGED |
| | 7-8 | | 5 RUNS | | INCREASE THRESHOLD BY 5% (MAX. 90%) |
| | 9-10 | | 3 RUNS | | |
| | 11+ | | 1 RUN | | |

**Table S2. Adaptive procedure for setting the neurofeedback threshold during training.**

To account for high variability of fMRI data across multiple days and runs, the LLR feedback threshold for each training run was adjusted given participant performance (how many trials generated positive feedback) on all previous runs since the beginning of the training day using an adaptive procedure designed to converge on giving feedback for approximately 1/3 of the twenty trials in each run.

| | PARTICIPANT ID | SESSION | RUN | LLR THRESHOLD | PROGRESS |
|---|---|---|---|---|---|
| 1 | 107 NUM TRIALS: 520 | #1 25/06/2018 | 1 | 70% | 0 |
| | | | 2 | 65% | 7 |
| | | | 3 | 65% | 4 |
| | | | 4 | 65% | 4 |
| | | | 5 | 65% | 7 |
| | | | 6 | 65% | 6 |
| | | #2 26/06/2018 | 1 | 65% | 2 |
| | | #3 27/06/2018 | 1 | 65% | 3 |
| | | | 2 | 65% | 2 |
| | | | 3 | 65% | 1 |
| | | | 4 | 60% | 5 |
| | | | 5 | 60% | 9 |
| | | | 6 | 60% | 9 |
| | | | 7 | 60% | 8 |
| | | #4 28/06/2018 | 1 | 60% | 5 |
| | | | 2 | 60% | 7 |
| | | | 3 | 60% | 11 |
| | | | 4 | 65% | 5 |
| | | | 5 | 65% | 3 |
| | | #5 29/06/2018 | 1 | 65% | 3 |
| | | | 2 | 65% | 1 |
| | | | 3 | 60% | 5 |
| | | | 4 | 60% | 9 |
| | | | 5 | 60% | 8 |
| | | | 6 | 60% | 8 |
| | | | 7 | 60% | 7 |
| | | #6 N/A | | | |
| 2 | 110 NUM TRIALS: 720 | #1 10/10/2018 | 1 | 70% | 6 |
| | | | 2 | 70% | 7 |
| | | | 3 | 70% | 2 |
| | | | 4 | 70% | 6 |
| | | | 5 | 70% | 11 |
| | | #2 11/10/2018 | 1 | 70% | 8 |
| | | | 2 | 70% | 7 |
| | | | 3 | 70% | 9 |
| | | | 4 | 70% | 6 |
| | | | 5 | 70% | 9 |
| | | #3 12/10/2018 | 1 | 70% | 10 |
| | | | 2 | 70% | 6 |
| | | | 3 | 70% | 4 |
| | | | 4 | 70% | 1 |
| | | | 5 | 65% | 0 |
| | | | 6 | 60% | 7 |
| | | #4 14/10/2018 | 1 | 60% | 12 |
| | | | 2 | 65% | 11 |
| | | | 3 | 70% | 8 |
| | | | 4 | 70% | 12 |
| | | | 5 | 75% | 3 |
| | | | 6 | 75% | 6 |
| | | | 7 | 75% | 0 |
| | | #5 15/10/2018 | 1 | 75% | 6 |
| | | | 2 | 75% | 5 |
| | | | 3 | 75% | 4 |
| | | | 4 | 75% | 2 |
| | | | 5 | 75% | 4 |
| | | | 6 | 75% | 7 |
| | | #6 16/10/2018 | 1 | 75% | 5 |
| | | | 2 | 75% | 3 |
| | | | 3 | 75% | 2 |
| | | | 4 | 75% | 1 |
| | | | 5 | 70% | 6 |
| | | | 6 | 70% | 10 |
| | | | 7 | 70% | 6 |
| 3 | 117 NUM TRIALS: 560 | #1 21/03/2019 | 1 | 70% | 9 |
| | | | 2 | 70% | 9 |
| | | | 3 | 70% | 6 |
| | | | 4 | 70% | 5 |
| | | | 5 | 70% | 9 |
| | | | 6 | 70% | 6 |
| | | #2 22/03/2019 | 1 | 70% | 3 |
| | | | 2 | 70% | 6 |
| | | | 3 | 70% | 1 |
| | | | 4 | 65% | 10 |
| | | | 5 | 65% | 8 |
| | | | 6 | 65% | 12 |
| | | | 7 | 70% | 2 |
| | | #3 23/03/2019 | 1 | 70% | 4 |
| | | | 2 | 70% | 5 |
| | | | 3 | 70% | 5 |
| | | | 4 | 70% | 5 |
| | | | 5 | 70% | 3 |
| | | | 6 | 65% | 9 |
| | | #4 24/03/2019 | 1 | 65% | 14 |
| | | | 2 | 70% | 6 |
| | | | 3 | 70% | 6 |
| | | | 4 | 70% | 8 |
| | | | 5 | 70% | 4 |
| | | | 6 | 70% | 2 |
| | | | 7 | 70% | 3 |
| | | #5 25/03/2019 | 1 | 70% | 8 |
| | | | 2 | 70% | 4 |
| | | #6 N/A | | | |
| 4 | 118 NUM TRIALS: 760 | #1 23/04/2019 | 1 | 70% | 4 |
| | | | 2 | 70% | 6 |
| | | | 3 | 70% | 2 |
| | | | 4 | 70% | 8 |
| | | | 5 | 70% | 4 |
| | | | 6 | 70% | 6 |
| | | #2 24/04/2019 | 1 | 70% | 2 |
| | | | 2 | 70% | 3 |
| | | | 3 | 70% | 3 |
| | | | 4 | 65% | 8 |
| | | | 5 | 65% | 7 |
| | | | 6 | 65% | 11 |
| | | #3 25/04/2019 | 1 | 65% | 7 |
| | | | 2 | 65% | 8 |
| | | | 3 | 65% | 6 |
| | | | 4 | 65% | 7 |
| | | | 5 | 65% | 9 |
| | | | 6 | 65% | 6 |
| | | #4 26/04/2019 | 1 | 65% | 8 |
| | | | 2 | 65% | 5 |
| | | | 3 | 65% | 4 |
| | | | 4 | 65% | 8 |
| | | | 5 | 65% | 9 |
| | | | 6 | 65% | 11 |
| | | | 7 | 70% | 2 |
| | | #5 27/04/2019 | 1 | 70% | 5 |
| | | | 2 | 70% | 11 |
| | | | 3 | 75% | 2 |
| | | | 4 | 75% | 3 |
| | | | 5 | 75% | 2 |
| | | | 6 | 70% | 9 |
| | | | 7 | 70% | 6 |
| | | #6 29/04/2019 | 1 | 70% | 7 |
| | | | 2 | 70% | 5 |
| | | | 3 | 70% | 5 |
| | | | 4 | 70% | 8 |
| | | | 5 | 70% | 10 |
| | | | 6 | 70% | 7 |
| 5 | 119 NUM TRIALS: 740 | #1 21/06/2019 | 1 | 70% | 5 |
| | | | 2 | 70% | 5 |
| | | | 3 | 70% | 8 |
| | | #2 22/06/2019 | 1 | 70% | 6 |
| | | | 2 | 70% | 8 |
| | | | 3 | 70% | 6 |
| | | | 4 | 70% | 7 |
| | | | 5 | 70% | 13 |
| | | | 6 | 75% | 1 |
| | | | 7 | 70% | 7 |
| | | #3 23/06/2019 | 1 | 70% | 6 |
| | | | 2 | 70% | 7 |
| | | | 3 | 70% | 7 |
| | | | 4 | 70% | 7 |
| | | | 5 | 70% | 10 |
| | | | 6 | 70% | 9 |
| | | | 7 | 75% | 4 |
| | | #4 24/06/2019 | 1 | 75% | 2 |
| | | | 2 | 75% | 2 |
| | | | 3 | 75% | 4 |
| | | | 4 | 75% | 7 |
| | | | 5 | 75% | 3 |
| | | | 6 | 75% | 5 |
| | | | 7 | 75% | 1 |
| | | #5 25/06/2019 | 1 | 75% | 2 |
| | | | 2 | 75% | 3 |
| | | | 3 | 75% | 8 |
| | | | 4 | 75% | 5 |
| | | | 5 | 75% | 5 |
| | | | 6 | 75% | 11 |
| | | | 7 | 80% | 6 |
| | | #6 26/06/2019 | 1 | 80% | 4 |
| | | | 2 | 80% | 0 |
| | | | 3 | 75% | 3 |
| | | | 4 | 75% | 5 |
| | | | 5 | 75% | 9 |
| | | | 6 | 75% | 6 |
| 6 | 121 NUM TRIALS: 760 | #1 14/07/2019 | 1 | 70% | 4 |
| | | | 2 | 70% | 3 |
| | | | 3 | 70% | 1 |
| | | | 4 | 65% | 4 |
| | | | 5 | 65% | 3 |
| | | | 6 | 65% | 9 |
| | | #2 15/07/2019 | 1 | 65% | 10 |
| | | | 3 | 65% | 2 |
| | | | 4 | 65% | 8 |
| | | | 5 | 65% | 3 |
| | | | 6 | 65% | 6 |
| | | | 7 | 65% | 4 |
| | | #3 16/07/2019 | 1 | 65% | 9 |
| | | | 2 | 65% | 9 |
| | | | 3 | 65% | 9 |
| | | | 4 | 70% | 9 |
| | | | 5 | 75% | 4 |
| | | | 6 | 75% | 1 |
| | | #4 17/07/2019 | 1 | 75% | 0 |
| | | | 2 | 70% | 4 |
| | | | 3 | 70% | 2 |
| | | | 5 | 70% | 1 |
| | | | 6 | 65% | 4 |
| | | | 7 | 65% | 0 |
| | | #5 18/07/2019 | 1 | 65% | 2 |
| | | | 2 | 65% | 6 |
| | | | 3 | 65% | 6 |
| | | | 4 | 65% | 7 |
| | | | 5 | 65% | 4 |
| | | | 6 | 65% | 7 |
| | | | 7 | 65% | 4 |
| | | #6 19/07/2019 | 1 | 65% | 6 |
| | | | 2 | 65% | 4 |
| | | | 3 | 65% | 7 |
| | | | 4 | 65% | 6 |
| | | | 5 | 65% | 6 |
| | | | 6 | 65% | 4 |
| | | | 7 | 65% | 9 |
| 7 | 124 NUM TRIALS: 800 | #1 23/08/2019 | 1 | 70% | 6 |
| | | | 2 | 70% | 5 |
| | | | 3 | 70% | 4 |
| | | | 4 | 70% | 9 |
| | | | 5 | 70% | 7 |
| | | | 6 | 70% | 8 |
| | | | 7 | 70% | 5 |
| | | #2 24/08/2019 | 1 | 70% | 9 |
| | | | 2 | 70% | 10 |
| | | | 3 | 70% | 5 |
| | | | 4 | 70% | 5 |
| | | | 5 | 70% | 4 |
| | | | 6 | 70% | 6 |
| | | | 7 | 70% | 6 |
| | | #3 25/08/2019 | 1 | 70% | 5 |
| | | | 2 | 70% | 5 |
| | | | 3 | 70% | 5 |
| | | | 4 | 70% | 10 |
| | | | 5 | 70% | 8 |
| | | #4 26/08/2019 | 1 | 70% | 4 |
| | | | 2 | 70% | 2 |
| | | | 3 | 70% | 2 |
| | | | 4 | 70% | 4 |
| | | | 5 | 70% | 6 |
| | | | 6 | 70% | 7 |
| | | | 7 | 70% | 1 |
| | | #5 27/08/2019 | 1 | 70% | 9 |
| | | | 2 | 70% | 7 |
| | | | 3 | 70% | 11 |
| | | | 4 | 75% | 3 |
| | | | 5 | 75% | 1 |
| | | | 6 | 70% | 6 |
| | | | 7 | 70% | 5 |
| | | #6 28/08/2019 | 1 | 70% | 7 |
| | | | 2 | 70% | 4 |
| | | | 3 | 70% | 3 |
| | | | 4 | 70% | 5 |
| | | | 5 | 70% | 10 |
| | | | 6 | 70% | 6 |
| | | | 7 | 70% | 2 |
| 8 | 126 NUM TRIALS: 760 | #1 28/08/2019 | 1 | 70% | 0 |
| | | | 2 | 65% | 2 |
| | | | 3 | 65% | 3 |
| | | | 5 | 65% | 2 |
| | | | 6 | 60% | 6 |
| | | | 7 | 60% | 9 |
| | | #2 29/08/2019 | 1 | 60% | 6 |
| | | | 2 | 60% | 5 |
| | | | 3 | 60% | 2 |
| | | | 4 | 60% | 4 |
| | | | 5 | 60% | 3 |
| | | | 6 | 60% | 4 |
| | | | 7 | 60% | 4 |
| | | #3 30/08/2019 | 1 | 60% | 9 |
| | | | 2 | 60% | 12 |
| | | | 3 | 65% | 10 |
| | | | 5 | 70% | 10 |
| | | | 6 | 75% | 9 |
| | | | 7 | 80% | 7 |
| | | #4 31/08/2019 | 1 | 80% | 0 |
| | | | 2 | 75% | 8 |
| | | | 3 | 75% | 1 |
| | | | 4 | 70% | 8 |
| | | | 5 | 70% | 9 |
| | | | 6 | 70% | 0 |
| | | | 7 | 65% | 5 |
| | | #5 02/09/2019 | 1 | 65% | 10 |
| | | | 2 | 65% | 10 |
| | | | 3 | 65% | 11 |
| | | | 4 | 70% | 1 |
| | | | 5 | 65% | 10 |
| | | #6 03/09/2019 | 1 | 65% | 9 |
| | | | 2 | 65% | 7 |
| | | | 3 | 65% | 4 |
| | | | 4 | 65% | 4 |
| | | | 5 | 65% | 10 |
| | | | 6 | 65% | 3 |
| | | | 7 | 65% | 4 |
| 9 | 130 NUM TRIALS: 740 | #1 02/10/2019 | 1 | 70% | 13 |
| | | | 2 | 75% | 7 |
| | | | 3 | 75% | 6 |
| | | | 4 | 75% | 7 |
| | | | 5 | 75% | 8 |
| | | | 6 | 75% | 7 |
| | | #2 03/10/2019 | 2 | 75% | 7 |
| | | | 4 | 75% | 3 |
| | | | 5 | 75% | 5 |
| | | #3 04/10/2019 | 1 | 75% | 9 |
| | | | 2 | 75% | 7 |
| | | | 3 | 75% | 6 |
| | | | 4 | 75% | 10 |
| | | | 5 | 75% | 10 |
| | | | 6 | 75% | 10 |
| | | | 7 | 80% | 5 |
| | | #4 05/10/2019 | 1 | 80% | 1 |
| | | | 2 | 75% | 9 |
| | | | 3 | 75% | 5 |
| | | | 4 | 75% | 6 |
| | | | 5 | 75% | 0 |
| | | | 6 | 70% | 11 |
| | | | 7 | 75% | 8 |
| | | #5 06/10/2019 | 1 | 75% | 7 |
| | | | 2 | 75% | 1 |
| | | | 3 | 70% | 12 |
| | | | 4 | 75% | 8 |
| | | | 5 | 75% | 8 |
| | | | 6 | 75% | 5 |
| | | | 7 | 75% | 1 |
| | | #6 07/10/2019 | 1 | 75% | 9 |
| | | | 2 | 75% | 9 |
| | | | 3 | 75% | 4 |
| | | | 4 | 75% | 4 |
| | | | 5 | 75% | 4 |
| | | | 6 | 75% | 2 |
| | | | 7 | 75% | 2 |
| 10 | 129 NUM TRIALS: 800 | #1 11/10/2019 | 1 | 70% | 5 |
| | | | 2 | 70% | 0 |
| | | | 3 | 65% | 6 |
| | | | 4 | 65% | 5 |
| | | | 5 | 65% | 5 |
| | | #2 12/10/2019 | 1 | 65% | 6 |
| | | | 2 | 65% | 7 |
| | | | 3 | 65% | 5 |
| | | | 4 | 65% | 13 |
| | | | 5 | 70% | 5 |
| | | | 6 | 70% | 4 |
| | | | 7 | 70% | 3 |
| | | #3 13/10/2019 | 1 | 70% | 11 |
| | | | 2 | 75% | 7 |
| | | | 3 | 75% | 4 |
| | | | 4 | 75% | 3 |
| | | | 5 | 75% | 7 |
| | | | 6 | 75% | 5 |
| | | | 7 | 75% | 4 |
| | | #4 14/10/2019 | 1 | 75% | 7 |
| | | | 2 | 75% | 6 |
| | | | 3 | 75% | 9 |
| | | | 4 | 75% | 7 |
| | | | 5 | 75% | 9 |
| | | | 6 | 75% | 9 |
| | | | 7 | 75% | 0 |
| | | #5 15/10/2019 | 1 | 75% | 1 |
| | | | 2 | 70% | 9 |
| | | | 3 | 70% | 5 |
| | | | 4 | 70% | 6 |
| | | | 5 | 70% | 5 |
| | | | 6 | 70% | 5 |
| | | | 7 | 70% | 1 |
| | | #6 16/10/2019 | 1 | 70% | 11 |
| | | | 2 | 75% | 3 |
| | | | 3 | 75% | 4 |
| | | | 4 | 75% | 1 |
| | | | 5 | 70% | 3 |
| | | | 6 | 70% | 5 |
| | | | 7 | 70% | 8 |

| LEGEND | |
|---|---|
| PARTICIPANT ID | Unique identifier assigned to each participant recruited for the study between May 2018 and October 2019, includes total number of neurofeedback training trials for this participant (520--800) |
| SESSION | Date of number of neurofeedback training session |
| RUN | Number of functional neurofeedback training run for current session |
| LLR THRESHOLD | Minimum log-likelihood ratio used to give positive feedback on current RUN (see Methods and Table S4 for details about how this was set) |
| PROGRESS | Number of trials in current RUN where positive feedback was given to participant (min. 0, max. 20) |

**Table S3. Neurofeedback training run details.**

We report details on every feedback run for every training session of every participant, including number of trials with positive feedback and corresponding feedback LLR threshold for each run computed using the procedure detailed in Table S2.

| LLR COMPUTATION | Main Effect: TRAINED - CONTROL | | | | BEHAVIOR CORRELATION | |
|---|---|---|---|---|---|---|
| | mean | s.e.m. | t | p | r | p |
| TR 1 (main result) | 0.369 | 0.154 | 2.27 | 0.049 | 0.761 | 0.011 |
| TRs 1-2 | 0.400 | 0.154 | 2.46 | 0.036 | 0.654 | 0.040 |
| TRs 1-3 | 0.335 | 0.137 | 2.32 | 0.045 | 0.603 | 0.065 |
| TRs 1-4 | 0.268 | 0.108 | 2.35 | 0.044 | 0.562 | 0.091 |
| TRs 1-5 | 0.191 | 0.079 | 2.30 | 0.047 | 0.524 | 0.120 |

**Table S4. Effects of neurofeedback training are similar when using LLR measured from any number of TRs from each trial, compared to using the first TR only.**

LLR difference score computed using model estimates from a variable number of TRs across each neurofeedback trial (1-5; participants received at most 5 instances of feedback per trial). We observed similar results in all cases to when using only the first TR from each trial, with minor weakening of the reported main effect of neurofeedback training as more TRs were added, likely due to fMRI adaptation effects (Grill-Spector & Malach, 2001; Aguirre, 2007) across the length of the 16s trials.

| | PARTICIPANT ID | PRE-TEST DATE | PRE-TEST AG | PRE-TEST BH | PRE-TEST CI | PRE-TEST DJ | PRE-TEST EK | PRE-TEST FL | POST-TEST DATE | POST-TEST AG | POST-TEST BH | POST-TEST CI | POST-TEST DJ | POST-TEST EK | POST-TEST FL | TRAINED CATEGORIES |
|---|---|---|---|---|---|---|---|---|---|---|---|---|---|---|---|---|
| 1 | **107** | 18/06/2018 | 2.479 | 1.519 | 2.261 | 3.027 | 2.236 | 2.185 | 30/06/2018 | 3.251 | 2.442 | 1.956 | 3.045 | 2.078 | 3.174 | EFGHIJ vs. KLABCD |
| 2 | **110** | 20/07/2018 | 1.589 | 1.676 | 1.482 | 2.363 | 1.734 | 1.317 | 17/10/2018 | 1.793 | 1.908 | 2.233 | 2.158 | 2.164 | 2.069 | FGHIJK vs. LABCDE |
| 3 | **117** | 26/02/2019 | 1.771 | 2.070 | 1.551 | 1.650 | 2.267 | 1.284 | 26/03/2019 | 1.073 | 1.136 | 1.018 | 1.105 | 1.144 | 0.994 | CDEFGH vs. IJKLAB |
| 4 | **118** | 28/03/2019 | 2.682 | 2.527 | 2.037 | 1.926 | 1.390 | 1.299 | 30/04/2019 | 0.282 | 0.531 | 0.540 | 0.317 | 0.608 | 0.501 | BCDEFG vs. HIJKLA |
| 5 | **119** | 12/06/2019 | 2.114 | 1.173 | 2.702 | 1.716 | 2.823 | 1.982 | 27/06/2019 | 1.047 | 2.748 | 2.587 | 2.011 | 1.702 | 1.528 | ABCDEF vs. GHIJKL |
| 6 | **121** | 24/06/2019 | 2.083 | 2.404 | 1.360 | 1.538 | 1.361 | 1.542 | 20/07/2019 | 2.945 | 4.300 | 1.513 | 1.735 | 2.447 | 11.856 | DEFGHI vs. JKLABC |
| 7 | **124** | 13/08/2019 | 1.970 | 1.566 | 2.535 | 1.450 | 2.685 | 1.941 | 29/08/2019 | 2.347 | 2.713 | 2.703 | 2.630 | 2.552 | 3.024 | DEFGHI vs. JKLABC |
| 8 | **126** | 21/08/2019 | 1.370 | 2.039 | 2.124 | 1.723 | 2.227 | 1.783 | 04/09/2019 | 1.407 | 2.128 | 2.009 | 1.248 | 1.132 | 1.842 | EFGHIJ vs. KLABCD |
| 9 | **130** | 19/09/2019 | 1.703 | 1.647 | 1.708 | 1.994 | 0.941 | 1.363 | 08/10/2019 | 1.315 | 1.608 | 1.389 | 0.990 | 1.086 | 1.655 | BCDEFG vs. HIJKLA |
| 10 | **129** | 19/09/2019 | 1.414 | 1.753 | 1.491 | 1.302 | 1.876 | 1.511 | 17/10/2019 | 1.637 | 1.479 | 1.876 | 1.237 | 2.500 | 2.089 | CDEFGH vs. IJKLAB |

| | PARTICIPANT ID | PSYCHOMETRIC FUNCTION SLOPE SHARPENING: (POST-PRE/(POST+PRE) AG | BH | CI | DJ | EK | FL | TRAINED | NEUTRAL | CONTROL | EFFECT (T-U) |
|---|---|---|---|---|---|---|---|---|---|---|---|
| 1 | **107** | 0.135 | 0.233 | -0.072 | 0.003 | -0.037 | 0.185 | 0.184 | 0.056 | -0.017 | 0.201 |
| 2 | **110** | 0.060 | 0.065 | 0.202 | -0.045 | 0.110 | 0.222 | 0.133 | 0.007 | 0.166 | -0.033 |
| 3 | **117** | -0.245 | -0.291 | -0.207 | -0.198 | -0.329 | -0.127 | -0.228 | -0.222 | -0.249 | 0.021 |
| 4 | **118** | -0.810 | -0.653 | -0.581 | -0.717 | -0.391 | -0.443 | -0.554 | -0.512 | -0.731 | 0.177 |
| 5 | **119** | -0.338 | 0.402 | -0.022 | 0.079 | -0.248 | -0.129 | 0.029 | 0.077 | -0.233 | 0.262 |
| 6 | **121** | 0.171 | 0.283 | 0.053 | 0.060 | 0.285 | 0.770 | 0.471 | 0.284 | 0.057 | 0.414 |
| 7 | **124** | 0.087 | 0.268 | 0.032 | 0.289 | -0.025 | 0.218 | 0.153 | 0.121 | 0.161 | -0.008 |
| 8 | **126** | 0.013 | 0.021 | -0.028 | -0.160 | -0.326 | 0.016 | 0.017 | -0.006 | -0.243 | 0.260 |
| 9 | **130** | -0.129 | -0.012 | -0.103 | -0.336 | 0.072 | 0.097 | -0.132 | -0.003 | -0.070 | -0.062 |
| 10 | **129** | 0.073 | -0.085 | 0.114 | -0.026 | 0.143 | 0.161 | 0.152 | 0.024 | 0.015 | 0.137 |
| | AVERAGE EFFECT | | | | | | | 0.022 | -0.017 | -0.115 | 0.137 |

LEGEND

Trained Directions Psychometric Function Slope
Neutral Directions Psychometric Function Slope
Untrained Directions Psychometric Function Slope

| | | |
|---|---|---|
| PARTICIPANT ID | | Unique identifier assigned to each participant recruited for the study between May 2018 and October 2019 |
| PRE-TEST | Date | Date of behavioral pre-test |
| | AG | Psychometric function slope estimate in behavioral pre-test for AG direction |
| | BH | Psychometric function slope estimate in behavioral pre-test for BH direction |
| | CI | Psychometric function slope estimate in behavioral pre-test for CI direction |
| | DJ | Psychometric function slope estimate in behavioral pre-test for DJ direction |
| | EK | Psychometric function slope estimate in behavioral pre-test for EK direction |
| | FL | Psychometric function slope estimate in behavioral pre-test for FL direction |
| POST-TEST | Date | Date of behavioral post-test |
| | AG | Psychometric function slope estimate in behavioral post-test for AG direction |
| | BH | Psychometric function slope estimate in behavioral post-test for BH direction |
| | CI | Psychometric function slope estimate in behavioral post-test for CI direction |
| | DJ | Psychometric function slope estimate in behavioral post-test for DJ direction |
| | EK | Psychometric function slope estimate in behavioral post-test for EK direction |
| | FL | Psychometric function slope estimate in behavioral post-test for FL direction |
| TRAINED CATEGORIES | | Shorthand description of which half of the shape space circle corresponds to the partition used for neurofeedback training, e.g., shapes closest to ABCDEF represent category 1, while shapes closest to GHIJKL represent category 2 |
| SLOPE SHARPENING | AG | Psychometric function slope sharpening between pre- and post-test: AG direction (normalized difference score) |
| | BH | Psychometric function slope sharpening between pre- and post-test: BH direction (normalized difference score) |
| | CI | Psychometric function slope sharpening between pre- and post-test: CI direction (normalized difference score) |
| | DJ | Psychometric function slope sharpening between pre- and post-test: DJ direction (normalized difference score) |
| | EK | Psychometric function slope sharpening between pre- and post-test: EK direction (normalized difference score) |
| | FL | Psychometric function slope sharpening between pre- and post-test: FL direction (normalized difference score) |
| | Trained | Average sharpening for two directions at 75 degrees and 105 degrees to category boundary |
| | Neutral | Average sharpening for two directions at 45 degrees and 135 degrees to category boundary |
| | Control | Average sharpening for two directions at 15 degrees and 165 degrees to category boundary |
| | Effect | Trained - Control |

**Table S5. Estimated psychometric function slopes and changes between pre- (day 1) and post-tests (day 10), for all 6 directions in shape space and 10 participants.**

The embedded table legend provides detailed explanations for each column and field.

| | PARTICIPANT ID | PRE-TEST | | | | | | | POST-TEST | | | | | | |
|---|---|---|---|---|---|---|---|---|---|---|---|---|---|---|---|
| | | DATE | AG | BH | CI | DJ | EK | FL | DATE | AG | BH | CI | DJ | EK | FL |
| 1 | **107** | 18/06/2018 | 0.672 | 0.378 | 0.628 | 0.767 | 0.612 | 0.608 | 30/06/2018 | 0.674 | 0.622 | 0.549 | 0.711 | 0.564 | 0.808 |
| 2 | **110** | 20/07/2018 | 0.448 | 0.407 | 0.425 | 0.643 | 0.451 | 0.346 | 17/10/2018 | 0.546 | 0.497 | 0.621 | 0.645 | 0.604 | 0.573 |
| 3 | **117** | 26/02/2019 | 0.479 | 0.517 | 0.411 | 0.450 | 0.443 | 0.331 | 26/03/2019 | 0.338 | 0.320 | 0.324 | 0.330 | 0.338 | 0.328 |
| 4 | **118** | 28/03/2019 | 0.710 | 0.724 | 0.591 | 0.521 | 0.381 | 0.354 | 30/04/2019 | 0.078 | 0.457 | 0.131 | 0.090 | 0.183 | 0.145 |
| 5 | **119** | 12/06/2019 | 0.564 | 0.333 | 0.728 | 0.459 | 0.781 | 0.496 | 27/06/2019 | 0.274 | 0.733 | 0.711 | 0.555 | 0.348 | 0.421 |
| 6 | **121** | 24/06/2019 | 0.531 | 0.597 | 0.333 | 0.387 | 0.336 | 0.397 | 20/07/2019 | 0.641 | 0.803 | 0.433 | 0.498 | 0.635 | 1.224 |
| 7 | **124** | 13/08/2019 | 0.536 | 0.380 | 0.569 | 0.403 | 0.681 | 0.536 | 29/08/2019 | 0.671 | 0.654 | 0.741 | 0.717 | 0.654 | 0.741 |
| 8 | **126** | 21/08/2019 | 0.319 | 0.562 | 0.581 | 0.425 | 0.584 | 0.425 | 04/09/2019 | 0.386 | 0.562 | 0.460 | 0.331 | 0.291 | 0.515 |
| 9 | **130** | 19/09/2019 | 0.426 | 0.478 | 0.492 | 0.488 | 0.248 | 0.310 | 08/10/2019 | 0.373 | 0.465 | 0.380 | 0.261 | 0.289 | 0.433 |
| 10 | **129** | 19/09/2019 | 0.423 | 0.504 | 0.435 | 0.322 | 0.517 | 0.390 | 17/10/2019 | 0.444 | 0.379 | 0.507 | 0.322 | 0.701 | 0.537 |

| | TRAINED CATEGORIES | PSYCHOMETRIC FUNCTION SLOPE SHARPENING: POST-PRE | | | | | | | | | |
|---|---|---|---|---|---|---|---|---|---|---|---|
| | | AG | BH | CI | DJ | EK | FL | TRAINED | NEUTRAL | CONTROL | EFFECT (T-C) |
| 1 | EFGHIJ vs. KLABCD | 0.002 | 0.244 | -0.078 | -0.056 | -0.048 | 0.200 | 0.123 | 0.061 | -0.052 | 0.175 |
| 2 | FGHIJK vs. LABCDE | 0.098 | 0.090 | 0.196 | 0.002 | 0.153 | 0.227 | 0.143 | 0.050 | 0.190 | -0.047 |
| 3 | CDEFGH vs. IJKLAB | -0.141 | -0.197 | -0.088 | -0.120 | -0.105 | -0.003 | -0.054 | -0.130 | -0.142 | 0.088 |
| 4 | BCDEFG vs. HIJKLA | -0.632 | -0.267 | -0.460 | -0.431 | -0.199 | -0.209 | -0.315 | -0.334 | -0.449 | 0.135 |
| 5 | ABCDEF vs. GHIJKL | -0.290 | 0.399 | -0.018 | 0.096 | -0.433 | -0.075 | 0.039 | -0.017 | -0.182 | 0.221 |
| 6 | DEFGHI vs. JKLABC | 0.110 | 0.206 | 0.100 | 0.111 | 0.299 | 0.827 | 0.469 | 0.253 | 0.106 | 0.363 |
| 7 | DEFGHI vs. JKLABC | 0.135 | 0.274 | 0.172 | 0.314 | -0.027 | 0.206 | 0.170 | 0.124 | 0.243 | -0.072 |
| 8 | EFGHIJ vs. KLABCD | 0.066 | 0.001 | -0.122 | -0.094 | -0.293 | 0.090 | 0.034 | -0.016 | -0.193 | 0.227 |
| 9 | BCDEFG vs. HIJKLA | -0.053 | -0.013 | -0.112 | -0.227 | 0.041 | 0.123 | -0.093 | 0.006 | -0.033 | -0.060 |
| 10 | CDEFGH vs. IJKLAB | 0.021 | -0.125 | 0.072 | -0.001 | 0.184 | 0.147 | 0.165 | 0.010 | -0.026 | 0.192 |
| | | | | | | | | 0.068 | 0.000 | -0.054 | 0.122 |

**LEGEND**

Trained Directions Psychometric Function Slope
Neutral Directions Psychometric Function Slope
Control Directions Psychometric Function Slope

| | | |
|---|---|---|
| PARTICIPANT ID | | Unique identifier assigned to each participant recruited for the study between May 2018 and October 2019 |
| PRE-TEST | Date | Date of behavioral pre-test |
| | AG | Psychometric function slope estimate in behavioral pre-test for AG direction |
| | BH | Psychometric function slope estimate in behavioral pre-test for BH direction |
| | CI | Psychometric function slope estimate in behavioral pre-test for CI direction |
| | DJ | Psychometric function slope estimate in behavioral pre-test for DJ direction |
| | EK | Psychometric function slope estimate in behavioral pre-test for EK direction |
| | FL | Psychometric function slope estimate in behavioral pre-test for FL direction |
| POST-TEST | Date | Date of behavioral post-test |
| | AG | Psychometric function slope estimate in behavioral post-test for AG direction |
| | BH | Psychometric function slope estimate in behavioral post-test for BH direction |
| | CI | Psychometric function slope estimate in behavioral post-test for CI direction |
| | DJ | Psychometric function slope estimate in behavioral post-test for DJ direction |
| | EK | Psychometric function slope estimate in behavioral post-test for EK direction |
| | FL | Psychometric function slope estimate in behavioral post-test for FL direction |
| TRAINED CATEGORIES | | Shorthand description of which half of the shape space circle corresponds to the partition used for neurofeedback training, e.g., shapes closest to ABCDEF represent category 1, while shapes closest to GHIJKL represent category 2 |
| SLOPE SHARPENING | AG | Psychometric function slope sharpening between pre- and post-test: AG direction (difference score) |
| | BH | Psychometric function slope sharpening between pre- and post-test: BH direction (difference score) |
| | CI | Psychometric function slope sharpening between pre- and post-test: CI direction (difference score) |
| | DJ | Psychometric function slope sharpening between pre- and post-test: DJ direction (difference score) |
| | EK | Psychometric function slope sharpening between pre- and post-test: EK direction (difference score) |
| | FL | Psychometric function slope sharpening between pre- and post-test: FL direction (difference score) |
| | Trained | Average sharpening for two directions at 75 degrees and 105 degrees to category boundary |
| | Neutral | Average sharpening for two directions at 45 degrees and 135 degrees to category boundary |
| | Control | Average sharpening for two directions at 15 degrees and 165 degrees to category boundary |
| | Effect | Trained - Control |

**Table S6. Parameters for alternate estimates of psychometric function slopes and changes between pre- (day 1) and post-tests (day 10), for all 6 directions in shape space and 10 participants.**

To measure whether lapses in performance between the behavioral pre- (Day 1) and post-tests (Day 10) could potentially explain the effects of neurofeedback training on perception, we performed a separate joint estimate of the lapses (upper and lower), thresholds, and slopes for our psychometric functions. Corresponding psychometric functions are shown in Fig. S36.

| PARTICIPANT | QUESTION 1 |
|---|---|
| | Did you suspect that there were multiple categories of shapes during the experiment? If so, how many? On this map of the shape space, please draw the categories (you may draw any category boundaries, regardless of the diameters indicated, e.g., AG, BH, etc.). *(Author's note: boundaries drawn shown in Fig. S18).* |
| | **ANSWERS** |
| **107** | Initially, I thought there were 3 categories of shapes, but on the last day I realized they might have all been the same shape somehow. |
| **110** | Yes, maybe 4-5. |
| **117** | Yes, 3. |
| **118** | Yes, about 4 categories. *(Author's note: participant wrote 4, crossed it out, then wrote 5, crossed it out, then wrote 4 again while thinking about the question for more than minute).* |
| **119** | I sometimes suspected 2 and sometimes 3, depending on the day. |
| **121** | I identified 4 categories. |
| **124** | 3-4. |
| **126** | Yes. I thought there were two main categories of shapes, w/ each category having variants of the origin shape. |
| **130** | Yes, 2. |
| **129** | I thought there were about 4-5 categories; yes, I did suspect there were multiple categories. |

| PARTICIPANT | QUESTION 2 |
|---|---|
| | At the beginning of the experiment, we selected a random category boundary in this space by choosing a random diameter (straight line). If you had to guess, which straight line do you think separates Category 1 from Category 2? How confident are you about your choice on a scale from 1 to 5 (1 = not at all; 5 = very confident)? On this map of the shape space, please draw what you think the most likely category boundary was (you may draw any straight line, regardless of the diameters indicated, e.g., AG, BH, etc.). *(Author's note: boundaries drawn shown in Fig. S18).* |
| | **ANSWERS** |
| **107** | 4. *(Author's note: the participant drew a line on the form close to DJ, see Fig. S18).* |
| **110** | GA, 3. |
| **117** | A-J-F. 4. C-I 2. *(Author's note: the first category split refers to Question 1 above, the second category split refers to Question 2).* |
| **118** | DJ. 3. |
| **119** | A-G (3). |
| **121** | DJ. 4. |
| **124** | 3 (EK). |
| **126** | Direction: F->L. confidence: 4. |
| **130** | CI, 3. |
| **129** | DJ, confidence: 3. *(Author's note: after clarifying with the participant, the actual boundary they chose was AG = orthogoinal direction to DJ.)* |

| PARTICIPANT | QUESTION 3 |
|---|---|
| | When performing the main experimental task in the scanner (i.e., 'Generate a mental state that makes the shapes stop wobbling'), did you explicitly think about other shapes that were different from the ones on the screen? If so, to which of the A--L shapes at the exterior of the circle were they most similar to? Please share anything else you remember about the strategy or strategies you employed during the experiment. |
| | **ANSWERS** |
| **107** | I thought about the particular shape at hand and the shapes like A and D were much easier to stop. To get the shapes to altogether stop I repeatedly said "stop" in my head in a yodel kind of voice. But I never was able to stop the shapes like H. |
| **110** | No, I was mostly thinking/visualizing the shape stopping. |
| **117** | No. I imagined a static form of the shape very close to the shape and focused my attention on the boundaries. I also tried to focus on the static white background. *(Author's note: the shapes were black on an equiluminant grey background.)* |
| **118** | No, I tried to think about something that concentrated me completely. Like one shape reminded me of my mom's side profile. LOL. So I would think of her. |
| **119** | Yes, I imagined a star *(Author's note: participant drew a five-pointed star on the form mid-sentence)* , which was most similar to shapes A thru E. |
| **121** | I tried so many strategies, some ridiculously complex. I settled on recognizing & categorizing the shape -- deciding which category I thought it belonged to and focusing on that shape at rest. |
| **124** | Yes, I tried to look at the shape out of focus so that it would gradually morph ito different shapes. Sometimes this seemed to work. |
| **126** | I didn't really think about other shapes. I first quickly determined which of my two categories the shape on the screen belongs to, and from that point I picked a moving spot on the shape to focus on continuously. If the shape was in the "mounds" category, I would focus on the curve of the divet, if the shape was in the "butterfly" category, I would focus on the moving point that points into the "body/wings". |
| **130** | Shapes BCD were similar, as well as FGHI. I thought about filling up space for shapes BCD and shrinking space for FGHI. |
| **129** | No, I did not think of other shapes in order to make the current shape stop wiggling. My strategy was to repeat a word/phrase, as well as mentally visualize a corresponding scene when necessary. |

| PARTICIPANT | QUESTION 4 |
|---|---|
| | Were any of the shapes or shape types more memorable than others, i.e. did you think about certain trials as 'This is a pointy shape, I need to think about X'? Did you have explicit names for any of the shapes? Did these names evolve or change over the course of the experiment (i.e., days)? |
| | **ANSWERS** |
| **107** | I called shapes like A "rabbits", shapes like H "butterflies", and shapes like C "starfish". These names were the same over the course of the experiment. The A and C shapes were most memorable because I actually got more to stop. |
| **110** | C bug, F tree/cloud on a stick, and I little girl/Powerpuff girl. |
| **117** | Yes, shape H was distinct and felt was easiest to reduce jiggling. Shape E/D was memorable as a more difficult one to stop jiggling. |
| **118** | I had names for them. 1 girl, 2 elephant, 3 dude, 4 flower. They became easier to categorize as the days passed. |
| **119** | Yes. At first, A--D, J--L were walrus, E--H was butterfly, I--K was laughing frog. By the end, B--E was star, G--J was pacman. |
| **121** | Yes, I had names for the shape types and the names/categories changed over the runs. The central white pixel made an eye and the shapes were recognizable faces. |
| **124** | Yes. Shapes B--L reminded me of a bee, shapes E--H of a flower. No they did not change. |
| **126** | The names for my categories are mentioned above as "mounds" and "butterfly". These names came about in probably the third scanning session, when I really figured out my above mentioned techniques. |
| **130** | I remembered shape L as a rabbit, J as doll, E as clover, and C as Snoopy. The names remained the same. |
| **129** | Yes! For shapes that were more pointy in particular, I had to come up with a scene to accompany my repeated word (as these shapes were harder to stop from wiggling). I didn't have a name for shapes, but I did think that one shape looked like a pokemon. |

| PARTICIPANT | QUESTION 5 |
|---|---|
| | Did you focus on specific features of the shapes during the experiment in order to perform the task (e.g., the top pointy end, the bottom blobby end)? Did you feel that this worked, i.e. your performance improved by doing so? How confident are you about the effectiveness of this strategy on a scale from 1 to 5 (1 = useless; 5 = very effective)? |
| | **ANSWERS** |
| **107** | For the non-fmri portions I focused on curved or sharp edges and found this helpful (5). *(Author's note: the participant is referring to the two-alternative forced choice pre- and post-behavioral experiments.)* But for the scanner I viewed a shape more holistically as art of one of my categories. |
| **110** | Yes, like the wings on the bug and the curve of the girl's face. |
| **117** | I focused on a point in the shapes with the largest surface area. 3. I think this worked well for shape I. |
| **118** | Yes, I focused on specific parts when categorize them. It woked in terms of quickly categorizing a shape into i.e. girl and using a strategy that previously worked. 3. |
| **119** | Yes *(Author's note: participant drew a five-pointed star on the form mid-sentence with an arrow pointing towards the gap between two vertices)*, this part of B--E. It worked for these shapes. I never quite got it right for the other shapes. |
| **121** | I continuously made strategies about shape features since focusing on particular elements seemed to work at first. I had more success moving back and considering/focusing on the shape as a whole. I think my strategy was to see the shapes as old friends, to concentrate on recognizing them. |
| **124** | I tended to focus on parts of the shape that weren't wiggling as much or on all wiggly parts at the same time, I'd say these strategies worked the best for me. 4. |
| **126** | This is also mentioned in my answer to question 3. My "curve of the divet" seems to be your "bottom blobby end. Confidence: 5. When I was relaxed and able to focus it worked very well, but sometimes I simply got distracted or had too many thoughts going through my head for the technique to work. |
| **130** | I focused on the front (left) indent and is that didn't seem immediately clear, I look at the middle to compare the balance between the left and right. 4. |
| **129** | For shapes with less pointy edges, I focused on the center. For shapes with more pointy ends, I actually focused on the crevices between the pointy parts. I think that this strategy, after practicing, was 4.5 on a scale of effectiveness. |

| PARTICIPANT | QUESTION 6 |
|---|---|
| | Please share anything else you think might be relevant about your experience with our study. |
| | **ANSWERS** |
| **107** | It was a very interesting task that I didn't think I'd be able to do at first but was pleasantly surprised when I was able to stop shapes. |
| **110** | Some shape types were certainly harder to stop than others, the Bug (C,B,A,D,E) were all hard. |
| **117** | Visual fixation & mental distraction; I felt that whenever I was thikning about something else or visualizing something else in my head, while maintaining a solid/steady gaze was efective & less mentally taxing. |
| **118** | Having multiple days was useful. I felt it was a bit harder after the day break but not that bad. *(Author's note: this was one of the two participants who requested a 48h gap between two of the sessions for personal reasons).* |
| **119** | N/A. |
| **121** | This is so fascinating. |
| **124** | N/A. |
| **126** | Although the goal was to be totally focused, sometimes when I stopped the shape I had a single thought or scenario going through my head, almost as if that helped me stop the shape. |
| **130** | N/A. |
| **129** | N/A. |

**Table S7. Final debriefing form and summary of participant answers.**

At the conclusion of the study, but before being fully briefed on the goals of the experiment, participants were asked to complete a questionnaire about their experience, strategies for completing the task, as well as guesses (and drawings) of potential category boundaries on a never-before-seen picture of the circular stimulus space.

**Movie S1 (Separate File). Demo of oscillating shape neurofeedback trial.**

Video shows two example trials during a neurofeedback training run with no feedback (first trial, shape oscillation radius 1.875 a.d.u.) and simulated positive feedback that eventually stops the shape from oscillating (second trial, shape oscillation radii: 1.875 a.d.u., 1.250 a.d.u., 0.625 a.d.u., 0 a.d.u.).

**Movie S2 (Separate File). Detailed demo of oscillating shape neurofeedback trial.**

Video shows an example trial during a neurofeedback training run with simulated positive feedback that eventually stops the shape from oscillating (shape oscillation radii: 1.875 a.d.u., 1.250 a.d.u., 0.625 a.d.u., 0 a.d.u.). Top right corner shows simulated trajectory of current stimulus in neural shape model. Bottom left corner shows extent of shape oscillation in circular shape space.
